# Current methods integrating variant functional annotation scores have limited capacity to improve the power of genome-wide association studies

**DOI:** 10.1101/2020.11.25.396721

**Authors:** Jianhui Gao, Osvaldo Espin-Garcia, Andrew D. Paterson, Lei Sun

**Affiliations:** Division of Biostatistics, Dalla Lana School of Public Health, University of Toronto, Toronto, ON, Canada; Department of Biostatistics, Princess Margaret Cancer Centre, University Health Network, Toronto, ON, Canada; Program in Genetics and Genome Biology, The Hospital for Sick Children, Toronto, ON, Canada; Department of Statistical Sciences, Faculty of Arts and Science, University of Toronto, Toronto, ON, Canada

## Abstract

Functional annotations have the potential to increase the power of genome-wide association studies (GWAS) by prioritizing variants according to their biological function. Focusing on variant-specific annotation meta-scores including CADD (Kircher et al., 2014) and Eigen (Ionita-laza et al., 2016), we broadly examined GWAS summary statistics of 1,132 traits from the UK Biobank (Sudlow et al., 2015) using the weighted p-value approach (Genovese et al., 2006) and stratified false discovery control (sFDR) method (Sun et al., 2006). These 1,132 traits were rated by Benjamin Neale’s lab from the Broad Institute as having medium to high confidence for their heritability estimates.

Averaged across the 1,132 UK Biobank traits, sFDR was more robust to uninformative meta-scores, but the weighted p-value method identified more variants using CADD or Eigen, based on performance measures that included type I error control, recall, precision, and relative efficiency. Our application results were consistent with those from an extensive simulation study using three different designs, including leveraging the real genetic data combined with simulated genomic data and vice versa.

We also considered the recent FINDOR method (Kichaev et al., 2019), which leverages a set of individual 75 functional annotations into GWAS. An earlier application of FINDOR to 27 traits selected from the z7 category (SNP-heritability p-value < 1.27 × 10^−12^ by Nealelab) detected 13%-20% additional genome-wide significant loci as compared to the standard annotation-free GWAS, which we confirmed. Moreover, across all 438 traits in the z7 category, 46,631 out 59,764 (80%) significant loci discovered are common across the three data-integration methods.

However, across all the 1,132 UK Biobank traits examined, the median [Q1,Q3] of the total numbers of new, genome-wide significant independent loci were 0 [0, 3] by FINDOR, 0 [0, 2] by weighted p-value, and 0 [0, 0] by sFDR. Notably, 162 traits (89%) in the nonsig trait category (SNP-heritability p-value > 0.05, “likely reflecting limited statistical power rather than a true lack of heritability” by Nealelab) had no new discoveries after data-integration by any of the three methods. Our findings suggest that more informative scores or new data integration methods are warranted to further improve the power of GWAS by leveraging the variant functional annotations.

## Introduction

In the last decade, genome-wide association studies (GWAS) have enabled the discovery and identification of thousands of genetic loci across a wide range of phenotypes (Visscher et al., 2017). However, despite their increasingly large sample sizes (e.g. *n* > 100, 000) there is a need to improve the often modest power of GWAS, considering that genetic effect sizes of truly associated SNPs are believed to be small for most complex human traits (Spencer et al., 2009).

Earlier work have integrated linkage results or summary statistics from independent GWAS of the same or related traits to increase the power of a GWAS (e.g., Buniello et al., 2019, Ott et al., 2015). To integrate information across multiple GWAS, meta-analysis (Cochran, 1954) and Fisher’s method (Fisher, 1938) are two standard and powerful approaches. For example, meta-analysis of summary statistics has been shown to be as powerful as mega-analysis using the original individual-level data, when there is no heterogeneity between the studies (Lin and Zeng, 2010, Sung et al., 2014). On the other hand, Fisher’s method is more robust to differential directions of effect by combining p-values from different studies.

Recently, it has been shown that variant functional annotations can predict the biological relevance of a variant (e.g., Adzhubei et al., 2010, Davydov et al., 2010, Dunham et al., 2012, Kundaje et al., 2015). To overcome limitations such as incomparable metrics of measurement and differential ascertainment biases across different annotations, several authors have proposed methods to integrate many diverse annotations into one single measure: a meta-score (e.g., Ionita-laza et al., 2016, Kircher et al., 2014, Lu et al., 2015, Ritchie et al., 2014, Shihab et al., 2015). For instance, Kircher et al. (2014) combined more than 60 genomic features into one combined annotation dependent depletion (CADD) meta-score to provide a measure of the relative deleteriousness for each variant, while Ionita-laza et al. (2016) developed Eigen, a functional meta-score of similar nature using an unsupervised spectral approach.

Despite the popularity of using these meta-scores for genomic studies (e.g., Li et al., 2020, Liang et al., 2019, Pereira et al., 2019), their potential for improving power of GWAS has not been well studied or understood. To integrate GWAS summary statistics with meta-scores, in addition to meta-analysis and Fisher’s method, we also consider the weighted p-value approach (Genovese et al., 2006)and the stratified false discovery rate (sFDR) control method (Sun et al., 2006), which extended the traditional methodology of FDR control (Benjamini and Hochberg, 1995). Both weighted p-value and sFDR methods have been used to leverage linkage evidence (e.g., Roeder et al., 2006, Yoo et al., 2010), gene-expression data (e.g., Keel et al., 2020, Li et al., 2013), and pleiotropy (e.g., Andreassen et al., 2013) to increase power of GWAS. Here we use these data-integration methods to integrate CADD or Eigen functional meta-scores with GWAS summary statistics of 1,132 phenotypes from the UK Biobank data (Sudlow et al., 2015).

Integrating functional annotation scores with GWAS summary statistics has been previously studied. Recently, Kichaev et al. (2019) proposed a modified weighted p-value-based method called FINDOR to leverage polygenic functional enrichment to improve power of GWAS. To achieve this, FINDOR uses a stratified linkage disequilibrium (LD) score regression method (Finucane et al., 2015) to compute the expected 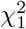 statistic for each GWAS SNP, by regressing the observed GWAS 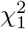 statistics against 75 functional annotations (Gazal et al., 2017) that are tagged by each SNP. FINDOR then stratifies the GWAS SNPs into 100 equally-sized bins based on their expected GWAS 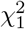 values and applies bin-specific weights to the corresponding GWAS p-values. An application of FINDOR by Kichaev et al. (2019) to 27 traits selected from the UK Biobank data (Sudlow et al., 2015) showed that the method was able to improve power of GWAS by identifying additional associated variants.

Focusing on integrating functional meta-scores with GWAS summary statistics to improve power of GWAS, our study here is different from the FINDOR evaluation of Kichaev et al. (2019) in several ways. Firstly, unlike FINDOR, we study methods that prioritize GWAS findings based on external information alone. That is, the weighting factor and stratification are determined solely based on the annotations, independent of the observed GWAS summary statistics to minimize potential over-fitting. Secondly, instead of using many individual functional scores we utilize existing meta-scores such as CADD and Eigen, which are already calibrated and easier to implement in practice. Thirdly, we focus on evaluating methods’ robustness to the possibility of uninformative functional annotations, because our understanding of the genomic functionality of a genetic variant is incomplete and evolving. Finally, we comprehensively examine all 1,132 UK Biobank traits for which the confidence for their SNP-heritability estimates were considered medium to high by Benjamin Neale’s lab from the Broad Institute (hereafter referred to as Nealelab; Web Resources).

We first provide technical details for the four data-integration methods to be examined, namely meta-analysis, Fisher’s method, weighted p-value, and sFDR. We then describe the genetic and genomic data to be integrated, namely the summary statistics of the 1,132 UK Biobank traits from Nealelab, and the CADD and Eigen functional annotation meta-scores from their respective websites (Web Resources).

For a comprehensive evaluation of the methods, we first detail our simulation study designs, including leveraging the observed genomic data combined with simulated genetic data or vice versa, or using only simulated data. We then describe in details the various measures used to compare method performance, including the traditional family-wise error rate (FWER), as well as FDR, power, recall, precision, and relative efficiency.

In addition to the extensive simulation studies, our empirical method evaluation includes an applications that integrates the UK Biobank GWAS summary statistics with CADD or Eigen meta-scores, analyzing close to 8 million SNPs for 1,132 complex traits. When appropriate, we also include the FINDOR method (using 75 individual functional scores) in both the simulation and application studies.

## Materials and Methods

### The integration methods: meta-analysis, Fisher’s method, weighted p-value, and stratified false discovery rate (sFDR) control

#### Notation and set-up

Let *z_i_* and *p_i_* be the association test statistic and its corresponding p-value for SNP *i, i* = 1, …*m*, from a genome-wide association study, the primary data of interest. Without loss of generality, we assume *z_i_* follows *N*(0, 1), the standard normal distribution, under the null hypothesis of no association between the SNP and the GWAS trait under the study.

Let *z_i,add_* and *p_i,add_* be additional information available for the SNP, based on data *independent* of *z_i_* and *p_i_* from the GWAS. Note that *z_i,add_* may or may not be normally distributed depending on the application setting, e.g. *z_i,add_* can be the CADD (Kircher et al., 2014) or Eigen (Ionita-laza et al., 2016) functional meta-score available for SNP *i*, which will be the focus of our study.

#### Meta-analysis and Fisher’s method

For the meta-analysis approach, we first assume the best-case scenario where *z_i,add_* is normally distributed. We then use the inverse variance approach (Hedges and Vevea,1998) to integrate *z_i_* and *z_i,add_*,

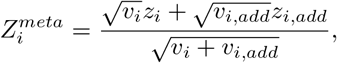

where the weights, *v_i_* and *v_i,add_*, are inverse variance estimates associated with *z_i_* and *z_i,add_*, respectively, from the GWAS study and the additional study available for data integration. Under the null hypothesis of no association *and* assuming the functional meta-score is uninformative, 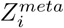 is *N*(0, 1) distributed.

Fisher’s method combines p-values instead of the test statistics. That is,

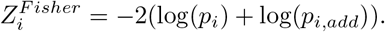

Fisher’s method is omnibus to directions of effect, and as a result it can be more powerful than meta-analysis when signs of *z_i_* and *z_i,add_* differ. Under the null that both p-values, *p_i_* and *p_i,add_* are, independently, Unif(0,1) distributed, 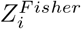 is 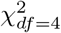 distributed.

Although meta-analysis and Fisher’s method are applicable in many scientific settings, their applications to genetic association studies are typically restricted to combining association evidence from multiple GWAS studies of the same phenotype in the same population. This is because the statistical power of meta-analysis (and Fisher’s method) relies on the assumption of homogeneity beyond direction of effect (Thompson, 1994). In practice, given two families of multiple tests, the underlying compositions of the null and alternative hypotheses may differ, unless the two studies used the same study design and data ascertainment scheme, including phenotype definition, genotyping platform, environmental exposure, and study population (Begum et al., 2012). When the truly associated SNPs do not completely overlap between the different studies, using random-effect instead of fixed-effect meta-analysis does not guarantee improved power, because it violates the assumption that the effect sizes come from the *same* distribution.

The use of meta-analysis and Fisher’s method is also questionable when *z_i_* and *z_i,add_* from the two studies offer different types of information. In essence, the use of weights *v_i_* and *v_i,add_* notwithstanding, meta-analysis and Fisher’s method implicitly assume *z_i_* and *z_i,add_* carry ‘exchangeable’ information. For our study, however, *z_i_* is the genetic association summary statistic, while *z_i,add_* is the genomic annotation meta-score. Thus, meta-analysis and Fisher’s method are likely to be sub-optimal for the purpose of this study. However, for completeness we include the two classical data-integration methods in our initial method evaluation.

For a practical implementation of meta-analysis when *z_i,add_* is the CADD or Eigen metascore, we let *v_i,add_* = *v_i_* as it is unclear what is the variance of *z_i,add_*, the functional meta-score; using the sample estimate is not appropriate as it assumes the underlying variances of *z_i,add_*’s are the same across the *i* = 1, …*m* SNPs. Further, we use the inverse normal transformation to re-scale *z_i,add_* while keeping the sign of the re-scaled *z_i,add_* to be the same as *z_i_*, creating the best-case scenario for the meta-analysis; see Supplementary Information S.1 for details. Similarly, for a practical implementation of Fisher’s method, we use a rank-based transformation and let *p_i,add_* = (rank of *z_i,add_*/*m*), which is also related the phred-scaled CADD and Eigen scores which we discuss later.

#### The weighted p-value approach

Unlike meta-analysis and Fisher’s method, which assume *z_i_* and *z_i,add_* carry similar information, the weighted p-value approach (Genovese et al., 2006) treats *z_i_* and *z_i,add_* differently. That is, the method considers *z_i_* and *p_i_* as the primary data of interest, and it converts *z_i,add_* to *w_i_*, a weight to be applied to *p_i_*. Thus, the weighted p-value approach is an attractive method for the purpose of our study, where the primary data are GWAS summary statistics, and the additional information available are genomic functional scores derived *independently* from the GWAS of interest.

For a valid weighted p-value implementation, the *w_i_*’s must satisfy two conditions: *w_i_* ≥ 0 and 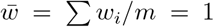 (Genovese et al., 2006). To convert *z_i,add_* to *w_i_*, Roeder et al. (2006) studied two possible weighting schemes: exponential, *w_i_* = *m*(exp(*β* × *z_i,add_*)/∑_*i*_ exp(*β* × *z_i,add_*)), and cumulative,

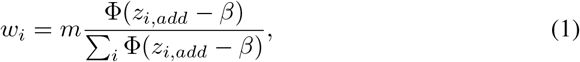

where Φ is the cumulative distribution function of the standard normal. In either case,

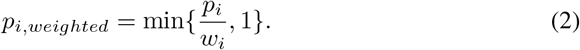

Here we choose the cumulative weighting scheme, with the recommended default value of *β* = 2 (Roeder et al., 2006). This is because the exponential converting scheme is highly sensitive to large values of *z_i,add_*, which is the case here; functional meta-scores can be as large as 80 (Kircher et al., 2014).

#### Stratified false discovery rate (sFDR) control

Unlike the weighted p-value approach that up-weights or down-weights each SNP according to its external information *z_i,add_*, the sFDR method separates the GWAS SNPs into different groups based on *z_i,add_*, which can be continuous or categorical (Sun et al., 2006). When *z_i,add_* is continuous, it has been shown that categorizing *z_i,add_* does not necessarily result in loss of power, as the additional information available are unlikely to be precise or completely informative (Yoo et al., 2010). In addition, sFDR is robust to the situation when *z_i,add_* is uninformative (i.e. random) or possibly misleading.

To implement sFDR in our setting where *z_i,add_* is the continuous functional meta-score, without loss of generality, we first stratify GWAS SNPs into two groups based on whether their meta-scores are among the top five percent or not, *irrespective* of *z_i_* and *p_i_*, the GWAS summary statistics. (The choice of the number of groups and the thresholds, however, is subjective, similar to the choice of the weighting scheme and *β* value for the weighted p-value approach above.) As a result, there are two groups of GWAS SNPs, where group 1 contains 5% of the GWAS SNPs with the highest functional meta-scores and group 2 contains the remaining SNPs. It is worth emphasizing that group 1 is only presumed to be the high-priority group, as the stratification is based on genomic *z_i,add_* alone, independent of the GWAS *z_i_* or *p_i_*.

We then apply FDR control, separately, to the two groups of GWAS p-values, but using the same pre-specified FDR *γ*% level. Following the sFDR method of Sun et al. (2006), for each group of SNPs we first convert their GWAS p-values, *p_i_*’s, to q-values, *q_i_*’s (Storey, 2002), and we then reject the SNPs with *q_i_* < *γ*%; this sFDR procedure controls the overall FDR at the *γ*% level. Although sFDR does not explicitly use weights, it has been shown to be a robust version of the weighted p-value approach, where all SNPs within a group have the same weights (Yoo et al., 2010).

Let *m^k^* be the number of SNPs in group *k*, and let 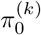 be the proportion of truly associated SNPs in the group. Within each group, we obtain q-values recursively (Storey,2002),

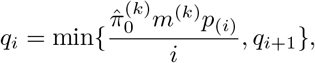

where *p*_(1)_ ≤ … ≤ *p*_(*i*)_ ≤ … ≤ *p*_(*m*)_ are the ordered GWAS p-values, and the procedure starts from 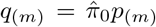. To obtain 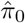, we choose the commonly used conservative estimate (Storey and Tibshirani, 2003),

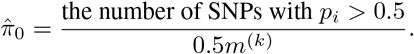

After rejecting SNPs with *q_i_* < *γ*% separately for each group of SNPs, group-specific weights can be inferred if desired(Yoo et al., 2010). Let *α*^(*k*)^ be the maximum GWAS p-values among the rejected SNPs for group *k*, the group-specific weight is,

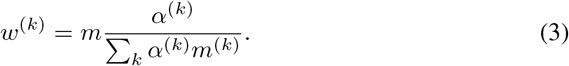

We can then obtain sFDR weighted p-values,

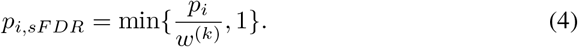

Without loss of generality, if group 1 has no rejections at the pre-specified FDR *γ*% level, we set *w*^(1)^ = 0 and *w*^(2)^ = *m*/*m*^(2)^. If both groups have no rejections at the *γ*% level, then *w*^(1)^ = *w*^(2)^ = 1. That is, the study is reduced to the unweighted case.

The sFDR group-specific weights, *w*^(*k*)^’s satisfy the constraints imposed by the weighted p-value approach (Genovese et al., 2006), and they have been shown to be a robust version of the SNP-specific *w_i_*’s (Yoo et al., 2010). If the additional information is truly informative, *w*^(1)^> 1 while *w*^(2)^ < 1, where *w*^(*k*)^’s can be considered as dichotomized *w_i_*’s of the weighted p-value approach. In that case, the weighted p-value approach is slightly more powerful than sFDR. On the other hand, if the information is just random noise, *w*^(1)^ ≈ *w*^(2)^ ≈ 1 for sFDR, while the weighted p-value method still up-weights or down-weights the GWAS p-values according to the individual *w_i_*’s values, which are proportional to the observed *z_i,add_*’s. In the event of misleading information, *w*^(1)^ < 1 while *w*^(2)^ > 1 even though group 1 was presumed to be the high-priority group. Thus, sFDR is robust to uninformative or even misleading added information.

### The UK Biobank GWAS summary statistics for 1,132 complex traits

We obtained the UK Biobank GWAS round 2 summary statistics from Nealelab (Web Resources). Nealelab performed association studies for 4,236 complex traits using regression, where the regression models included age, sex and the first 20 principal components as covariates, in addition to SNP genotypes which were coded additively. For each of these traits, Nealelab also applied the LD-score regression method (Bulik-Sullivan et al., 2015) to estimate the SNP-heritability, which ranges from 0% to 48%.

In addition to the point estimate and the p-value of testing if the SNP-heritability is 0%, Nealelab also rated the heritability p-value of each trait with a confidence level, primarily based on the effective sample size (*n* > 20, 000) used for the p-value calculation. Thus, we restricted our analysis to the 1,132 traits rated as with medium to high confidence by Nealelab, among which 531 are continuous and 601 are binary traits for which the effective sample sizes depend on the numbers of cases; see Figure S1 for a histogram of the case rates for the 601 binary traits.

Note that the SNP-heritability estimates for some of the 1,132 traits can still be very low, and their SNP-heritability testing p-values may not be significant. Nealelab then put these 1,132 traits into four groups: nonsig (182 traits with SNP-heritability testing p-value *p* > 0.05), nominal (277 traits; *p* < 0.05), z4 (235 traits; *p* < 3.17 × 10^−5^), and z7 (438 traits; *p* < 1.28 × 10^−12^), where “many of the non-significant results likely reflect limited statistical power rather than a true lack of heritability”. Figure S2 contrasts the 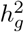 estimates of the 1,132 traits with their z-values of the SNP-heritability testing conducted by Nealelab.

The GWAS of Nealelab were restricted to *n* = 361, 194 individuals of white-British ancestry and 10.9 million variants that passed a set of quality control (QC) steps; see Web Resources for the detailed QC steps performed by Nealelab. Our analysis focused on *m* = 7, 895, 174 common bi-allelic autosomal SNPs. We excluded indel variants because their functional meta-scores are not available. We additionally excluded X-chromosomal variants because their association tests may not be optimal (Chen et al., 2021) and their functional annotations are not always available. Lastly, we excluded SNPs with minor allele frequency (MAF) less than 1% because their association p-values may not be reliable (Tang et al., 2020) and joint analysis of multiple rare variants simultaneously (Derkach et al., 2014) is beyond the scope of our study.

### CADD and Eigen functional meta-scores

We obtained the CADD meta-scores (v1.6), using the CADD tool (Rentzsch et al., 2019), and the Eigen meta-scores (v1.0), using the ANNOVAR tool (Wang et al., 2010), for all the 7,895,174 common bi-allelic autosomal SNPs.

In addition to the raw CADD meta-scores, the CADD tool also provides rank-based rescaled scores called phred scores, −10 log_10_(ranks of the raw scores/total number SNPs), which are positive and have better interpretation as compared with the raw scores. For example, a phred score of 10 or greater indicates that the SNP is predicted to be among the top 10% most deleterious among the human genome, while a phred score 20 or greater implies top 1% most deleterious.

For consistency between CADD and Eigen, we applied the same technique to obtain phred-scaled Eigen scores; hereafter scores represent phred-scaled scores unless specified otherwise. Figure S3 in Supplementary Information shows the histograms of CADD and Eigen scores; each is expected to be 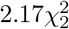 distributed, because (rank of the raw score/total number SNPs) is Unif(0,1) distributed, and as a result −2 log(Unif(0,1)) is 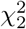 distributed.

Because Eigen scores were estimated using an unsupervised learning approach, in contrast to CADD scores, which were estimated using labeled data, we also compared these two scores genome-wide (Figure S4) and across four different consequence categories: missense, non-coding, synonymous, and protein truncating variants (PTV) (Figure S5). Variants in the missense and PTV categories tend to have higher CADD than Eigen scores, while variants in the non-coding and synonymous categories tend to have higher Eigen than CADD scores. However, overall the two meta-scores are consistent and lead to qualitatively comparable data-integration results, which we discuss next.

### Simulation study design I, leveraging the observed genomic data

Here we used the real CADD and Eigen functional meta-scores, combined with simulated GWAS summary statistics, to verify type I error control of the studied data-integration methods.

#### Simulated GWAS summary statistics under the null of no association combined with real functional annotation scores

We first simulated a *N*(0, 1) distributed trait for 1,756 individuals from the 1000 Genomes Project (Auton et al., 2015). The phenotype values were simulated *independently* of the genotypes of 422,923 bi-allelic, autosomal and common (MAF > 5%) SNPs that a) passed quality control conducted by Roslin et al. (2016) who studied the individuals from the five super-populations, and b) have available CADD and Eigen meta-scores, and c) have the 75 annotations used by FINDOR.

We then obtained GWAS summary statistics for the 422,923 SNPs by regressing the trait values of the 1,756 individuals on the additively coded genotypes. Because the trait values were randomly generated, independent of the genotypes and the populations, the resulting GWAS *z_i_*’s are *N*(0, 1) distributed and *p_i_*’s Unif(0,1) distributed, as expected under the null of no association and confirmed by the histograms of *z_i_*’s and *p_i_*’s from one randomly selected simulation run (Figure S6).

Finally, we integrated the GWAS summary statistics with their corresponding CADD (or Eigen) meta-scores using the four methods, meta-analysis, Fisher’s method, weighted p-value, and sFDR control as described above. Although Kichaev et al. (2019) showed that FINDOR calibrated well when a GWAS consists of a mixture of null and associated SNPs, we also examined the performance of FINDOR in this setting when all GWAS SNPs are under the null hypothesis of no association. We applied the FINDOR tool using the same set of LDscores and the 75 annotations (Gazal et al., 2017) that were used by Kichaev et al. (2019) for their study; see Web Resources.

#### Method evaluation: family-wise error rate (FWER)

For each simulation replicate (i.e. a GWAS simulated under the null of no association), we obtained the number of false positives using the conservative Bonferroni corrected significance level, *α* = 0.05/422923 = 1.2 × 10^−7^. We repeated the simulation, independently, 50,000 times, and calculated the family-wise error rate as the proportion of the number of replicates with at least one significant finding. Assuming the true FWER is 0.05, we expect the estimate obtained from the 50,000 independent simulation replicates to have a standard error of 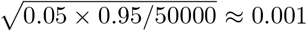. Thus, a method with a FWER estimate outside [0.047, 0.053] can be considered inaccurate.

### Simulation study design II, leveraging the observed genetic data

Here we used the UK Biobank GWAS summary statistics, combined with permuted CADD and Eigen scores, to evaluate robustness of the methods to random annotation scores. Prior to the permutation, we examined the similarity of functional annotations between SNPs in linkage disequilibrium.

#### Permuted functional annotation scores combined with real GWAS summary statistics

To obtain a null set functional annotation meta-scores that are independent of GWAS summary statistics, we randomly permuted the observed CADD (or Eigen) scores between the SNPs. Although such permutation does not preserve the potential correlation between functional scores of nearby SNPs, for the purpose of this study, it provides a valid set of annotation scores that are independent of the GWAS summary statistics. Nevertheless, we examined the similarity of annotation scores between SNPs in linkage disequilibrium.

Using CADD as an example, let *CADD_i_* and *CADD_j_* be the annotation scores of SNPs *i* and *j*, respectively. We first defined a pair-wise similarity measure as 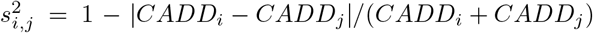, which is bounded between 0 and 1, where 1 means two scores are identical whereas a value close to 0 suggests a lack of similarity. We then contrast 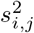 with 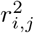, the traditional LD measure of genotype similarity between two SNPs. Results in Figure S7 show that there is no clear concordance between the two measures. A closer examination of 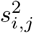 and 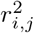 for two randomly selected regions in Figure S8, and the contrast between variant-specific CADD meta-score and LD score across the genome in Figure S9, led to the same conclusion that functional scores of SNPs in LD are not necessarily similar.

After we permuted the functional scores of the 7,895,174 common, bi-allelic autosomal SNPs, for each of the 1,132 traits of the UK Biobank data, we integrated the GWAS summary statistics with the permuted annotation scores, using meta-analysis, Fisher’s method, weighted p-value, and sFDR control. We were not able to evaluate FINDOR here, because FINDOR implements the LD SCoring (LDSC) tool (Bulik-Sullivan et al., 2015) and the validity of using LDSC for permuted annotations is not clear.

#### Method evaluation: *Recall*, *Precision* and *FDR*

Before data integration, we first used *α* = 5 × 10^−8^ to identify genome-wide significance findings (Dudbridge and Gusnanto, 2008) based on the summary statistics of the 1,132 UK Biobank GWAS by Nealelab. For the purpose of this simulation study, we treated these findings, *m*_1,*t*_, as the total number of truly associated SNPs to be discovered after data-integration for each trait *t*, *t* = 1,…, 1, 132. In addition to counting the number of significant SNPs per GWAS, we also counted the number of significant independent loci. We first defined independent loci using the LDclumping algorithm of PLINK (v1.07) (Purcell et al., 2007), with a sliding window of 1 Mb and a LD *r*^2^ threshold of 0.1 as per standard practice. We then considered each independent locus significant if the locus contained at least one genome-wide significant SNP.

After data-integration, integrating the GWAS summary statistics with the *permuted* functional scores for each trait *t*, we then used the same *α* = 5 × 10^−8^ to identify genome-wide significance findings (SNPs or loci as defined above), denoted as *P_t_*. Among the *P_t_* positives, we defined false positives, *FP_t_*, as the new findings that were not part of the *m*_1,*t*_ findings, because the information used for data integration were permuted functional scores. Similarly, we defined *TP_t_* = *P_t_* − *FP_t_* as the number of true positives for trait *t*.

Finally, we defined and calculated recall, precision and false discovery rate by

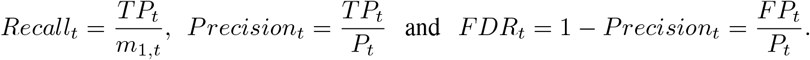

*Recall* is conceptually the same as *Power*, defined later for simulation studies where we know the ground truth and *m*_1,*t*_ SNPs were simulated as truly associated SNPs. We calculated *Recall_t_* only when *m*_1,*t*_ > 0. That is, for the 409 out 1,132 GWAS with no significant findings *before* data-integration (i.e. *m*_1,*t*_ = 0) we did not calculate *Recall_t_*. Regardless of if *m*_1,*t*_ = 0, for GWAS with no significant findings *after* data-integration, (i.e. *P_t_* = 0; 558 by meta-analysis, 505 by Fisher’s method, 440 by weighted p-value, and 415 by sFDR), we conservatively defined *Precision_t_* = 1 and *FDR_t_* = 0.

### Simulation study design III, varying the informativeness of genomic information

To further investigate method performance in the presence of completely informative, partially informative, uninformative, or even misleading added information, we performed an additional set of simulation studies. Although linkage disequilibrium is an important aspect of GWAS, given the simulation study design I and our findings in the results section, the simulation studies here focused on independent SNPs to delineate other potentially influencing factors.

Without loss of generality, we assumed the total number of SNPs *m* = 10, 000, among which the first *m*_1_ = 100 SNPs are truly associated. The corresponding summary statistics *z_i_*’s were drawn, independently, from *N*(*μ*_1_, 1) for the *m*_1_ associated SNPs, and from *N*(0, 1) for the remaining null SNPs. The top left plot in Figure S10 shows the Manhattan plot for one simulated GWAS replicate with *μ*_1_ = 3; we also varied *μ*_1_ from 0.1 to 4 to represent different power of a GWAS.

We then assumed *z_i,add_*’s as the additional information available, which were drawn, independently, from *N*(*μ_add_*, 1) for *m_add_* SNPs and *N*(0, 1) for the remaining SNPs. Importantly, the locations of the *m_add_* SNPs may differ from those of the *m*_1_ associated SNPs. That is, the additional information available for a truly associated SNP may be random noise. On the other hand, for a null SNP with no association (i.e. *z_i_* drawn from *N*(*μ*_1_,1)), *z_i,add_* could be drawn from *N*(*μ_add_*, 1), representing misleading information. We also varied *μ_add_*, which may or may not be the same as *μ*_1_.

Using *m*_1_ = 100 and *μ*_1_ = 3 as an example for the GWAS component, we considered the following eight scenarios for the additional information available for data integration (Figure S10), which fall into four categories.

- Category I is completely informative (homogeneity): (1) *m_add_* = 100, *μ_add_* = 3 and locations of the *m_add_* SNPs perfectly match those of *m*_1_ GWAS truly associated SNPs.
- Category II is partially informative: (2) *m_add_* = 100 and *μ_add_* = 1.5; (3) *m_add_* = 50 and *μ_add_* = 3; (4) madd = 50, *μ_add_* = 1.5, and all madd SNPs coincide with (some of) the *m*_1_ SNPs.
- Category III is (partially or completely) misleading: (5) *m_add_* = 100 and *μ_add_* = 3; (6) *m_add_* = 100 and *μ_add_* = 1.5, but in both scenarios only 50 out of the *m_add_* SNPs coincide with 50 of the *m*_1_ SNPs. And (7) *m_add_* = 100 and *μ_add_* = 3, but none of the *m_add_* SNPs coincide with the *m*_1_ SNPs.
- Category IV is uninformative: (8) *m_add_* = 0 and *μ_add_* = 0. That is, the additional information available is white noise.

For each of the eight scenarios, we simulated 1,000 data replicates, independently of each other. For each replicate, we then applied the four data-integration methods that are suitable for this simulation study, namely meta-analysis, Fisher’s method, weighted p-value, and sFDR control. Finally, we evaluated the methods using various performance measures, which we describe below.

#### Method evaluation: power and relative efficiency (RE)

We first used the Bonferroni corrected threshold to declare significance (i.e. p-value < 0.05/10000 = 5 × 10^−6^). Let *P_rep_t__* be the number of positives for each of the 1000 simulation replicates after data-integration, we defined power as

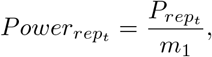

the proportion of the truly associated GWAS SNPs that were found after data-integration, which is similar to *Recall_t_* defined earlier in simulation study design I.

As the Bonferroni approach can be conservative, we explored two alternative decision rules: fixed-region and fixed-FDR rejections. The fixed-region rule rejected the top *k* SNPs (e.g. *k* = 100), while the fixed-FDR rule rejected SNPs by controlling FDR at *γ*% level (e.g. *γ*% = 5%or 20%). For each rejection rule, we then calculated the power as described above.

In the context of multiple hypothesis testing, the performance of a method can be evaluated by alternative measures, such as relative efficiency. To this end, we first ranked all the truly associated *m*_1_ SNPs based on the GWAS data summary statistics alone, denoted as *R_baseline_*. We then ranked these SNPs after data-integration, denoted as *R_method_*, based on the magnitude of their *Z^meta^, Z^Fisher^, p_weighted_*, and *p_sFDR_* statistics as defined in the method section. Finally, after averaging *R_baseline_* and *R_method_* across the 1000 simulated replicates, we defined relative efficiency as

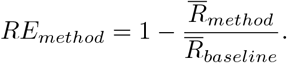

A positive *RE_method_* value means the truly associated *m*_1_ SNPs are ranked higher, on average, after data-integration using the method; a *RE_method_* value of zero means that the date-integration method did not improve performance; and a negative *RE_method_* value suggests that the data-integration effort was counter-productive.

### Application, integrating UK Biobank GWAS summary data with functional annotations

We analyzed the 7,895,174 common, bi-allelic autosomal SNPs of the UK Biobank data, using the weighted p-value approach and sFDR methods to integrate their GWAS summary statistics with the CADD (or Eigen) meta-scores, for each of the 1,132 traits as described before. We excluded meta-analysis and Fisher’s method from data application, because severe robustness issues (to partially informative, uninformative, or misleading *z_i,add_*) were found in simulation studies; see results for details. For comparison, we also implemented FINDOR using the set of 75 publicly available annotations recommended by the authors; see Web Resources.

To evaluate method performance, we first counted the total number of significant, independent loci (at the 5 × 10^−8^ level) found after data integration, the same measure used by FINDOR. As a baseline, we also counted the number of significant loci identified based on the GWAS data alone (without data integration with annotation scores) for each of the 1,132 traits. We then calculated *Recall*, the proportion of the initial GWAS findings that were retained after data-integration, as defined early for the simulation studies. Finally, we used *New Discoveries* to represent the number of new genome-wide significant findings at the 5 × 10^−8^ level.

## Results

### Results of simulation design I, leveraging the observed genomic data

Here we integrated real functional annotations with simulated GWAS summary statistics that were simulated under the null of no association. The performance measure here is the empirical family-wise error rate, estimated from 50,000 simulated replicates.

Table 1 shows that the empirical FWER is 0.0496 for the baseline analysis (i.e. using the null GWAS summary statistics alone). For the five different data-integration methods, the empirical rates were 0.0477, 0.0366, 0.0501, 0.0474, and 0.0537 for meta-analysis, Fisher’s method, weight p-value, sFDR, and FINDOR, respectively, where FINDOR is the only method with slightly liberal type I error rate.

**Table 1:**
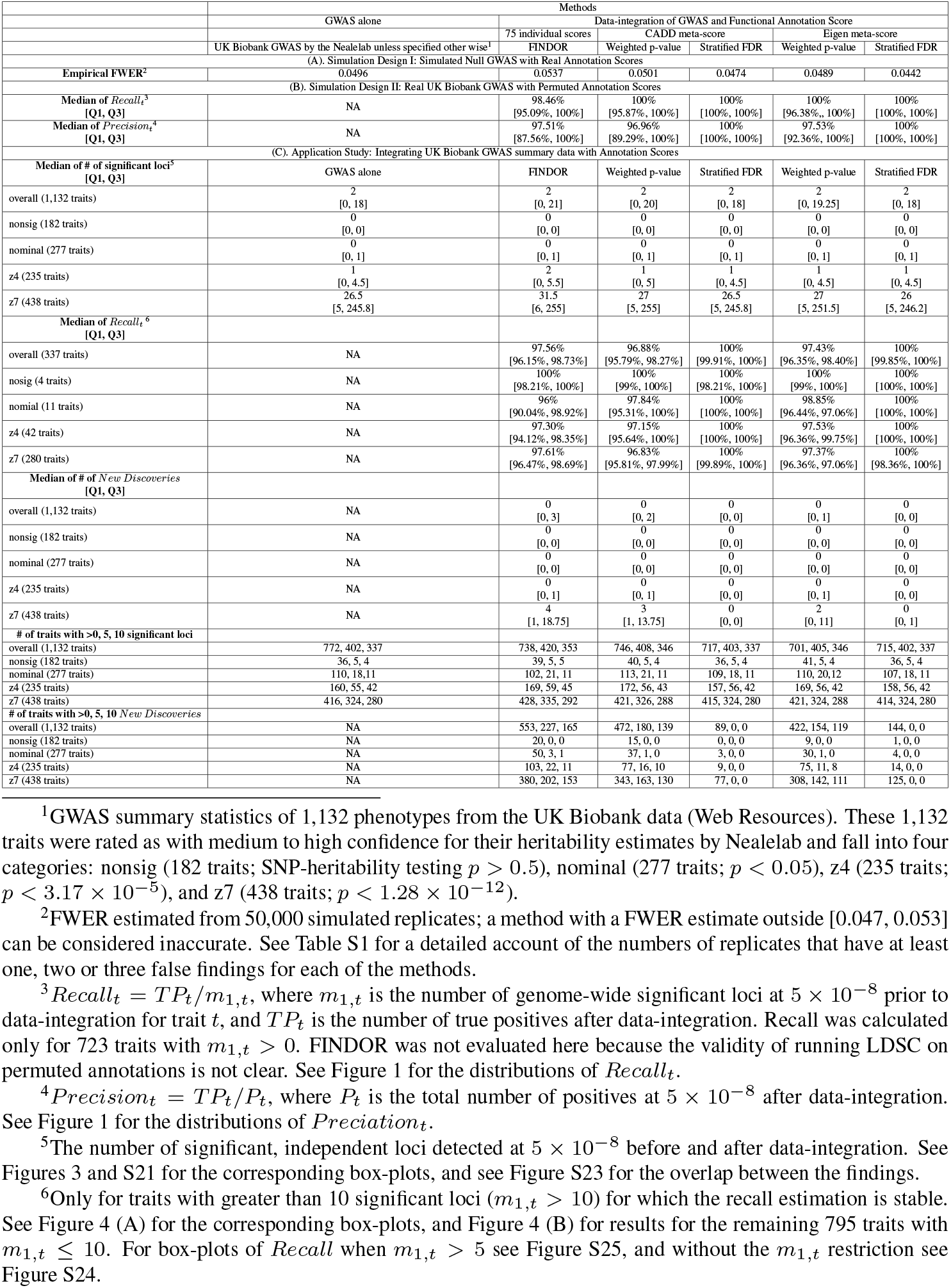
A summary of the simulation and application results.

Table S1 also provides a detailed account of the numbers of replicates, out of a total of 50,000 replicates, that have at least one, two or three false findings for each of the methods; no method had more than three false findings per GWAS. Although a method with an empirical FWER estimate outside [0.047, 0.053] can be considered inaccurate, overall all methods have reasonable type I error control in this setting.

### Results of simulation design II, leveraging the observed genetic data

Here we integrated real UK Biobank GWAS summary statistics of the 1,132 traits with *permuted* CADD (or Eigen) meta-scores; FINDOR was not applicable here. The suitable performance measures here are *Recall_t_* = *TP_t_*/*m*_1,*t*_ and *Precision_t_* = 1 − *FDR_t_* = *TP_t_*/*P_t_*, where *m*_1,*t*_ was defined as the number of genome-wide significant GWAS findings prior to data-integration for trait *t*, and *P_t_* and *TP_t_* are the numbers of positives and true positives after data-integration.

The *Recall* results shown in Figure 1 confirm that meta-analysis and Fisher’s method are not suitable for integrating functional annotations with GWAS summary statistics. Among the 723 GWAS with at least one significant finding prior to data integration (*m*_1,*t*_ > 0), 574, 627, 717, and 692 GWAS have at least one significant finding after data-integration using, respectively, meta-analysis, Fisher’s method, weighted p-value, and sFDR to integrated permuted CADD scores.

**Figure 1:**
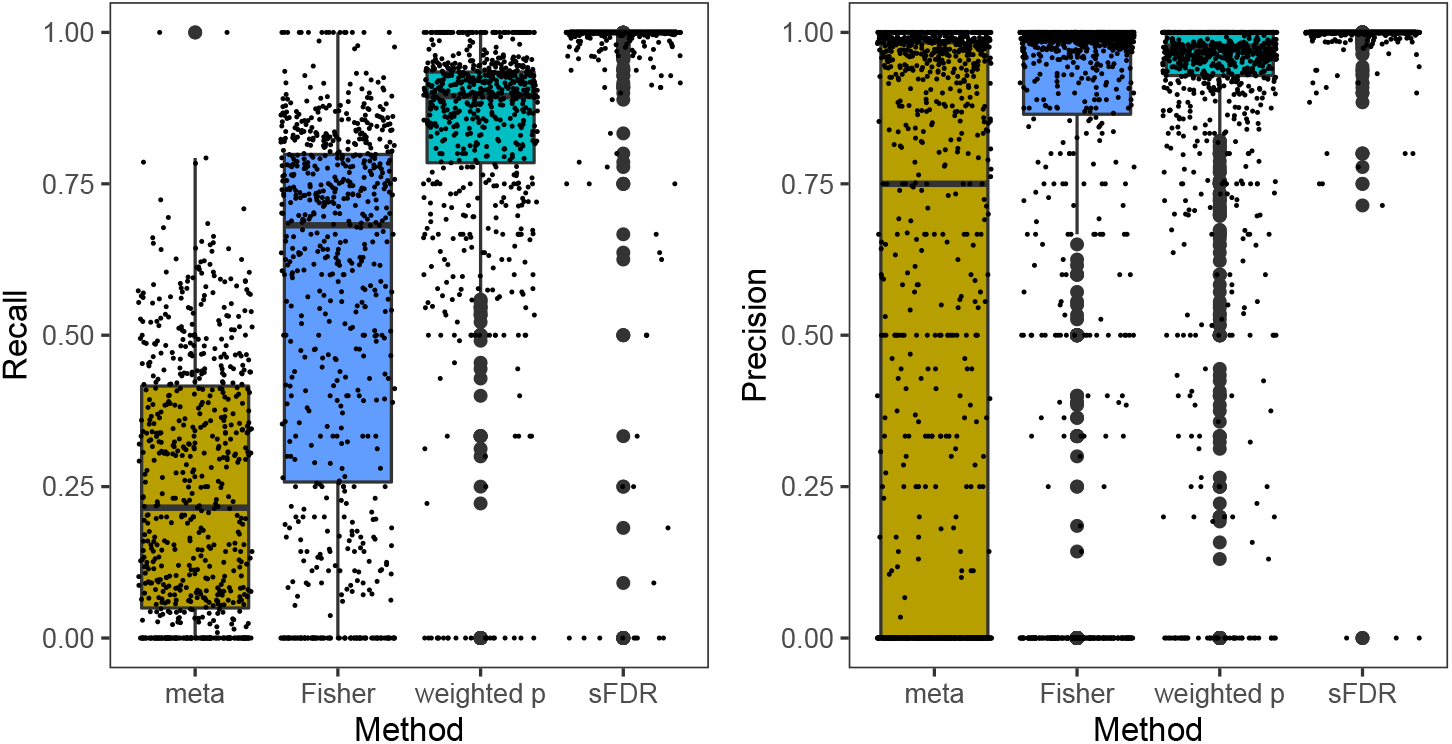
The *Recall* and *Precision* rates obtained from simulation study design II, integrating the 1,132 UK Biobank GWAS summary statistics with *permuted* CADD functional meta-scores, using meta-analysis, Fisher’s method, the weighted p-value approach, and the stratified FDR control. *Recall_t_* = *TP_t_*/*m*_1,*t*_ and *Precision_t_* = 1 − *FDR_t_* = *TP_i_*/*P_i_*, where *m*_1,*t*_ is the number of genome-wide significant independent loci prior to data-integration for trait *t*, and *P_t_* and *TP_t_* are the numbers of positives and true positives after data-integration; see Table 1 for additional results. Independent loci were defined using PLINK’s LDclumping algorithm with a 1 Mb window and an *r*^2^ threshold of 0.1.

Across the 723 GWAS with *m*_1,*t*_ > 0, the median [Q1, Q3] *Recall* rates are 66.67% [50%, 73.34%] for meta-analysis and 84.23% [70%, 92.15%] for Fisher’s method, after integrating permuted CADD scores. In contrast, these values are 100% [95.87%, 100%] for the weight p-value method and 100% [100%, 100%] for the sFDR control (Figure 1; Table 1). The *Precision* results in Figure 1 and Table 1 are consistent with the *Recall* results.

These results consistently show the sensitivity issue of meta-analysis and Fisher’s methods, and they confirm that sFDR is more robust than the weighted p-value approach which was demonstrated by Yoo et al. (2010) in a different setting, integrating linkage results with GWAS. Results stratified by the four types of traits analyzed, nonsig, nominal, z4, and z7 (Figure S11), counting significant SNPs instead of loci (Figure S12), or using permuted Eigen scores (Figure S13) lead to the same conclusion.

### Results of simulation design III, varying the informativeness of genomic information

Here we simulated GWAS summary statistics with some SNPs truly associated, and we then simulated additional data with varying degree of informativeness, including uninformative or possibly misleading. We used power and relative efficiency (RE) to compare methods, where *RE* was defined as one minus (the average ranks of the truly associated SNP after data-integration) divided by (their average base-line ranks using GWAS data alone).

The *RE* results in Figure 2 are consistent with those from simulation study design II. While meta-analysis and Fisher’s method work well when the additional information is completely informative (i.e. the two data resources are homogeneous with each other), they are not suitable data-integration methods for other settings.

**Figure 2:**
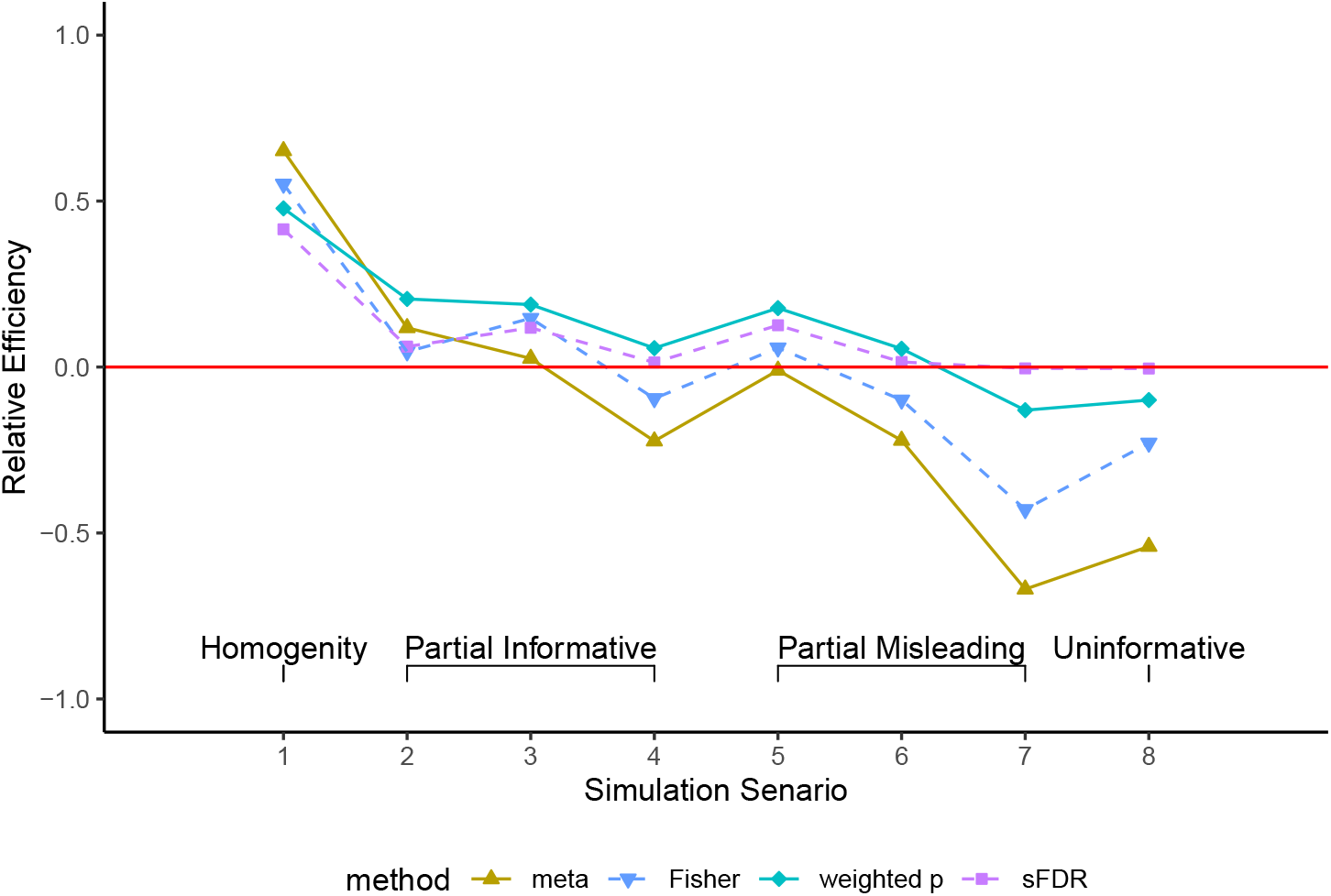
The relative efficiency (*RE*) obtained from simulation study design III, integrating simulated GWAS summary statistics with simulated additional information with varying degrees of informativeness, using meta-analysis, Fisher’s method, the weighted p-value approach, and the stratified FDR control. There were 10,000 independent SNPs, among which 100 were truly associated whose summary statistics were drawn from *N*(3,1); the rest from *N*(0, 1). For the additional information available for data integration, the details of the eight simulation scenarios are provided in the text and illustrated in Figure S10. *RE* is one minus (the average ranks of the truly associated SNP after data-integration) divided (by their average base-line ranks using GWAS data alone), averaged across 1000 simulation replicates.

The power results in Figure S14 are consistent with the *RE* results, across different rejection rules including controlling FWER at 5%, rejecting top 100 ranked SNPs, and controlling FDR at 5% or 20%. Results are also consistent when *μ*_1_ varies from 0.1 to 4 to represent different power of a GWAS (Figures S15–S20). Thus, we excluded meta-analysis and Fisher’s method from the application study.

### Results of the application study, integrating UK Biobank GWAS summary data with functional annotations

We applied three data-integration methods to integrate GWAS summary statistics (between each of the 7,895,174 common SNPs and each of the 1,132 UK Biobank traits) with, respectively, 75 individual annotation scores using FINDOR, and CADD (or Eigen) meta-scores using weighted p-value and sFDR methods. In addition to *Recall* and *Precision*, we also examined the total number of significant, independent loci detected at the 5 × 10^−8^ level, the performance measure used by the original FINDOR paper.

Figure 3 shows the total number of significant, independent loci identified by using GWAS alone (as a baseline), or FINDOR, weight p-value and sFDR data-integration methods, stratified by the four types of traits analyzed (nonsig, nominal, z4, and z7). Overall, integrating existing functional annotations with UK Biobank GWAS association statistics did not lead to striking improvement in terms of the total number of significant loci, as compared with the baseline (i.e. using GWAS data alone) irrespective of the data integration methods. Results of using Eigen (Figure S21) or counting SNPs instead of independent loci (Figure S22) are characteristically similar.

**Figure 3:**
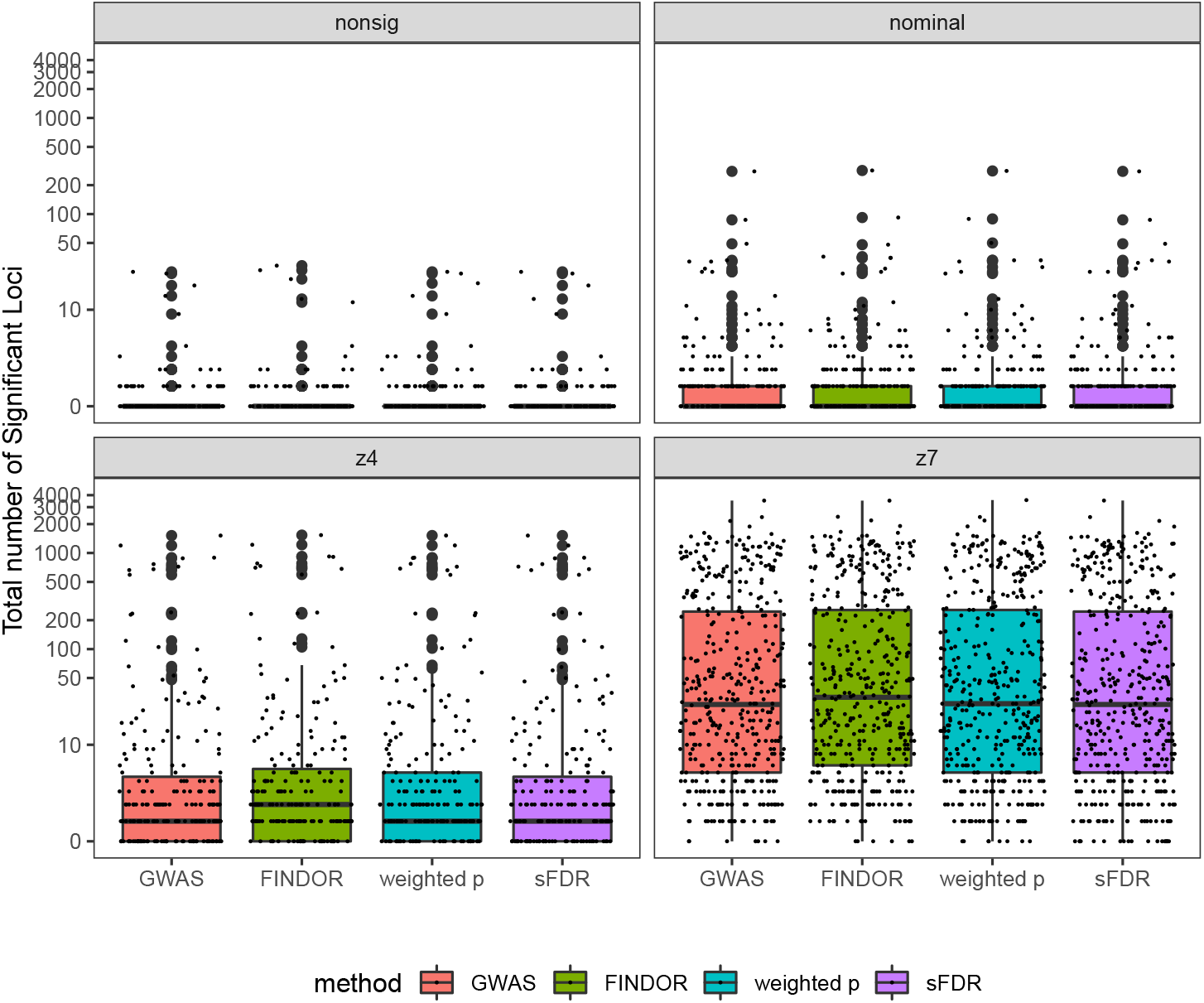
The total numbers of genome-wide significant independent loci of the UK Biobank GWAS application study, before and after data-integration with functional annotations, stratified by the four phenotype categories. In each figure, the total number of significant loci identified based on the UK Biobank GWAS data alone serves as a baseline. The GWAS baseline box-plot is followed by the box-plots for the total numbers of significant loci after integrating the UK Biobank GWAS summary statistics with functional annotations using FINDOR (using 75 individual annotation scores), and the weighted p-value and stratified FDR control methods (each using the CADD meta-score), analyzing 7,895,174 variants for each of the 1,132 UK Biobank traits. The 1,132 traits were rated by Nealelab as having medium to high confidence for their heritability estimates, and they fall into four categories: nonsig (182 traits; heritability testing p-value *p* > 0.05), nominal (277 traits; *p* < 0.05), z4 (235 traits; *p* < 3.17 × 10^−5^), and z7 (438 traits; *p* < 1.28 × 10^−12^). Independent loci were defined using PLINK’s LDclumping algorithm with a 1 Mb window and an *r*^2^ threshold of 0.1.

The loci identified by any of the three methods largely overlapped (Figure S23). For example, in the z7 category 44,595 out 56,153 (79%) genome-wide significant, independent loci discovered are common across the three methods. In general, FINDOR and weighted p-value methods have very similar performance, both with slightly more findings than sFDR, which is not surprising given the trade-off between power and robustness.

Figure S24 shows *Recall* rates stratified by the four trait categories for FINDOR (using a set of 75 annotations) and weighted p-value and sFDR (using CADD meta-scores). Table 1 shows the number of traits with *m*_1,*t*_ > 0 (the number of genome-wide significant findings before data integration), for which *Recall* rates were calculated for each of the four trait categories, as well as the median [Q1, Q3] of *Recall* for each of the three data-integration methods.

Consistent with the simulation results, results in Figure S24 and in Table 1 show that sFDR has the highest *Recall* rates across the four trait categories. However, as the estimation of *Recall* may not be stable when *m*_1,*t*_ is small, Figure 4 (A) shows the *Recall* rates for the 337 traits with *m*_1,*t*_ > 10, while Figure 4 (B) contrasts the number of significant loci preserved after data-integration with that before data-integration, for the remaining 789 traits with *m*_1,*t*_ ≤ 10.

**Figure 4:**
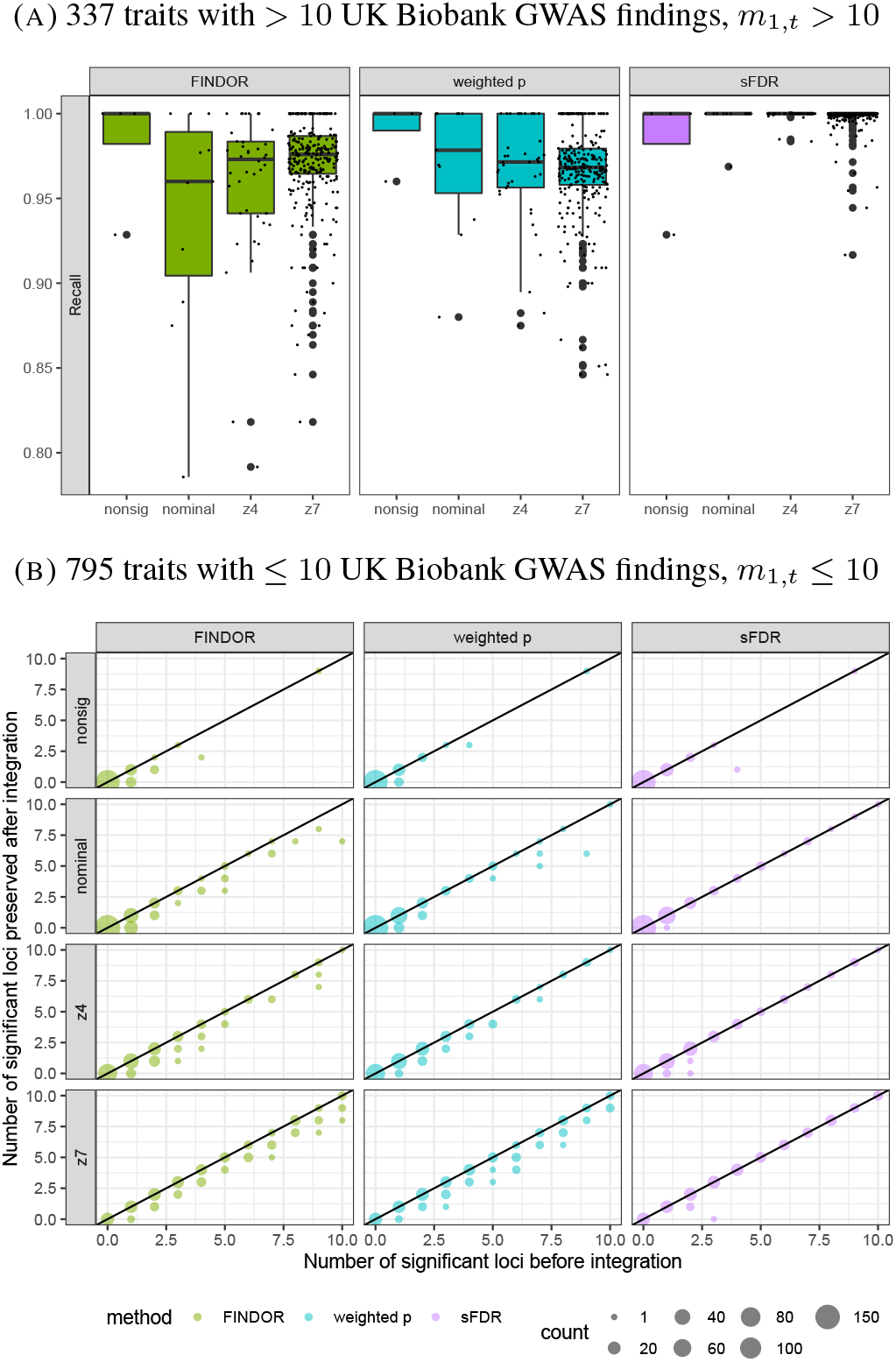
Results of the UK Biobank GWAS application study, before and after data-integration with functional annotations, stratified by the four phenotype categories. (A) *Recall_t_* = *TP_t_*/*m*_1,*t*_, where *m*_1,*t*_ is the number of genome-wide significant independent loci prior to data-integration for trait *t*, and *TP_t_* is the number of true positives after data-integration. *Recall* estimation is not stable when *m*_1,*t*_ is small so for *m*_1,*t*_ ≤ 10, (B) contrasts the number of significant loci preserved after data-integration with *m*_1,*t*_. The three data-integration methods integrated the UK Biobank GWAS summary statistics with functional annotations using FINDOR (using 75 individual annotation scores), and the weighted p-value andstratified FDR control methods (each using the CADD meta-score), analyzing 7,895,174 variants for each of the 1,132 UK Biobank traits. The 1,132 traits were rated by Nealelabas having medium to high confidence for their heritability estimates, and they fall intofour categories: nonsig (182 traits; heritability testing *p* > 0.05), nominal (277 traits; *p* < 0.05), z4 (235 traits; *p* < 3.17 × 10^−5^), and z7 (438 traits; *p* < 1.28 × 10^−12^). Independent loci were defined using PLINK’s LDclumping algorithm with a 1 Mb window and an *r*^2^ threshold of 0.1.

Conclusion drawn from Figure 4 is consist with that from Figure S24. That is, sFDR has better *Recall* rate than FINDOR and weighted p-value approach, which is also supported by results in Figure S25 for *Recall* rates for the 402 traits with *m*_1,*t*_ > 5; results in Figures S26–S28 for the contrasts between the numbers of significant loci before and after data-integration, respectively, for 942 traits with *m*_1,*t*_ ≤ 50, 980 traits with *m*_1,*t*_ ≤ 100, and all 1,132 traits (i.e. *m*_1,*t*_ ≥ 0); results in Figure S29 for *Recall* using Eigen (instead of CADD meta-score) for the weighted p-value and sFDR methods. Interestingly, Figure S24 (A) shows that the *Recall* rate is the highest for traits from the *z*4 category regardless of the data-integration methods. However, Figure S24 (B) shows that overall, *Recall* rate increases as SNP-heritability estimate in creases, which is expected.

Despite the overall high median *Recall* rate, FINDOR and weighted p-value data-integration methods lose much more genome-wide significant loci found in GWAS than the sFDR approach (Figure 4). This behaviour reflects the trade-off between power and robustness as discussed before. Indeed, Figure 5 shows that FINDOR and weighted p-value are better than sFDR in terms of *New Discoveries*; using Eigen instead CADD lead to similar results (Figure S30 and Table 1).

**Figure 5:**
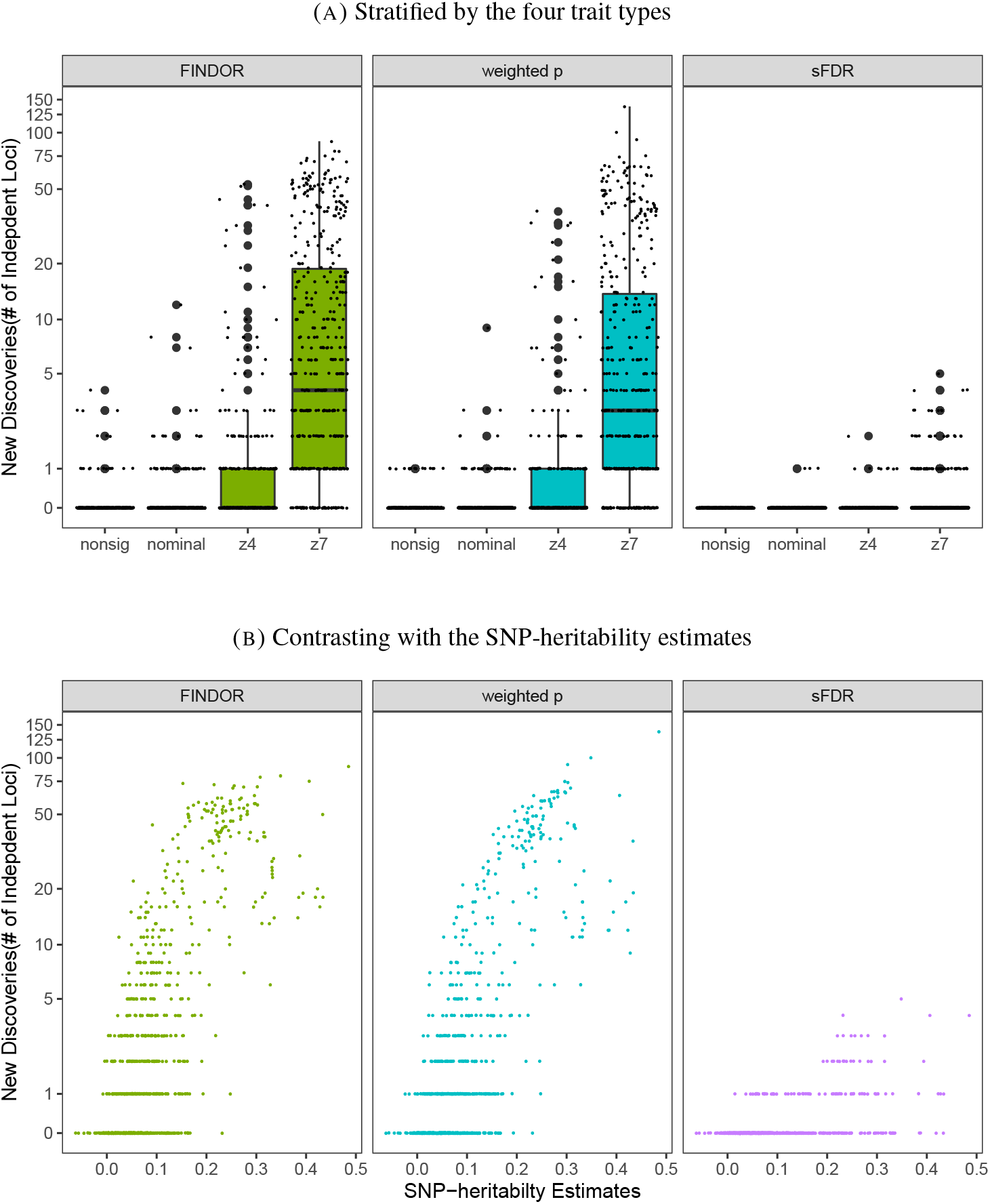
The total *New Discoveries* of the application study (A) stratified by the four trait types or (B) Contrasting with the SNP-heritability estimates. The three data-integration methods integrated the UK Biobank GWAS summary statistics with functional annotations using FINDOR (using 75 individual annotation scores), and the weighted p-value andstratified FDR control methods (each using the CADD meta-score), analyzing 7,895,174 variants for each of the 1,132 UK Biobank traits. The 1,132 traits were rated by Nealelabas having medium to high confidence for their heritability estimates, and they fall intofour categories: nonsig (182 traits; heritability testing *p* > 0.05), nominal (277 traits; *p* < 0.05), z4 (235 traits; *p* < 3.17 × 10^−5^), and z7 (438 traits; *p* < 1.28 × 10^−12^). Independent loci were defined using PLINK’s LDclumping algorithm with a 1 Mb window and an *r*^2^ threshold of 0.1.

Out of the total of 1,132 UK Biobank traits, 553 (48.8%), 472 (41.7%), 89 (7.8%) had *New Discoveries* after using, respectively, FINDOR, weighted p-value and sFDR methods to integrate the GWAS summary statistics with functional annotations; see Table 1 for the counts of *New Discoveries* stratified by the four trait categories. In addition, across the four trait types, 165 traits (15%) and 139 traits (12%) had more than 10 *New Discoveries* after using respectively FINDOR and the weighted p-value data-integration methods. However, the contrast of the numbers of significant loci before and after data-integration in Figures 3 and S28 show that all three data-integration methods have limited capacity to improve the power of GWAS.

## Discussion

We have set out to examine the potential of using functional annotation scores to improve power of genome-wide association studies. We comprehensively studied 1,132 GWAS summary statistics from the UK Biobank data, and two different functional meta-scores (CADD and Eigen) and 75 individual scores, using three different data-integration methods (weighted p-value, stratified FDR and FINDOR). Overall, the number of new genome-wide significant discoveries is limited across traits and methods (Figures 3 and S28), suggesting more informative scores or new data integration methods are needed to further improve power of GWAS by leveraging variant functional annotations. A closer examination of method performance and trait heritability (Figures 5 and S30) revealed that all methods performed better for traits with higher estimates of SNP-heritability. This suggests that even the current UK Biobank-level sample size may not be adequate for some complex traits.

Our study used CADD and Eigen as the functional meta-score available for data-integration. To the best our knowledge, CADD was the first meta-score in the literature and Eigen was the first to use unsupervised learning approach, and both meta-scores have been shown to be superior to other scores in genomic studies. However, the recent work by Li et al. (2020) has proposed annotation-PCs, a new alternative meta-score that warrants further investigation.

In one of our simulation studies, we permuted CADD (and Eigen) to provide a set of meta-scores that are independent of the GWAS summary statistics of the UK Biobank data. Although this approach is valid for examining type I error control, we were intrigued by the question of if there is a meta-score similarity between SNPs in strong linkage disequilibrium. To answer this question, we defined 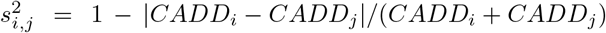 as the similarity measure between SNPs *i* and *j*. Interestingly, there were no clear concordance between 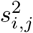 and *r*^2^, the LD measure for genotype similarity (Figures S7 and S8).

Throughout this paper, we have used the default tuning parameter values, *β* = 2 for the weighted p-value approach and *k* = 2 for the sFDR method. We did not tune the parameters to select values that lead to the ‘best’ results, for which valid result interpretation requires adjustment for the inherent data-dredging or selective inference. The choice of different *β* and *k* values, however, has an effect on method performance. Figure S31 shows the results of the full analysis of the UK Biobank application study. For the weighted p-value approach, the default *β* = 2 led to the highest number of *New Discoveries*, but at the same time it resulted in the lowest *Recall* rate. For the sFDR method, *k* = 10 or 20 lead to an increased number of *New Discoveries* as compared with the default *k* = 2, at the cost of slightly reduced *Recall* rates. Thus, unlike the previous linkage and GWAS integration setting, the default value of *k* = 2 for sFDR appears to be sub-optimal for integrating functional meta-score with GWAS.

Overall, results of all three methods examined, weighted p-value, sFDR and FINDOR, were largely consistent with each other as detailed in Figure S23, despite of their apparent differences in methodological approach: FINDOR uses multiple individual functional scores and adjusts for linkage disequilibrium, while weight p-value and sFDR use meta-scores without LD adjustment; FINDOR and weighted p-value methods use the scores as weights, while sFDR only uses the scores to stratify SNPs into different priority groups. Overall, FINDOR and weighted p-value behaved more similarly with each other than with sFDR, where FINDOR and weighted p-value led to more new discoveries in the UK Biobank application, while sFDR is the most robust with the highest recall and precision rates in both the simulation and application studies.

In contrast to FINDOR, the weights used by the weighted p-value approach and the variant stratification in sFDR depend solely on the external meta-scores (i.e. independent of the GWAS summary statistics). This is to reduce the potential over-fitting which, though not severe for FINDOR, was observed in our simulation studies where the empirical family-wise error rate of FINOR was 0.0537, estimated from 50,000 simulation replicates; the empirical FWERs of weighted p-value and sFDR were 0.0501 and 0.0474, respectively. Interestingly, the weighted p-value approach used here performs similarly as compared with FINDOR in terms of both *New Discoveries* (Figure 5) and *Recall* (Figure 4). This, and combined with the FWER control results in Tables 1 and S1, suggests that using GWAS p-values and LD-regression to derive weights for weighted p-value method is not necessary. Thus, with the consideration of ease-of-implementation, we recommend weighted p-value and sFDR for practical applications.

However, the overall number of novel loci discovered after data-integration is limited regardless of the methods used; see Figures 3 and S28 for the contrast of the numbers of significant loci before and after data-integration for all the 1,132 UK Biobank traits analyzed. Notably, 162 traits (89%) in the nonsig trait category had no new discoveries in terms of independent loci by any of the three methods; “many of the non-significant results likely reflect limited statistical power rather than a true lack of heritability” as noted by Nealelab. Thus, results from this study show the limitation of the current sample size available for GWAS of complex traits, the informativeness of the current functional annotation measures, or power of the current data-integration methods that integrate GWAS summary statistics with functional annotations.

## Acknowledgment

This research is funded by the Natural Sciences and Engineering Research Council of Canada (NSERC, RGPIN-04934 and RGPAS-522594 to LS), the Canadian Institutes of Health Research (CIHR, MOP-310732 to LS), and the University of Toronto McLaughlin Centre Accelerator Grants in Genomic Medicine (MC-2019-15 to LS and ADP).

## Web Resources

UK Biobank, https://www.ukbiobank.ac.uk

UK Biobank GWAS summary statistics from Nealelab, http://www.nealelab.is/uk-biobank

UK Biobank GWAS QC steps performed by Nealelab, https://github.com/Nealelab/UK_Biobank_GWAS#imputed-v3-variant-qc

UK Biobank SNP-heritbability estimates from Nealelab, https://nealelab.github.io/UKBB_ldsc/

The 1,000 Genome Projects, http://tcag.ca/tools/1000genomes.html

CADD(v1.6), https://cadd.gs.washington.edu

Eigen (v1.0) through ANNOVAR software, http://annovar.openbioinformatics.org/en/latest/user-guide/filter/#eigen-score-annotations

FINDOR, https://github.com/gkichaev/FINDOR

## Data and Code Availability

Data used in this work are GWAS summary statistics and functional annotation scores which are publicly available; see Web Resources. All codes used for data analyses and simulation studies are open-resource and available at https://github.com/jianhuig/Integrate-gwas.

## Supplementary Information

**Figure S1:**
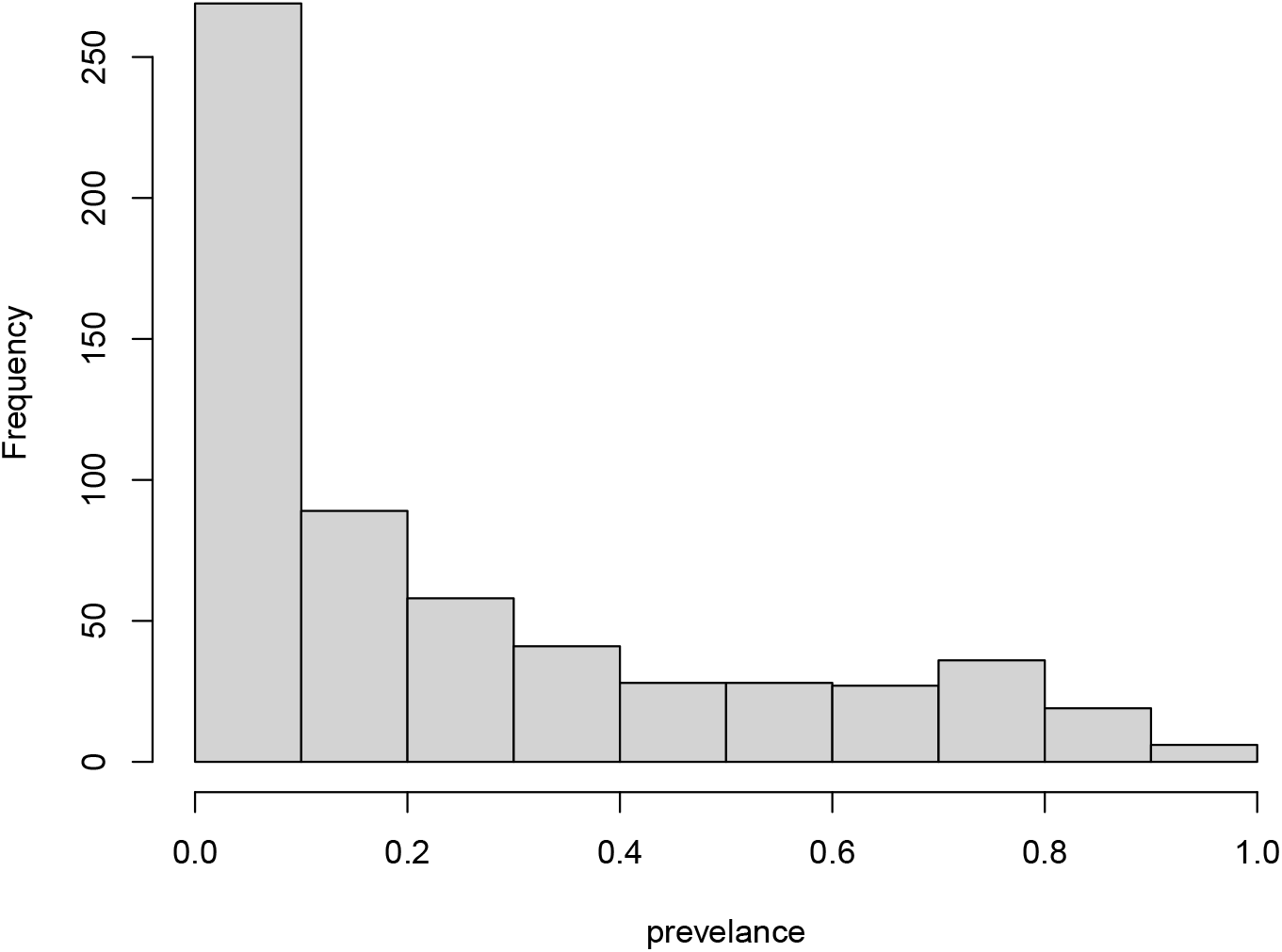
Histogram of prevalence for the 601 binary traits from the 1,132 UK Biobank traits analyzed in the application study.

**Figure S2:**
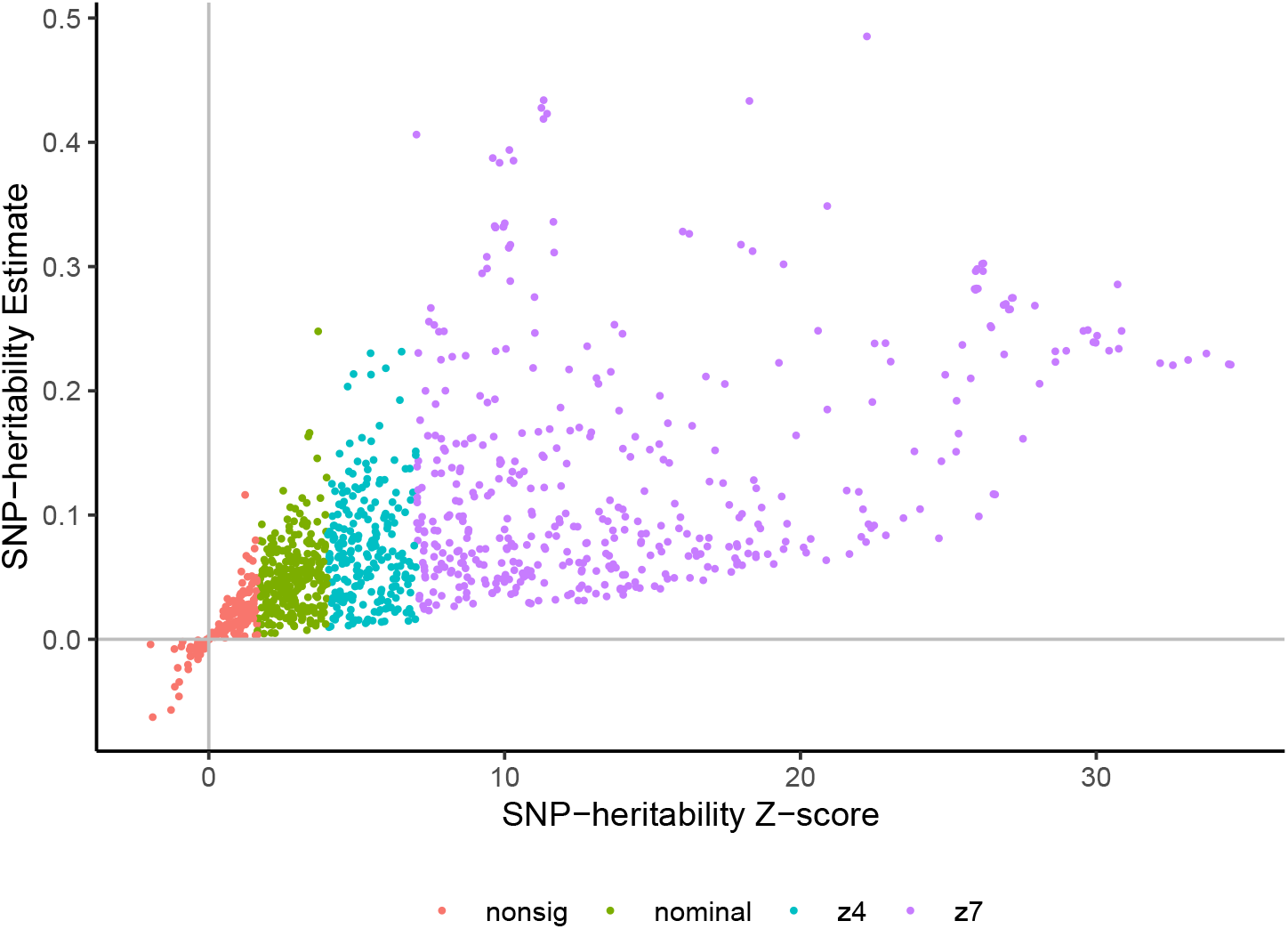
Contrast between SNP-heritability estimates and their z-scores for the 1,132 complex traits, based on the UK Biobank data analyzed by Nealelab. The 1,132 traits are categorized by Nealelab into four categories: nonsig (182 traits; *z* < 1.96), nominal (277 traits; 1.96 ≤ *z* < 4), z4 (235 traits; 4 ≤ *z* < 7), and z7 (438 traits; *z* ≥ 7).

### Inverse Normal Transformation

Unlike z score in simulations, however, phred score are all positive and only have relative meaning: higher value means that a variant is more likely to be deleterious. Since inverse variance meta-analysis and Fisher’s method need z scores or p-values to pool information, phred score need to be transformed so that small scores (close to 0) have large p-values. Without loss of generality, let *e_i_* be the Eigen phred score, we perform rank based inverse normal transformation by

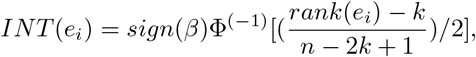

where Φ^−1^ is the inverse normal, rank(*e_i_*) is the rank of *e_i_* in descending order and *k* = 3/8 is the Blom offset (Blom, 1958). Because meta-analysis is directional specific, we manually fix the sign of transformed Eigen score to be consistent with sign of original GWAS effect *β_i_*, which is the best scenario for meta-analysis.

**Figure S3:**
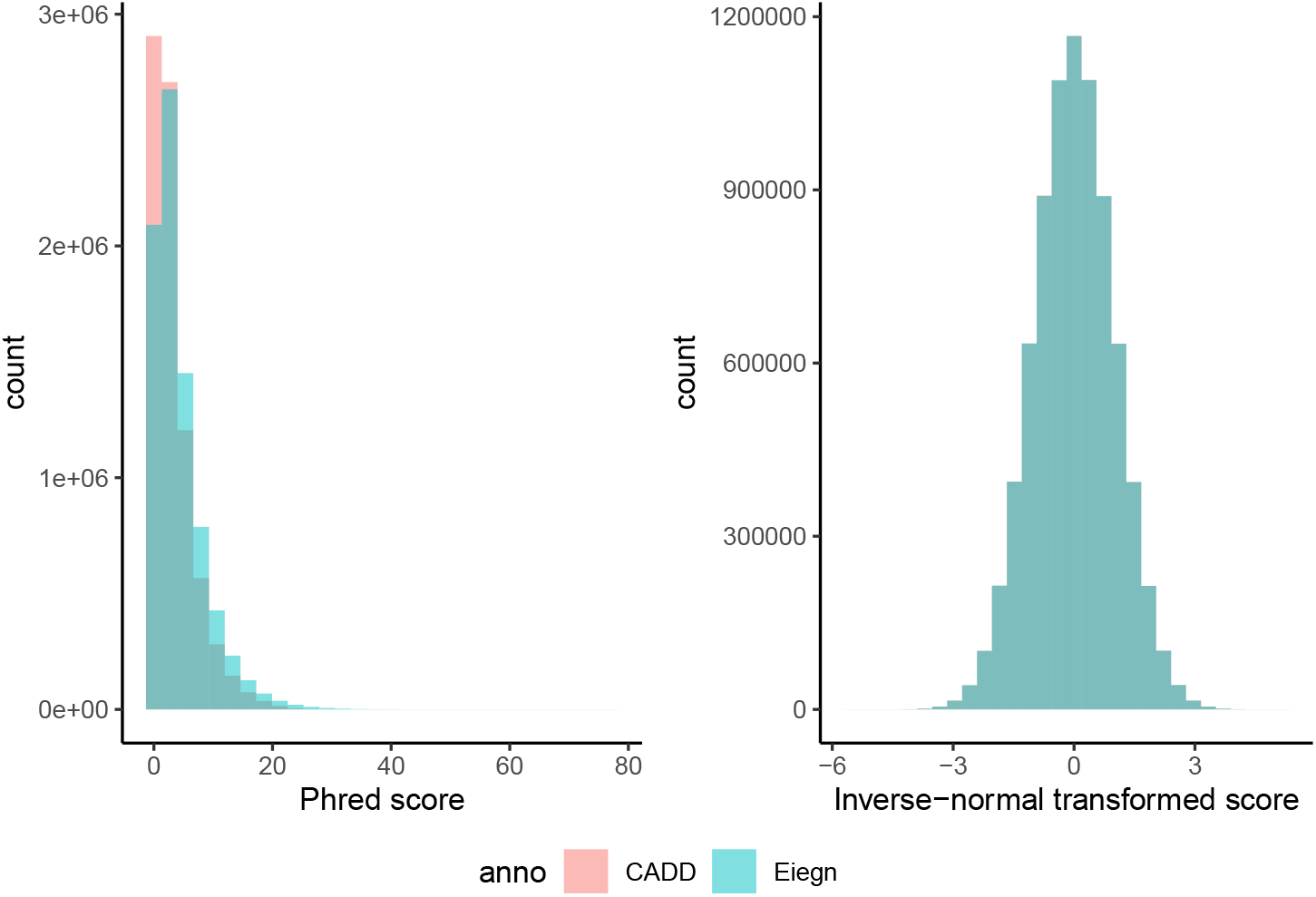
Distributions of CADD and Eigen functional meta-scores. The left figure shows the phred-scaled meta-scores, −10 log_10_(ranks of the raw scores/total number SNPs), and right figure shows the inverse-normal transformed meta-scores.

**Figure S4:**
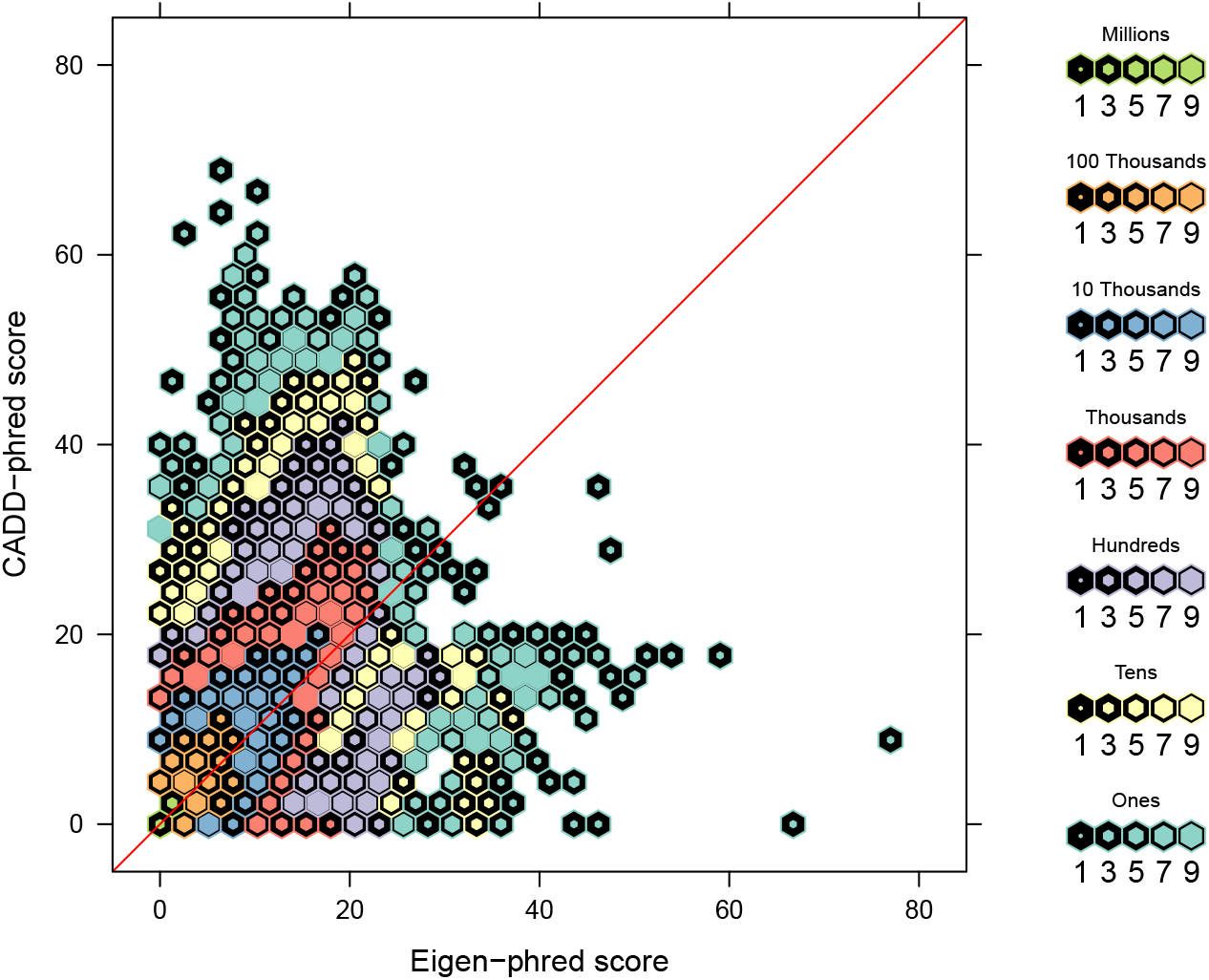
Contrast between CADD and Eigen functional meta-scores on the phred-scale. The phred scores are −10 log_10_(ranks of the raw scores/total number SNPs). The color and inner size of each hexagon represents the number of SNP counts.

**Figure S5:**
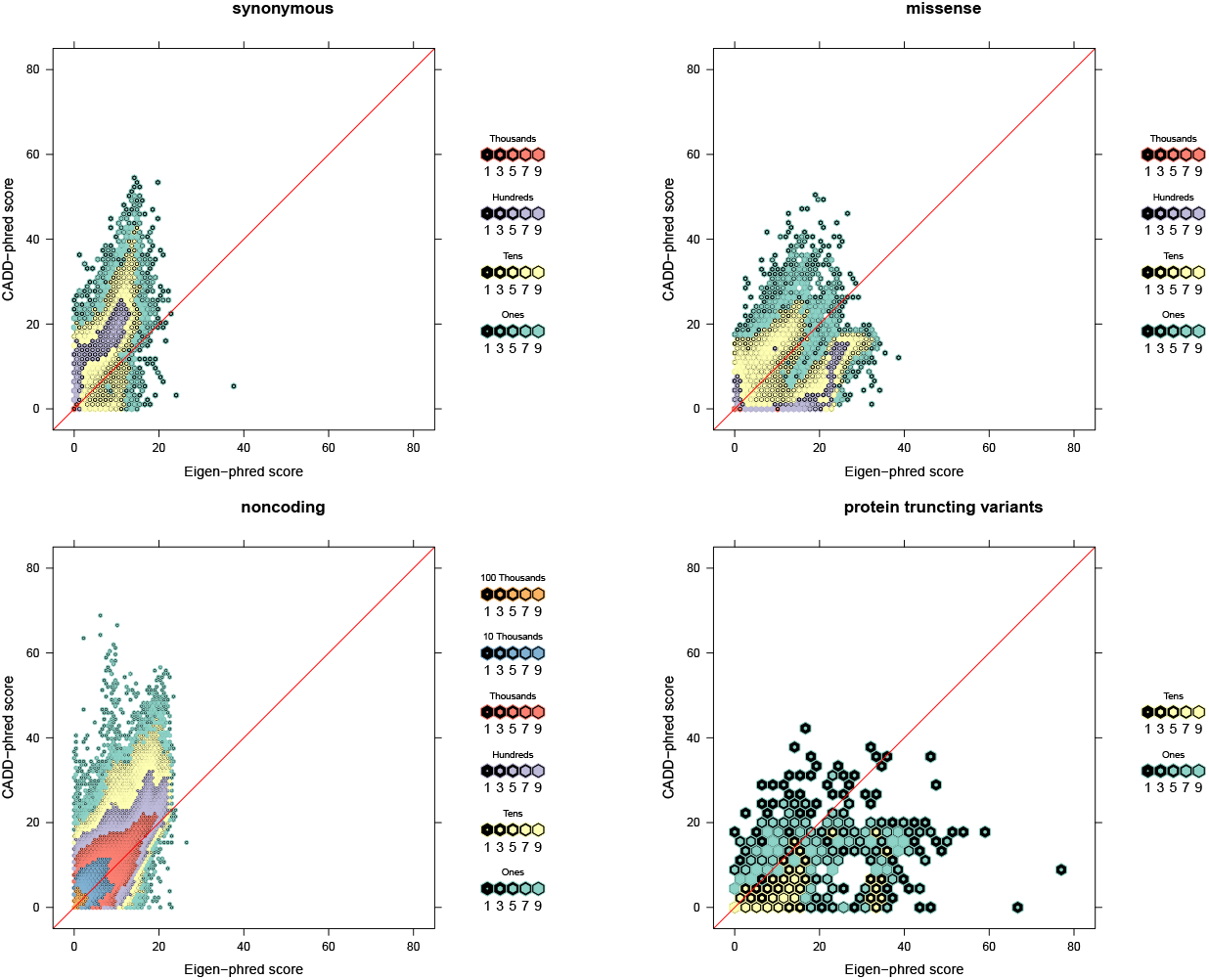
Contrast between CADD and Eigen functional meta-scores on the phred-scale. The phred scores are −10 log_10_(ranks of the raw scores/total number SNPs), stratified by four variant consequence categories: missense, non-coding, synonymous, and protein truncating variants. The color and inner size of each hexagon represents the number of SNP counts.

### Detailed simulation design

We first simulated GWAS p-value under the null on 1000 Genomes Project variants. We then applied FINDOR, weighted p-value, and sFDR on simulated p-values. The traditional FWER is defined as the proportion of replicates that find at least one false positive.

#### Genotype Data Description

- We obtained 1000 Genome Project genotypes from The Center of Applied Genomics at Sickkids (http://tcag.ca/tools/1000genomes.html).
- The data went through extensive data cleaning (http://tcag.ca/documents/tools/omni25_qcReport.pdf).
- In addition, we restricted our analysis to SNPs with MAF > 0.05.
- The dataset analyzed contains 1,756 independent samples and 422,923 bi-allelic SNPs.
- The proportion of non-missing genotypes was calculated per SNP. The call rate was excellent: 422,364 of 422,923 (99.7%) of the SNPs had call rate > 97%. We fast impute missing genotypes by randomly sampling acoording to allele frequencies.

#### Simulation Design

- We simulated a continuous trait for each sample using *Y* ~ *N*(0, 1), independent of the genotype data.
- We conducted a basic GWAS without adjusting any covariates. Because Y was randomly simulated and the population is homogenous, PCA adjustment is not critical; indeed, the GWAS p-values were Unif(0,1) distributed.
- We then run ldsc (https://github.com/bulik/ldsc) with reference *1000G_Phase3_baselineLD_ldscores.tgz*, frq *1000G_Phase3_frq.tgz* and weights *1000G_Phase3_weights_hm3_no_MHC.tgz* from Alke’s Group (https://alkesgroup.broadinstitute.org/LDSCORE/). In total 100 equal sized bins are created based on quantities of *E*(*χ*^2^).
- For FINDOR, we use weight 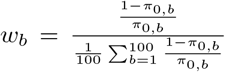, where *π*_0,*b*_ is the estimated proportion of null hypothesis in each bin using cubic spline proposed by Storey and Tibshirani. We reject null hypothesis if 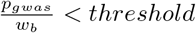.
- For weighted p-value and sFDR, we obtain CADD (v1.6), and using same methods we described under methods section in the paper.
- Finally, we repeat the simulation, independently, 50,000 times.

#### Results

**Figure S6:**
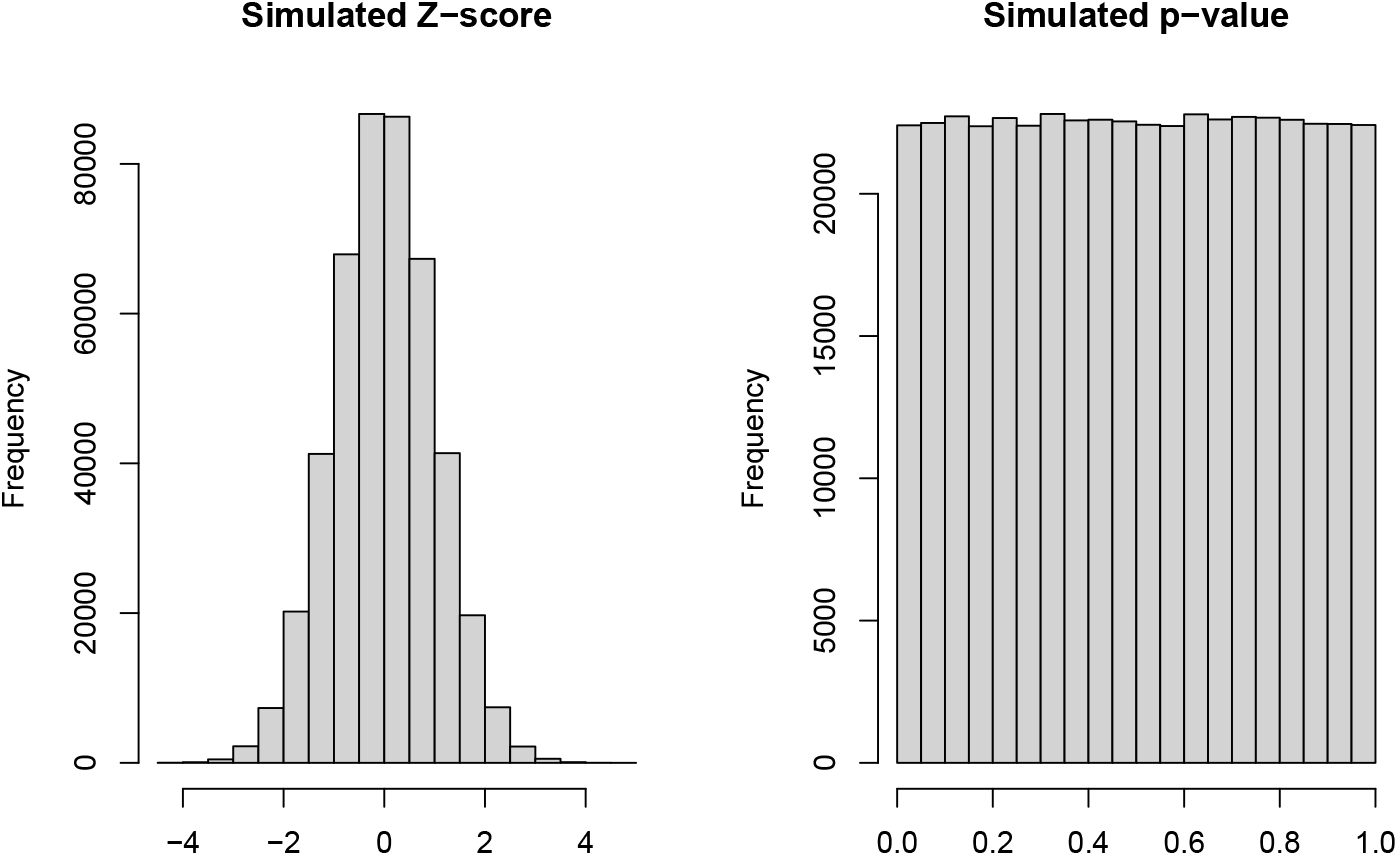
Histograms of GWAS Z values and p-values from the simulation study design III. A *N*(0, 1) distributed trait was simulated for 1,756 individuals from the 1000 Genomes Project, *independently* of the genotypes of 422,923 bi-allelic, autosomal and common (MAF > 5%) SNPs. The GWAS summary statistics for the 422,923 SNPs were obtained by regressing the trait values of the 1,756 individuals on the additively coded genotypes. Results shown here are from one simulation run, randomly selected from a total of 50,000 simulation runs.

- Table S1 shows the number of simulation replicates, out of a total of *k* = 50, 000, in which 0, 1, 2, or 3 SNPs were declared significant; no replicates had more than 3 false positive SNPs. Let *k_n_* represent number of replicates have *n* false findings. The sample estimate of (*k* − *k*_0_)/*k* = (*k*_1_ + *k*_2_ + *k*_3_)/*k* provides an estimate of the FWER for each of the method. Note that in this case where all SNPs are under the null, the estimates are also FDR estimates.
- Because the total number of SNPs in our dataset are around half million, we have applied the Beefaroni threshold 0.05/422923 = 1.2*e* − 7 instead of 5*e* − 8. Table S1 shows the results.

#### Conclusion and Discussion

We noticed an increased Type I error in FINDOR method than baseline method in both cases.

**Table S1.** Results of simulation study design I that leverages the real functional scores combined with GWAS summary statistics simulated under the null of no association. Breakdown of the number of false positive findings for the *k* = 50, 000 simulated replicates using the Bonferroni threshold of 1.2 × 10^−7^. Among the *k* = 50, 000 replicates, *k_n_*, *n* = 0, …, 3 represent number of replicates have *n* false findings. Assuming the true FWER is 0.05, we expect the estimate obtained from the 50,000 independent simulation replicates to have a standard error of 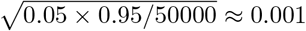. Thus, a method with a FWER estimate outside [0.047, 0.053] can be considered inaccurate.

| Method | $k_0$ | $k_1$ | $k_2$ | $k_3$ | FWER |
| --- | --- | --- | --- | --- | --- |
| Baseline | 47522 | 2409 | 68 | 1 | 0.04956 |
| Meta-analysis | 47617 | 2332 | 49 | 2 | 0.04766 |
| Fisher's method | 48172 | 1790 | 38 | 0 | 0.03656 |
| Weighted p-value | 47495 | 2443 | 62 | 0 | 0.05010 |
| stratified FDR | 47629 | 2303 | 67 | 1 | 0.04742 |
| FINDOR | 47316 | 2618 | 64 | 2 | 0.05368 |

Using CADD as an example, let *CADD_i_* and *CADD_j_* be the annotation scores of SNPs *i* and *j*, respectively. We first defined a pair-wise similarity measure as 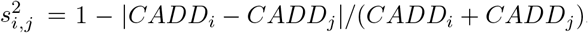, which is bounded between 0 and 1, where 1 means two scores are identical whereas a value close to 0 suggests a lack of similarity. We then contrast 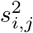 with 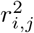, the traditional LD measure of genotype similarity between two SNPs. Results in Figure S7 show that there are no clear concordance between the two measures. A closer examination of a few randomly selected regions led to the same conclusion (Figure S8); see Supplementary Information for details.

**Figure S7:**
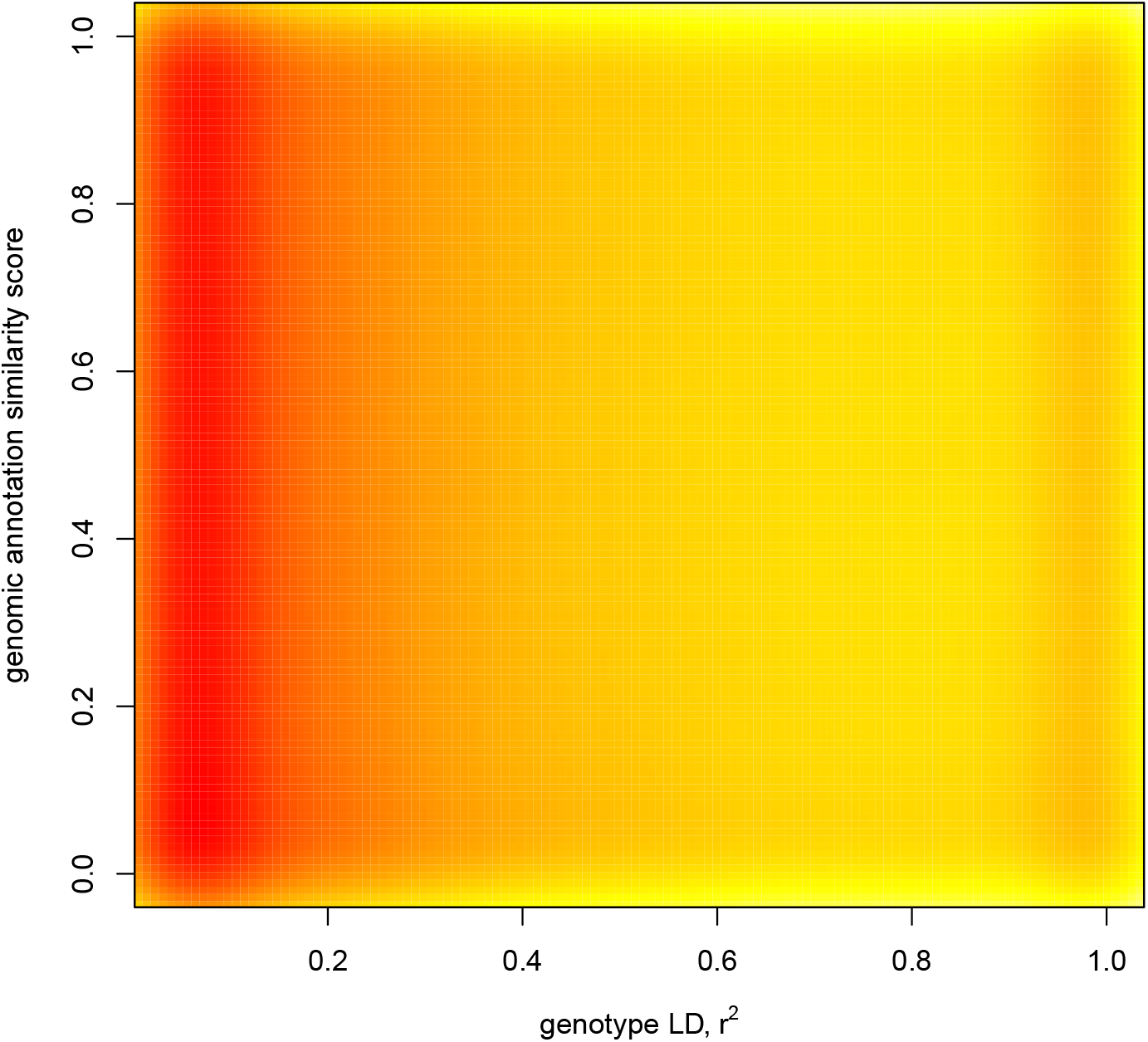
Contrast between 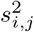, a pair-wise similarity measure of annotation scores of SNPs *i* and *j*, and 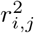, the linkage disequilibrium measure of genotype correlation, for the common bi-allelic autosomal SNPs of the UK Biobank data. Annotation similarity measure is defined as 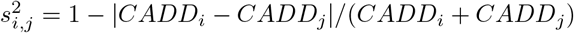, which is bounded between 0 and 1, where 1 means two scores are identical whereas a value close to 0 suggests a lack of similarity. The pair-wise LD 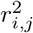 values were calculated for all possible pairs within 1*MB* on each chromosome using emeraLD (Quick et al., 2019). Area with darker color (red) represents higher number of points are observed in that region.

**Figure S8:**
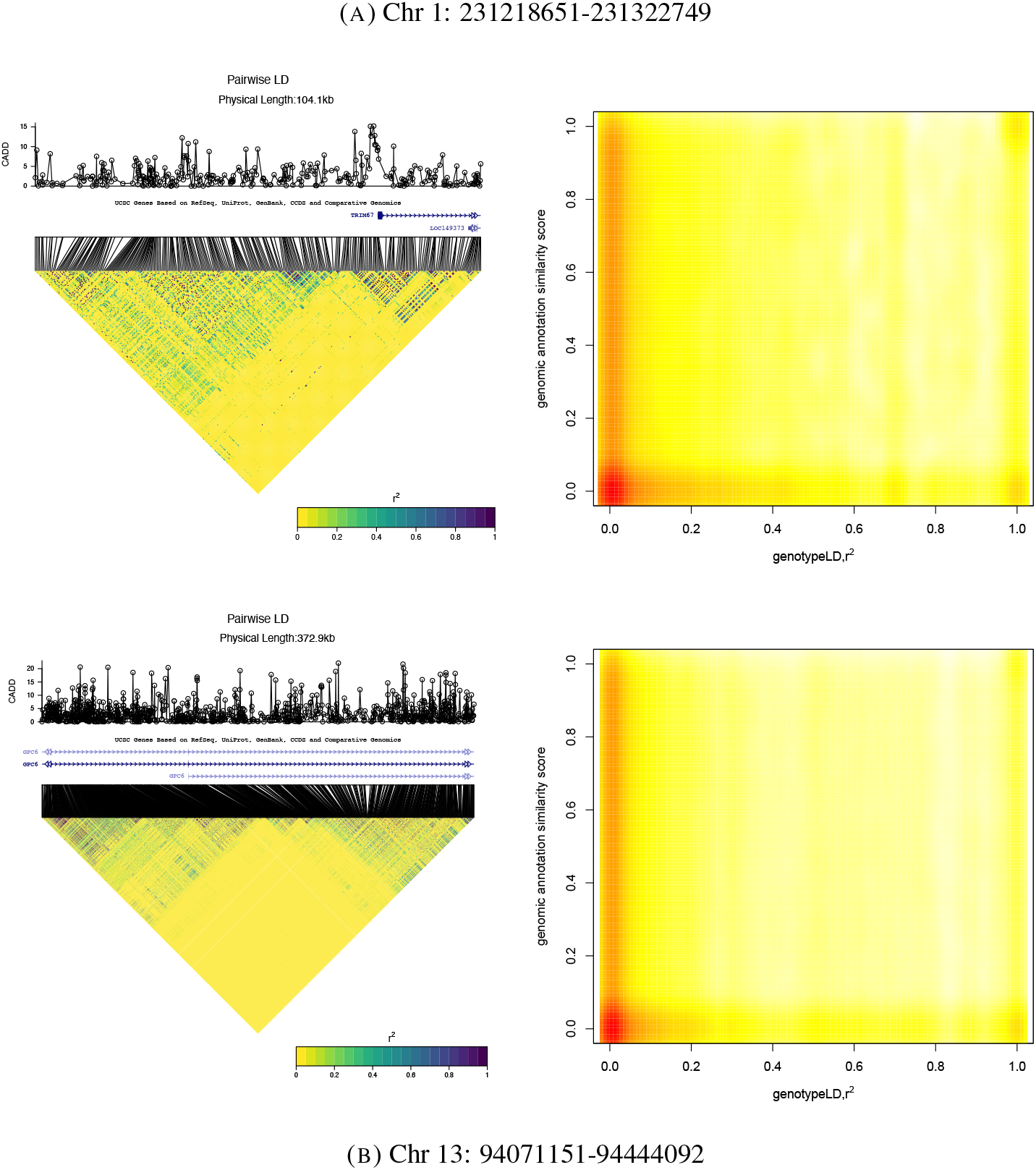
For two randomly selected regions, Left: the CADD scores and the standard pair-wise linkage disequilibrium (LD) plot along with gene information. Right: contrast between 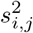, a pair-wise similarity measure of annotation scores of SNPs *i* and *j*, and 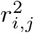, the LD measure of genotype correlation. Annotation similarity measure is defined as 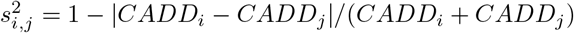, which is bounded between 0 and 1, where 1 means two scores are identical whereas a value close to 0 suggests a lack of similarity. Area with darker color (red) represents higher number of points are observed in that region.

**Figure S9:**
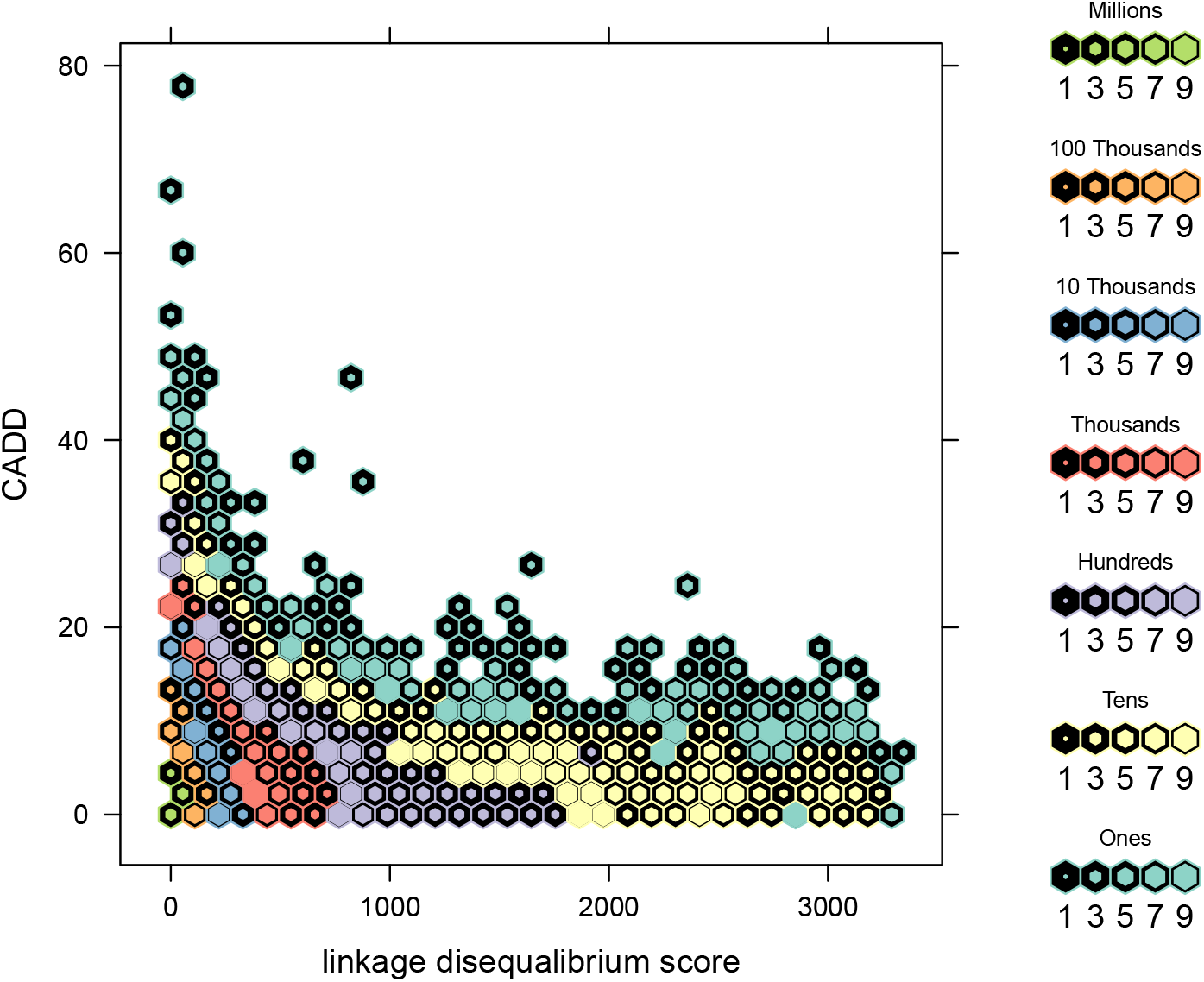
Contrast between CADD meta-score and LD score for the 7,895,174 common, bi-allelic autosomal SNPs of the UK Biobank data. For each variant *i*, the plot shows its CADD meta-score and the sum of 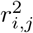 across all other *j* variants analyzed. The color and inner size of each hexagon represents the number of SNP counts.

**Figure S10:**
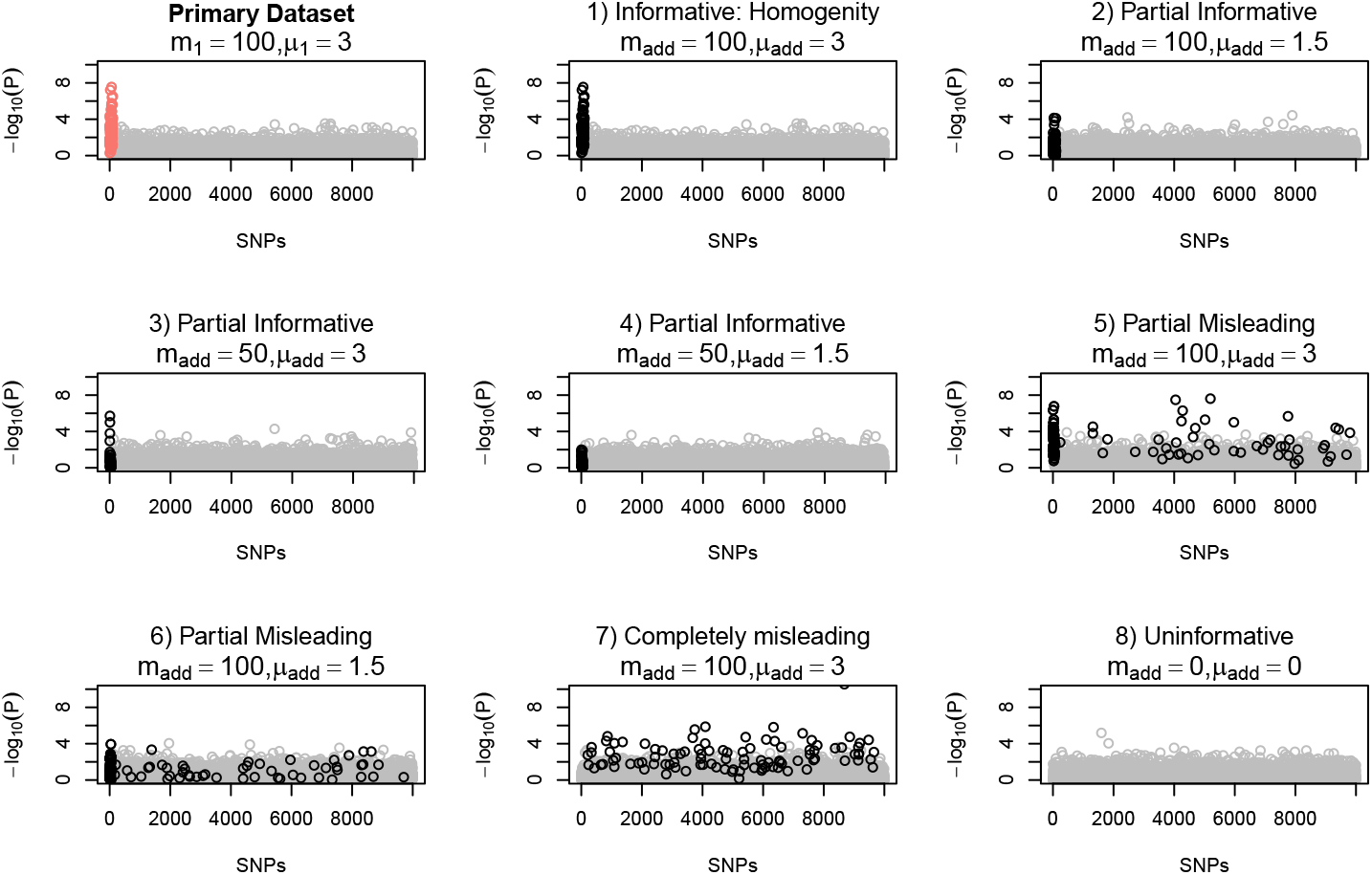
Illustration of Simulation study design III, varying the informativeness of genomic information. Among the total *m* = 10, 000 SNPs, the first *m*_1_ = 100 SNPs are truly associated (red circles) whose GWAS association summary statistics were drawn from *N*(*μ*_1_ = 3, 1); the remaining SNPs are not associated whose association summary statistics were drawn from *N*(0, 1). The top left figure shows the Manhattan plot of of one simulation run. The other figures demonstrate eight scenarios for the additional information available for data-integration. The summary statistics of the additional information available were drawn, independently, from *N*(*μ_add_*, 1) for *m_add_* SNPs (black circles) and *N*(0, 1) for the remaining SNPs; the locations of the *m_add_* SNPs may differ from those of the *m*_1_ associated SNPs. The eight scenarios fall into four categories. Category I is completely informative– (homogeneity): (1) *m_add_* = 100, *μ_add_* = 3 and locations of the *m_add_* SNPs perfectly match those of *m*_1_ GWAS truly associated SNPs. Category II is partially informative: (2) *m_add_* = 100 and *μ_add_* = 1.5; (3) *m_add_* = 50 and *μ_add_* = 3; (4) *m_add_* = 50, *μ_add_* = 1.5, and all *m_add_* SNPs coincide with (some of) the *m*_1_ SNPs. Category III is (partially or completely) misleading: (5) *m_add_* = 100 and *μ_add_* = 3; (6) *m_add_* = 100 and *μ_add_* = 1.5, but in both scenarios only 50 out of the *m_add_* SNPs coincide with 50 of the *m*_1_ SNPs. And (7) *m_add_* = 100 and *μ_add_* = 3, but none of the *m_add_* SNPs coincide with the *m*_1_ SNPs. Category IV is uninformative: (8) *m_add_* = 0 and *μ_add_* = 0. That is, the additional information available is white noise.

### Permuted Functional Annotations

Here we integrated real UK Biobank GWAS summary statistics for the 1,132 traits with *permuted* CADD (and Eigen) meta-scores, using meta-analysis, Fisher’s method, the weighted p-value approach, and the stratified FDR control. The suitable performance measures here are *Recall_t_* = *TP_t_*/*m*_1,*t*_ and *Precision_t_* = 1 − *FDR_t_* = *TP_t_*/*P_t_*, where *m*_1,*t*_ was defined as the number of genome-wide significant GWAS findings prior to data-integration for trait *t*, and *P_t_* and *TP_t_* are the numbers of positives and true positives after data-integration. *Recall* rates were obtained for 723 GWAS with *m*_1,*t*_ > 0. Regardless if the original GWAS has a finding or not, the *Precision* rates were conservatively defined as one for 558, 505, 440, and 415 GWAS with *P_t_* = 0 after data integration using, respectively, meta-scores, Fisher’s method, weighted p-value, and sFDR.

In addition to the Figure 1, Figure S11 shows the results stratified by SNP-heritability estimates of GWAS; Figure S12 shows the rate calculated using SNPs instead of loci; Figure S13 shows the results by integrating *permuted* Eigen instead of CADD.

**Figure S11:**
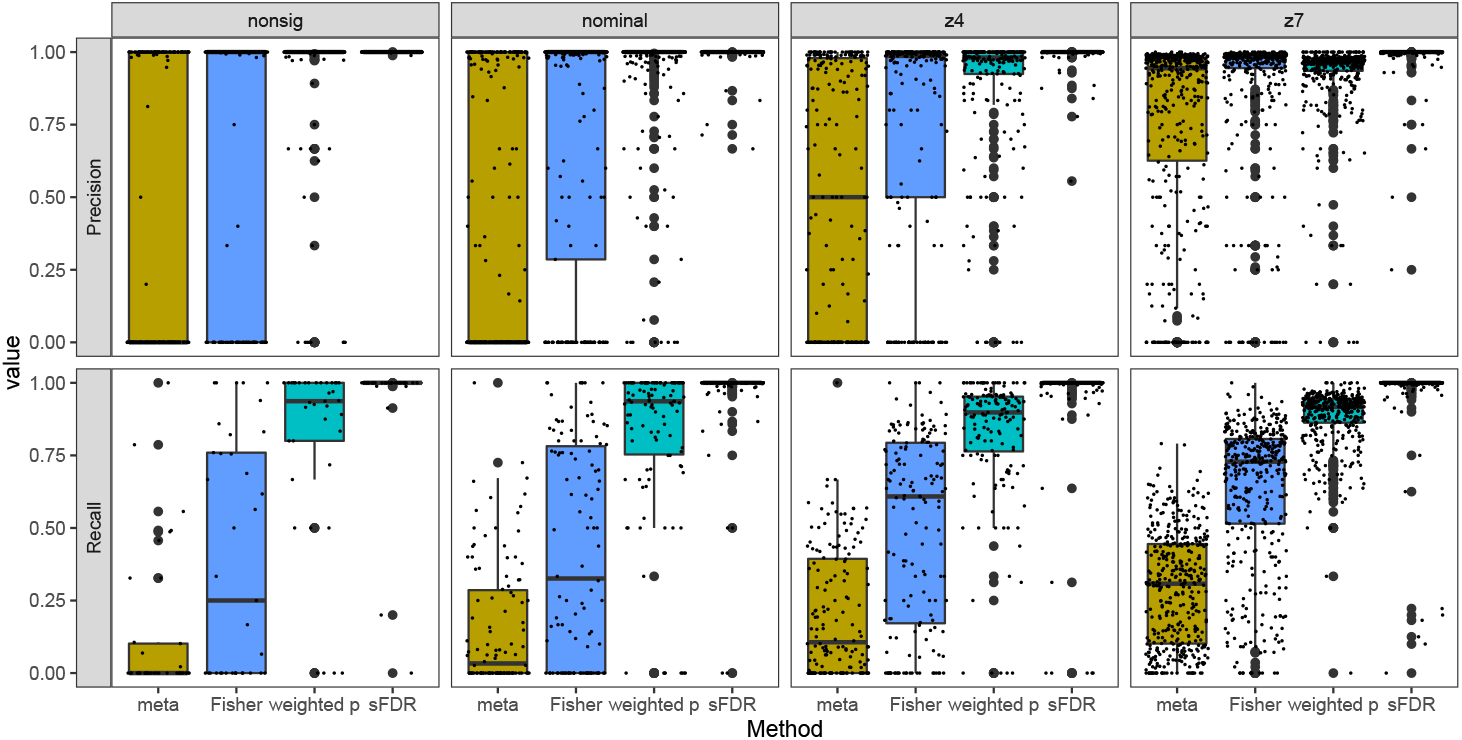
The trait-type-stratified *Recall* and *Precision* rates obtained from simulation study design II. The study integrated the 1,132 UK Biobank GWAS summary statistics with *permuted* CADD functional meta-scores, using meta-analysis, Fisher’s method, the weighted p-value approach, and the stratified FDR control. The 1,132 traits fall into four categories by Nealelab: nonsig (182 traits; *p* > 0.05), nominal (277 traits; *p* < 0.05), z4 (235 traits; *p* < 3.17 × 10^−5^), and z7 (438 traits; *p* < 1.28 × 10 × −12). *Recall_t_* = *TP_t_*/*m*_1,*t*_ and *Precision_t_* = 1 − *FDR_t_* = *TP_t_*/*P_t_*, where *m*_1,*t*_ is the number of genome-wide significant independent loci prior to data-integration for trait *t*, and *P_t_* and *TP_t_* are the numbers of positives and true positives after data-integration.

**Figure S12:**
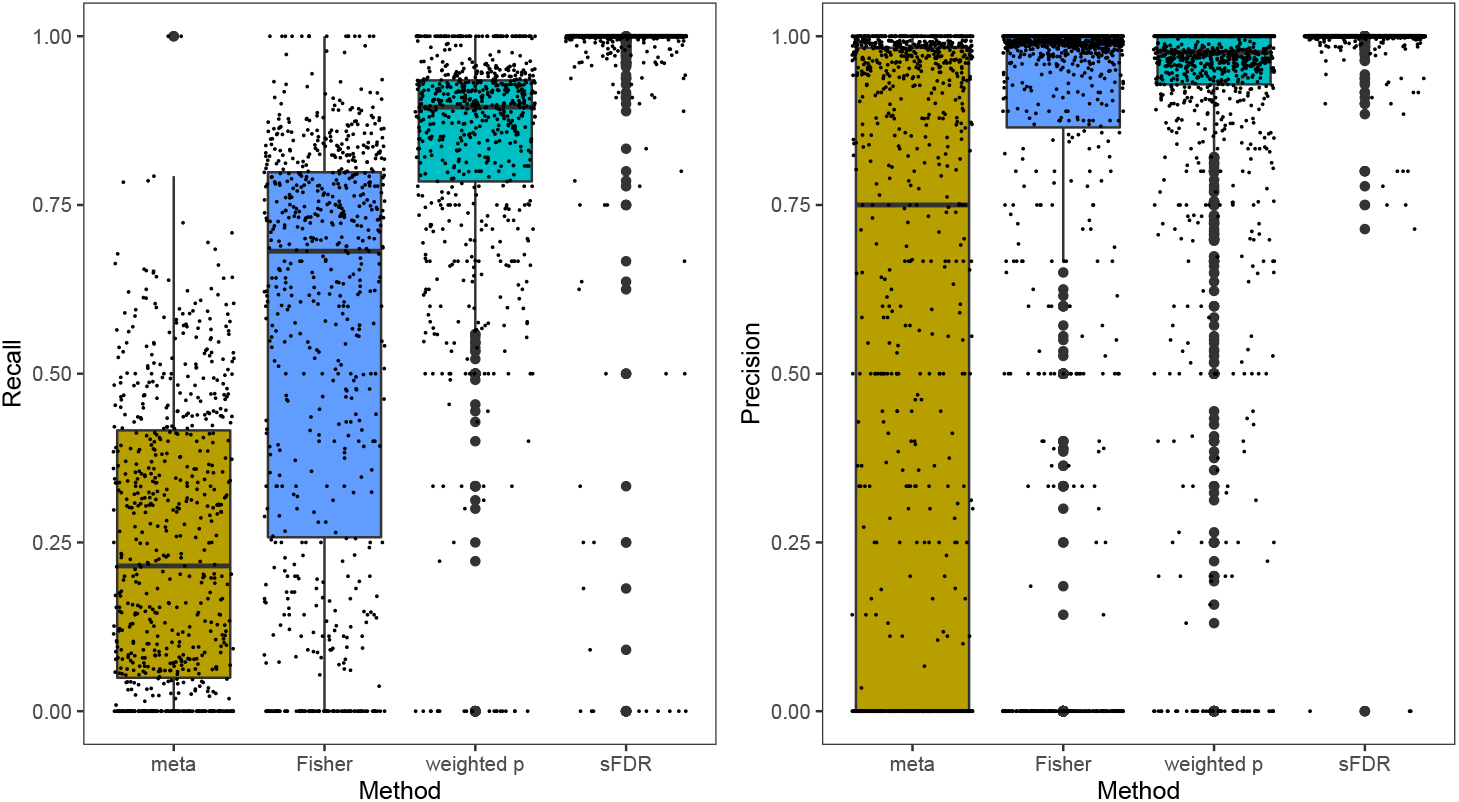
SNP results for the *Recall* and *Precision* rates obtained from simulation study design II. The study integrated the 1,132 UK Biobank GWAS summary statistics with *permuted* CADD functional meta-scores, using meta-analysis, Fisher’s method, the weighted p-value approach, and the stratified FDR control. *Recall_t_* = *TP_t_*/*m*_1,*t*_ and *Precision_t_* = 1 − *FDR_t_* = *TP_t_*/*P_t_*, where *m*_1,*t*_ is the number of genome-wide significant SNPs prior to data-integration for trait *t*, and *P_t_* and *TP_t_* are the numbers of positives and true positives after data-integration.

**Figure S13:**
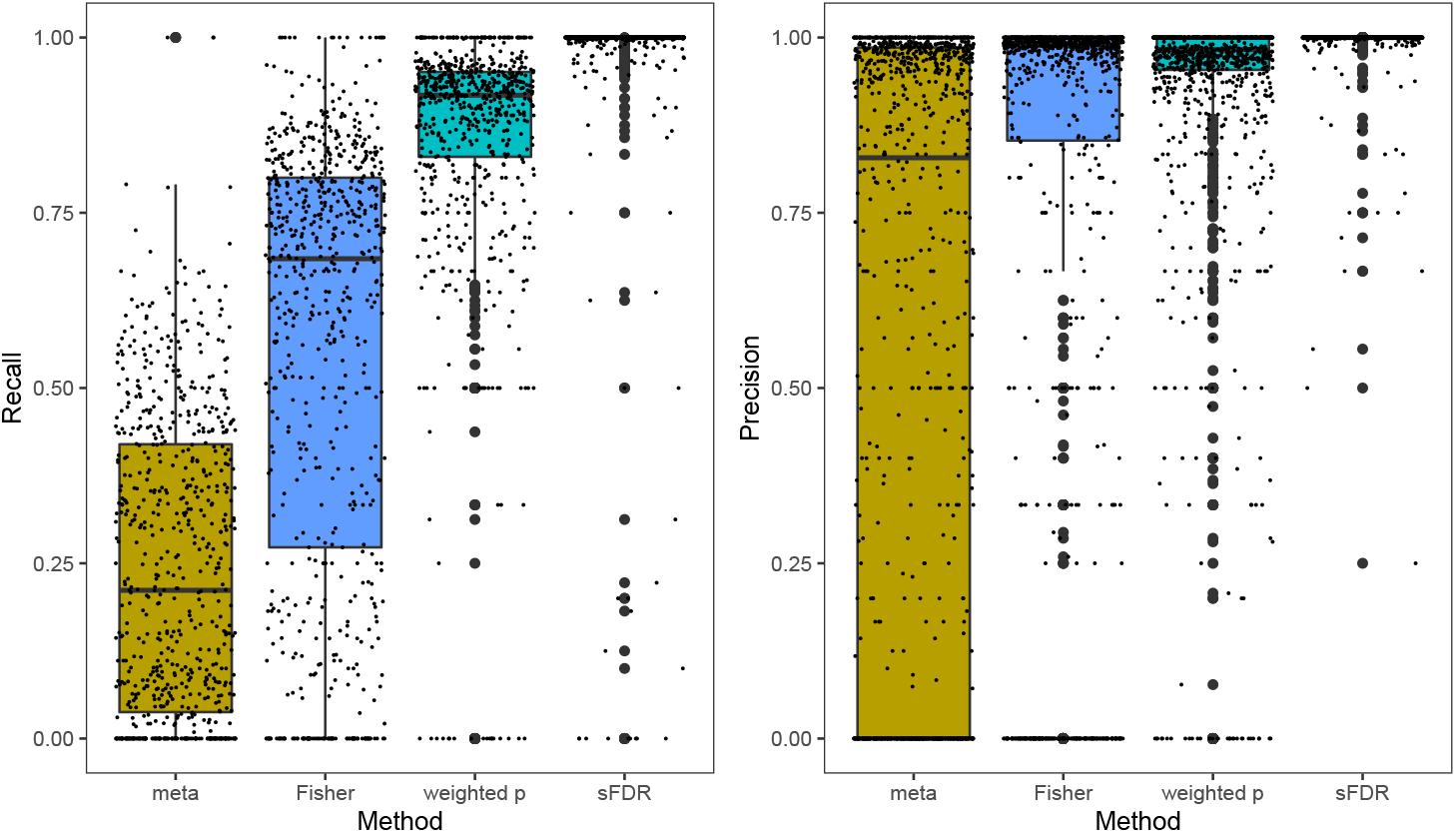
Eigen results for the *Recall* and *Precision* rates obtained from simulation study design II. The study integrated the 1,132 UK Biobank GWAS summary statistics with *permuted* Eigen functional meta-scores, using meta-analysis, Fisher’s method, the weighted p-value approach, and the stratified FDR control. *Recall_t_* = *TP_t_*/*m*_1,*t*_ and *Precision_t_* = 1 − *FDR_t_* = *TP_t_*/*P_t_*, where *m*_1,*t*_ is the number of genome-wide significant independent loci prior to data-integration for trait *t*, and *P_t_* and *TP_t_* are the numbers of positives and true positives after data-integration.

**Figure S14:**
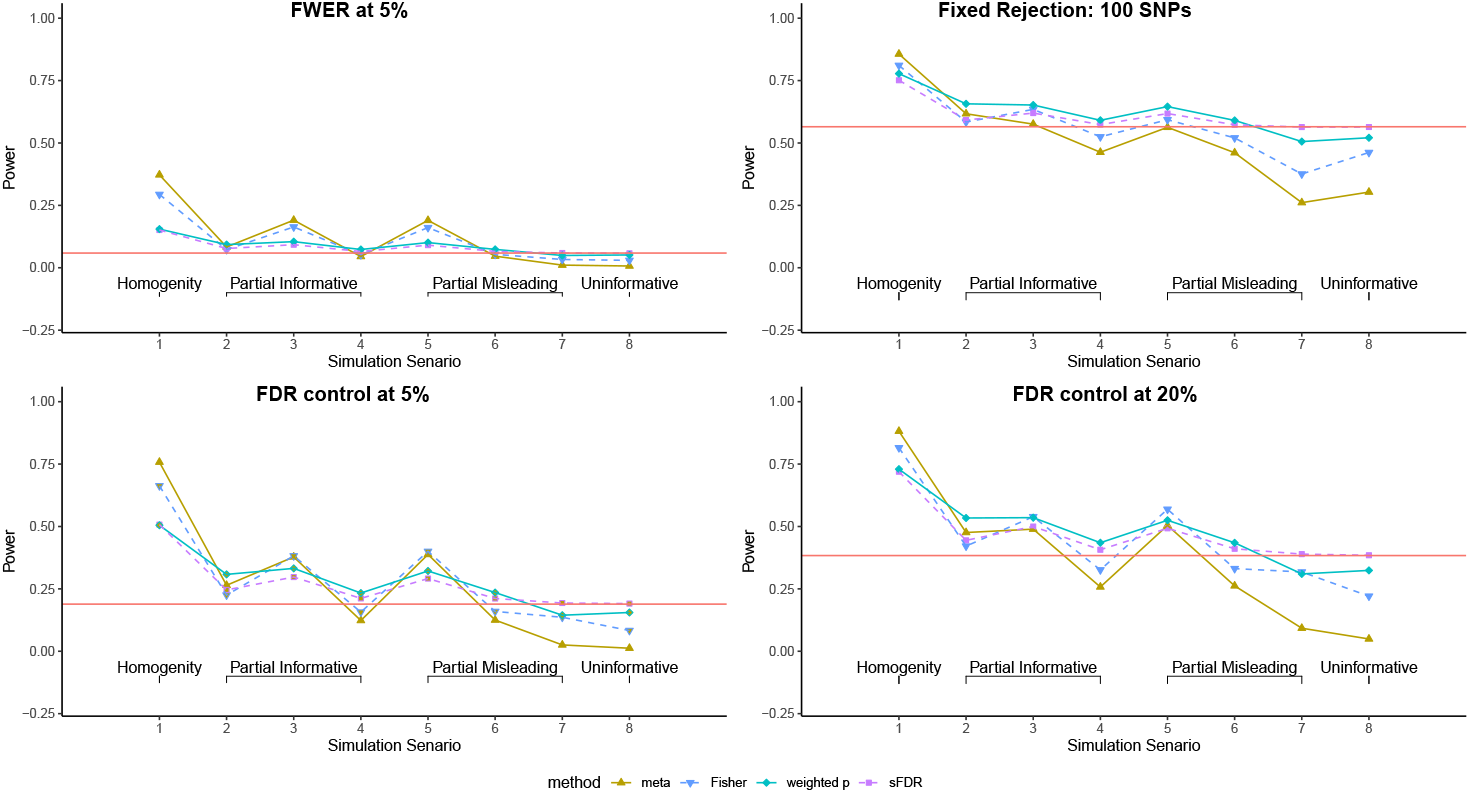
Power obtained from simulation study design III. The study integrated simulated GWAS summary statistics with simulated additional information with varying degrees of informativeness, using meta-analysis, Fisher’s method, the weighted p-value approach, and the stratified FDR control. There are 10,000 independent SNPs among which 100 are truly associated whose summary statistics were drawn from *N*(3,1); the rest from *N*(0, 1). For the additional information available for data integration, the details of the eight simulation scenarios are provided in the text and illustrated in Figure S10. Power is the proportion of true signals detected after data integration, estimated from 1,000 simulation replicates, using four different decision rules, controlling family-wise error rate at 0.05, rejecting top 100 ranked SNPs, and controlling false discovery rate at 5% or 20%. The red line represents baseline power of using GWAS data alone without data-integration.

**Figure S15:**
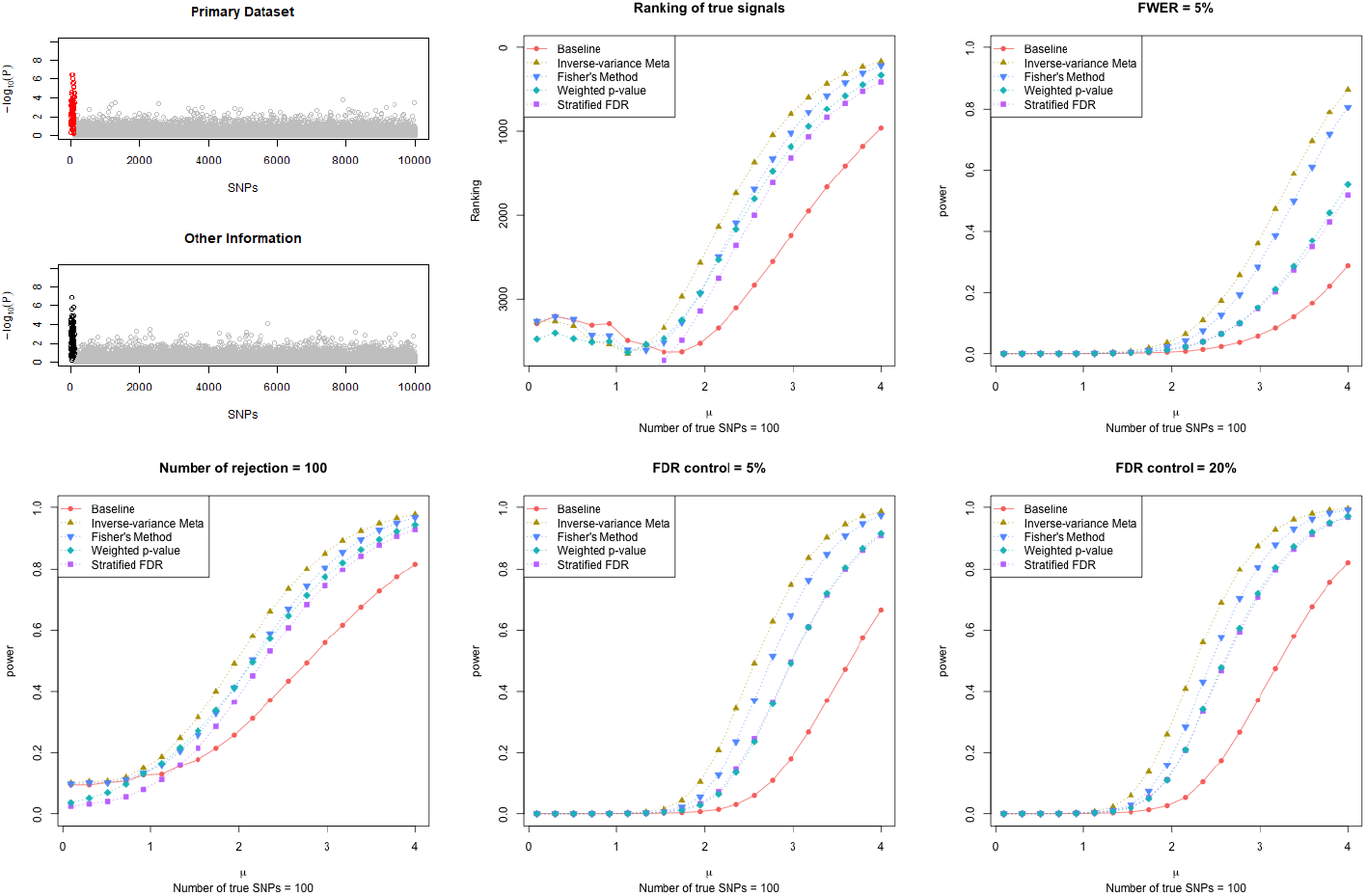
Results of simulation study design III for scenario (1) in Figure S10 from Category I type of informativeness, with the true effect size *μ*_1_ for the *m*_1_ causal SNPs varying from 0.1 to 4. Completely Informative (Homogeneity): *m_add_* = 100, *μ_add_* = *μ*_1_ and locations of the *m_add_* SNPs perfectly match those of *m*_1_ GWAS truly associated SNPs. We assumed the total number of SNPs *m* = 10, 000, among which the first *m*_1_ = 100 SNPs (in red on top-left panel) are truly associated. The corresponding summary statistics *z_i_*’s were drawn, independently, from *N*(*μ*_1_,1) for the *m*_1_ associated SNPs, and from *N*(0, 1) for the remaining null SNPs. We then assumed *z_i,add_*’s (in black on top-left panel) as the additional information available, which were drawn, independently, from *N*(*μ_add_*, 1) for *m_add_* SNPs and *N*(0, 1) for the remaining SNPs.

**Figure S16:**
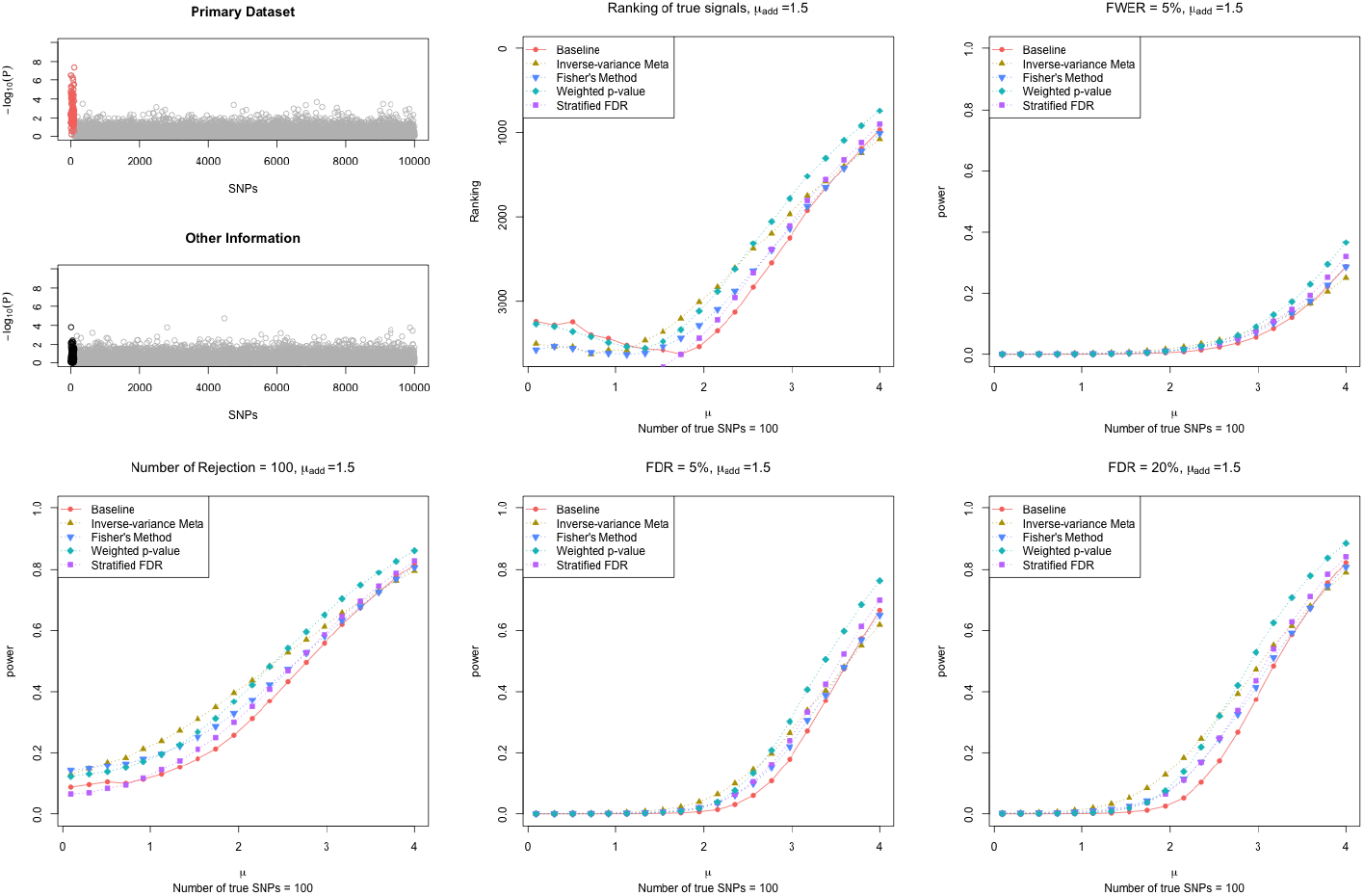
Results of simulation study design III for scenario (2) in Figure S10 from Category II type of informativeness, with the true effect size *μ*_1_ for the *m*_1_ causal SNPs varying from 0.1 to 4. Partially Informative: *m_add_* = 100, *μ_add_* = 1.5, and all *m_add_* SNPs coincide with (some of) the *m*_1_ SNPs. We assumed the total number of SNPs *m* = 10, 000, among which the first *m*_1_ = 100 SNPs (in red on top-left panel) are truly associated. The corresponding summary statistics zi’s were drawn, independently, from *N*(*μ*_1_,1) for the *m*_1_ associated SNPs, and from *N*(0, 1) for the remaining null SNPs. We then assumed *z_i,add_*’s (in black on top-left panel) as the additional information available, which were drawn, independently, from *N*(*μ_add_*, 1) for *m_add_* SNPs and *N*(0, 1) for the remaining SNPs.

**Figure S17:**
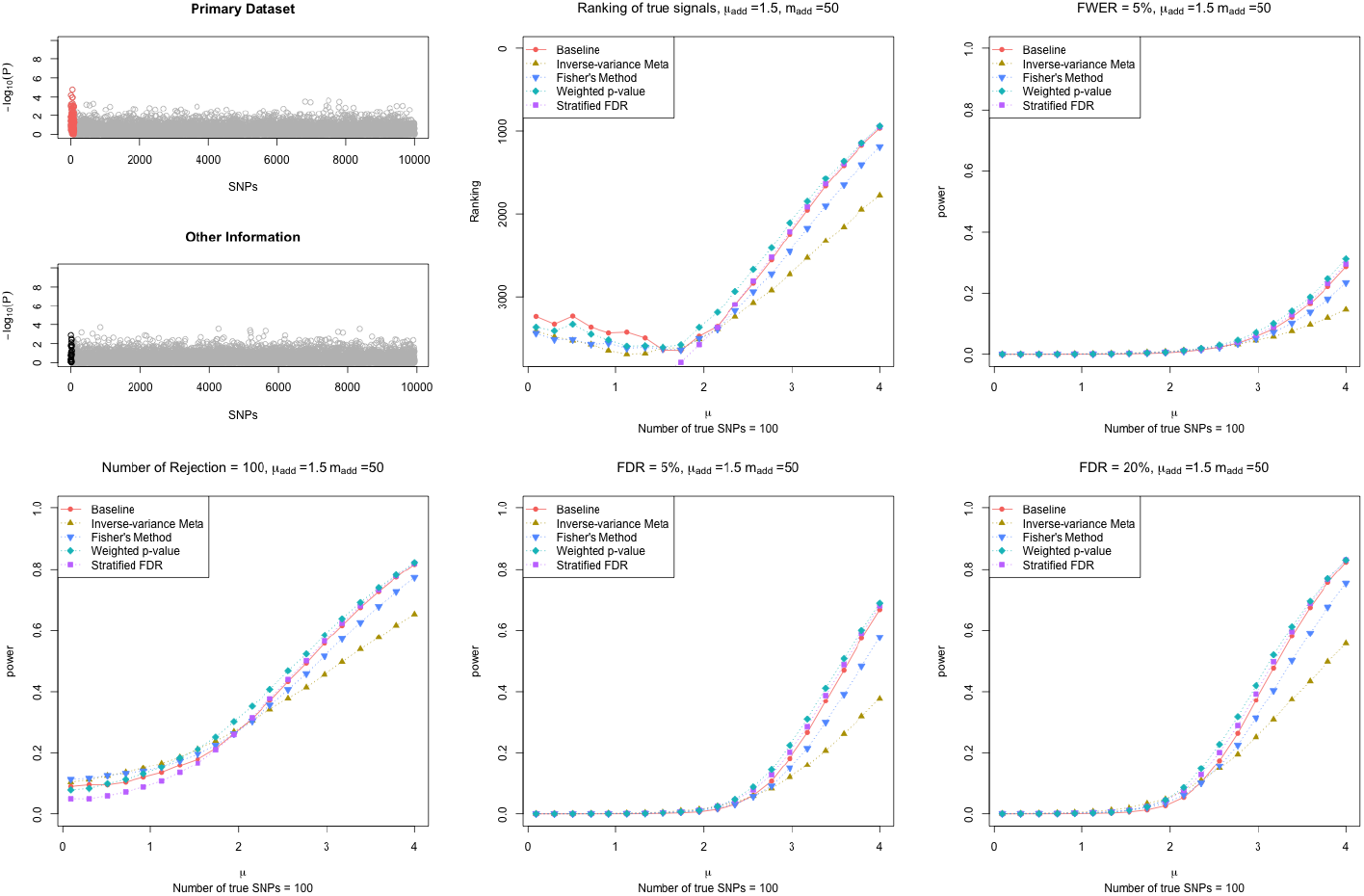
Results of simulation study design III for scenario (4) in Figure S10 from Category II type of informativeness, with the true effect size *μ*_1_ for the *m*_1_ causal SNPs varying from 0.1 to 4. Partially Informative: *m_add_* = 50, *μ_add_* = 1.5, and all *m_add_* SNPs coincide with (some of) the *m*_1_ SNPs. We assumed the total number of SNPs *m* = 10, 000, among which the first *m*_1_ = 100 SNPs (in red on top-left panel) are truly associated. The corresponding summary statistics ¾’s were drawn, independently, from *N*(*μ*_1_,1) for the *m*_1_ associated SNPs, and from *N*(0, 1) for the remaining null SNPs. We then assumed *z_i,add_*’s (in black on top-left panel) as the additional information available, which were drawn, independently, from *N*(*μ_add_*, 1) for *m_add_* SNPs and *N*(0, 1) for the remaining SNPs.

**Figure S18:**
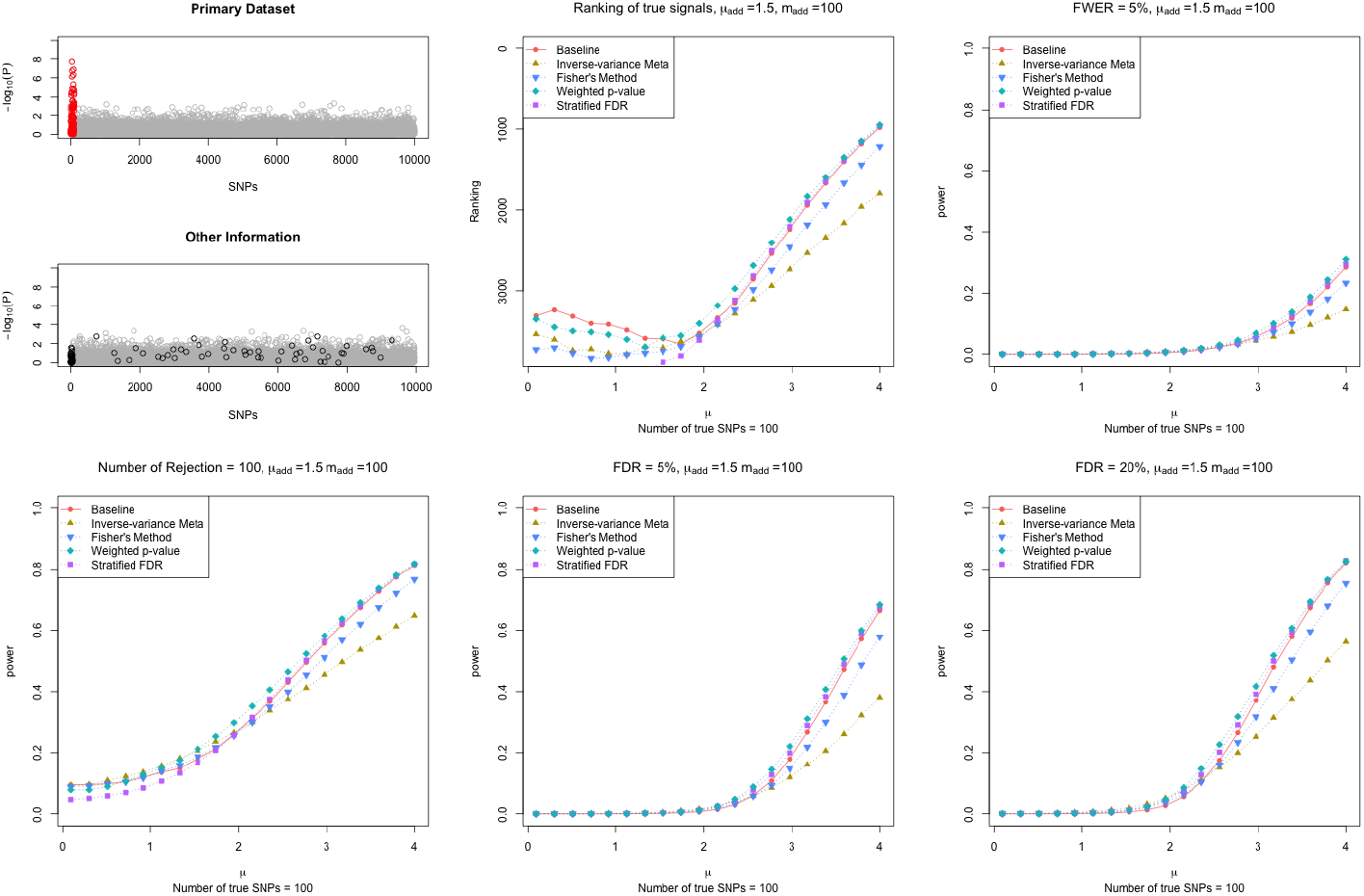
Results of simulation study design III for scenario (6) in Figure S10 from Category III type of informativeness, with the true effect size *μ*_1_ for the *m*_1_ causal SNPs varying from 0.1 to 4. Partially Misleading: *m_add_* = 100, *μ_add_* = 1.5 but only 50 out of the *m_add_* SNPs coincide with 50 of the *m*_1_ SNPs. We assumed the total number of SNPs *m* = 10, 000, among which the first *m*_1_ = 100 SNPs (in red on top-left panel) are truly associated. The corresponding summary statistics zi’s were drawn, independently, from *N*(*μ*_1_,1) for the *m*_1_ associated SNPs, and from *N*(0, 1)for the remaining null SNPs. We then assumed *z_i,add_*’s (in black on top-left panel) as the additional information available, which were drawn, independently, from *N*(*μ_add_*, 1) for *m_add_* SNPs and *N*(0, 1) for the remaining SNPs.

**Figure S19:**
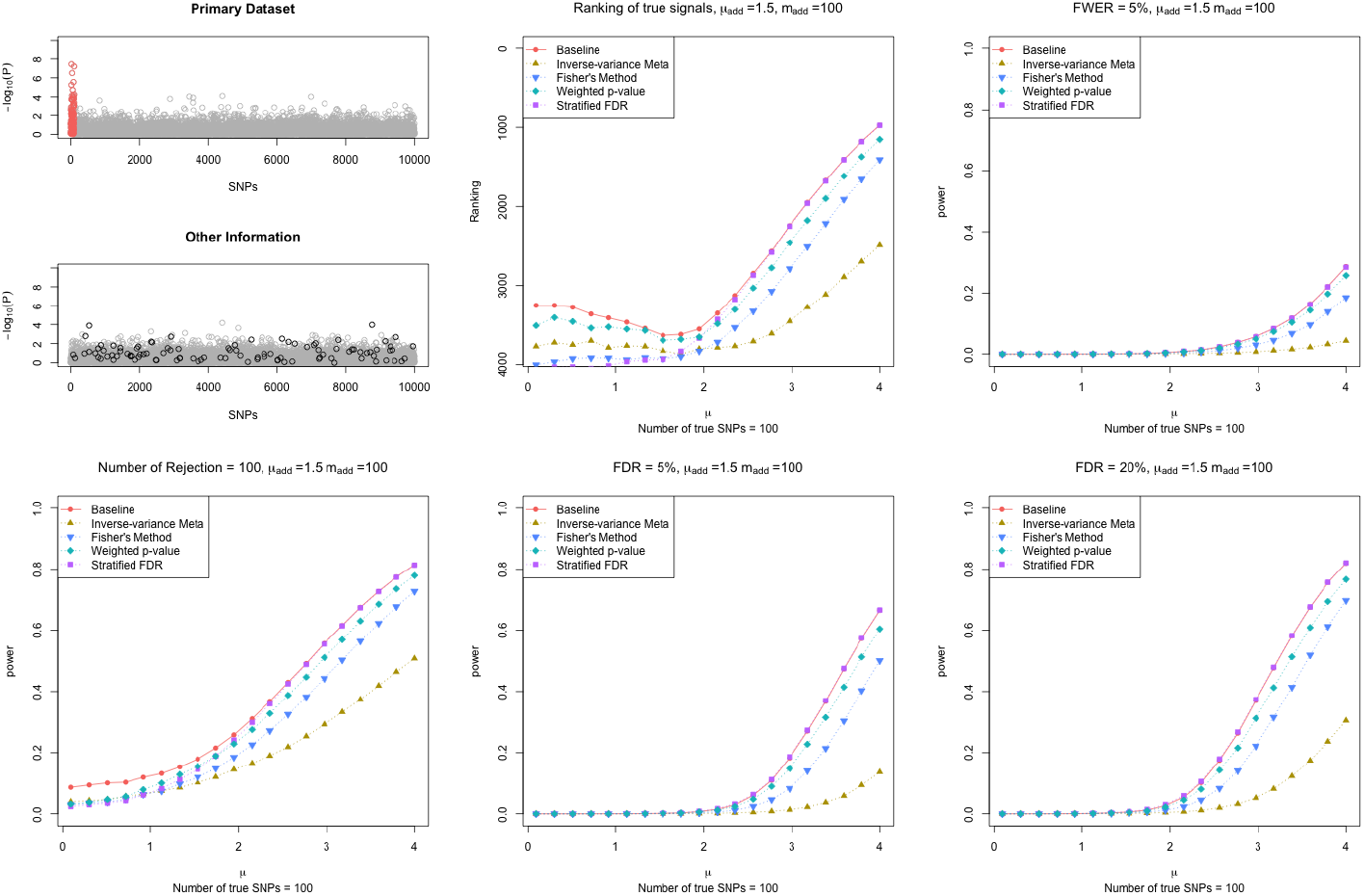
Results of simulation study design III for scenario (7) in Figure S10 from Category III type of informativeness, with the true effect size *μ*_1_ for the *m*_1_ causal SNPs varying from 0.1 to 4. Completely Misleading: *m_add_* = 100 and *μ_add_* = 1.5, but none of the *m_add_* SNPs coincide with the *m*_1_ SNPs. We assumed the total number of SNPs *m* = 10, 000, among which the first *m*_1_ = 100 SNPs (in red on top-left panel) are truly associated. The corresponding summary statistics zi’s were drawn, independently, from *N*(*μ*_1_,1) for the *m*_1_ associated SNPs, and from *N*(0, 1) for the remaining null SNPs. We then assumed *z_i,add_*’s (in black on top-left panel) as the additional information available, which were drawn, independently, from *N*(*μ_add_*, 1) for *m_add_* SNPs and *N*(0, 1) for the remaining SNPs.

**Figure S20:**
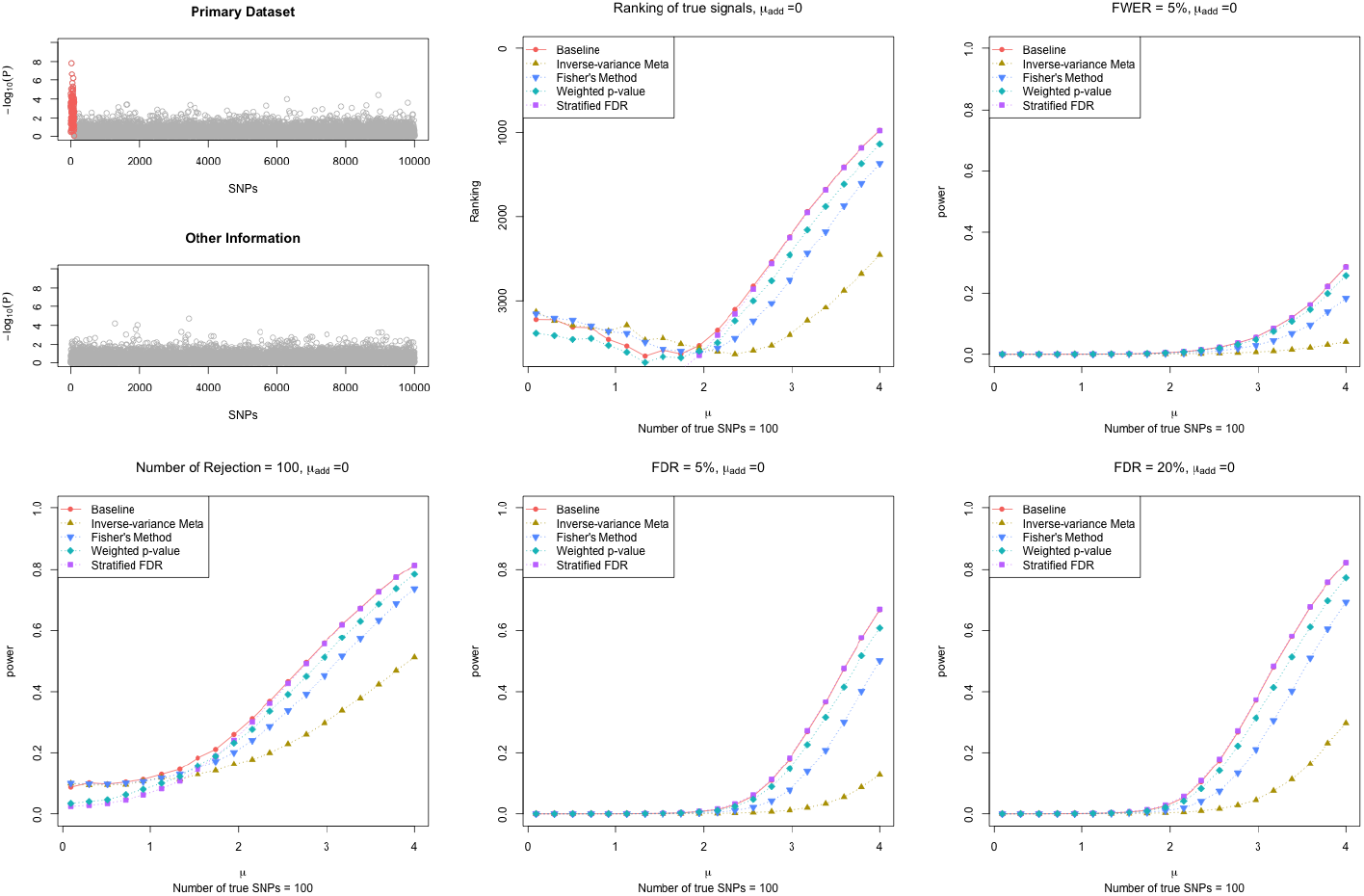
Results of simulation study design III for scenario (8) in Figure S10 from Category I type of informativeness, with the true effect size *μ*_1_ for the *m*_1_ causal SNPs varying from 0.1 to 4. Uninformative: *m_add_* = 0 and *μ_add_* = 0. That is, the additional information available is white noise. We assumed the total number of SNPs *m* = 10, 000, among which the first *m*_1_ = 100 SNPs (in red on top-left panel) are truly associated. The corresponding summary statistics *z_i_*’s were drawn, independently, from *N*(*μ*_1_,1) for the *m*_1_ associated SNPs, and from *N*(0, 1) for the remaining null SNPs. We then assumed *z_i,add_*’s (in black on top-left panel) as the additional information available, which were drawn, independently, from *N*(*μ_add_*, 1) for *m_add_* SNPs and *N*(0, 1) for the remaining SNPs.

**Figure S21:**
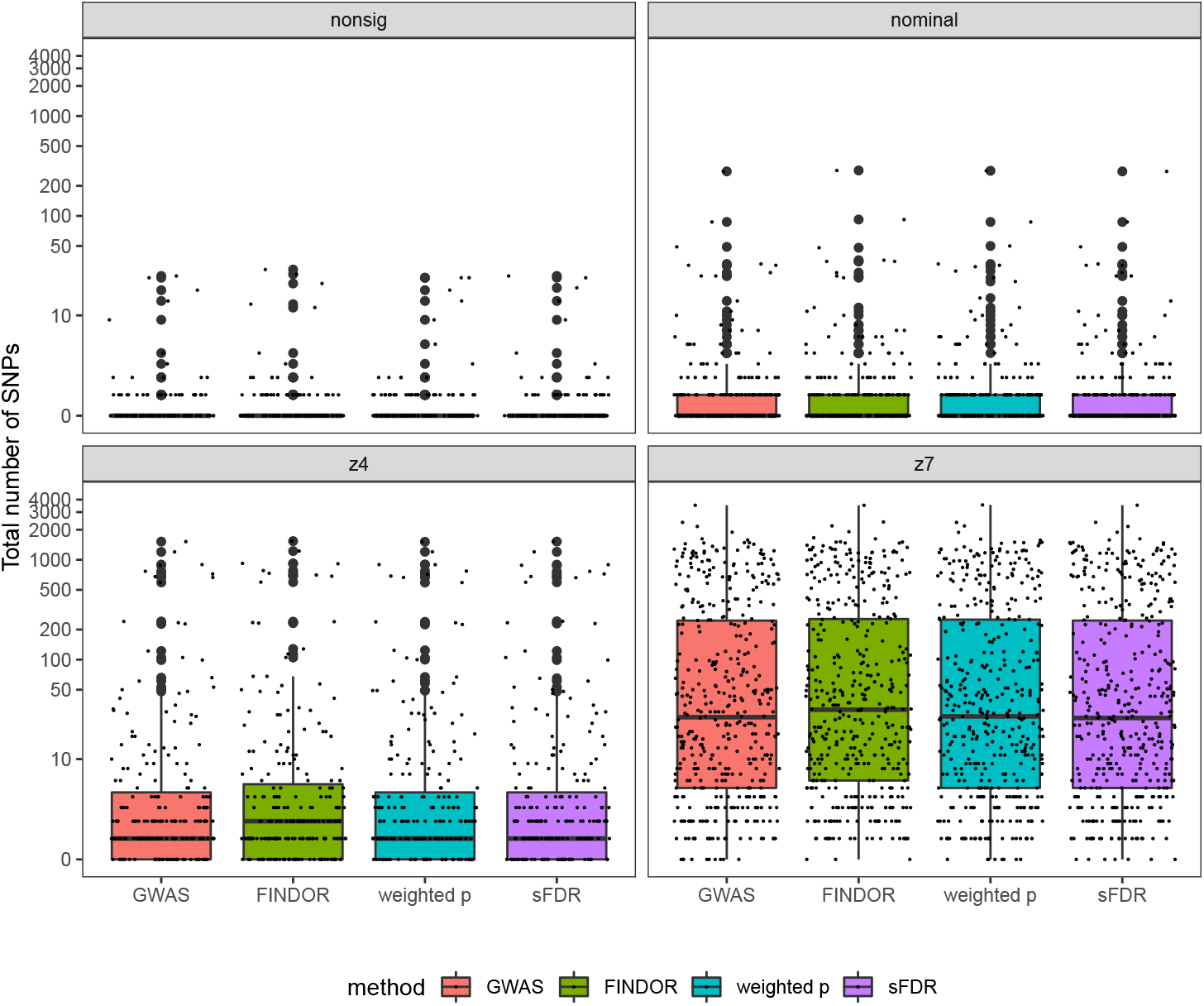
Results of using Eigen meta-scores for the total numbers of genome-wide significant independent loci of the UK Biobank GWAS application study, before and after data-integration with functional annotations, stratified by the four phenotype categories. In each figure, the total number of significant loci identified based on the UK Biobank GWAS data alone serves as a baseline. The GWAS baseline box-plot is followed by the box-plots for the total numbers of significant loci after integrating the UK Biobank GWAS summary statistics with functional annotations using FINDOR (using 75 individual annotation scores), and the weighted p-value and stratified FDR control methods (each using the Eigen meta-score), analyzing 7,895,174 variants for each of the 1,132 UK Biobank traits. The 1,132 traits were rated by Nealelab as having medium to high confidence for their heritability estimates, and they fall into four categories: nonsig (182 traits; heritability testing p-value *p* > 0.05), nominal (277 traits; *p* < 0.05), z4 (235 traits; *p* < 3.17 × 10^−5^), and z7 (438 traits; *p* < 1.28 × 10^−12^). Independent loci were defined using PLINK’s LDclumping algorithm with a 1 Mb window and an *r*^2^ threshold of 0.1.

**Figure S22:**
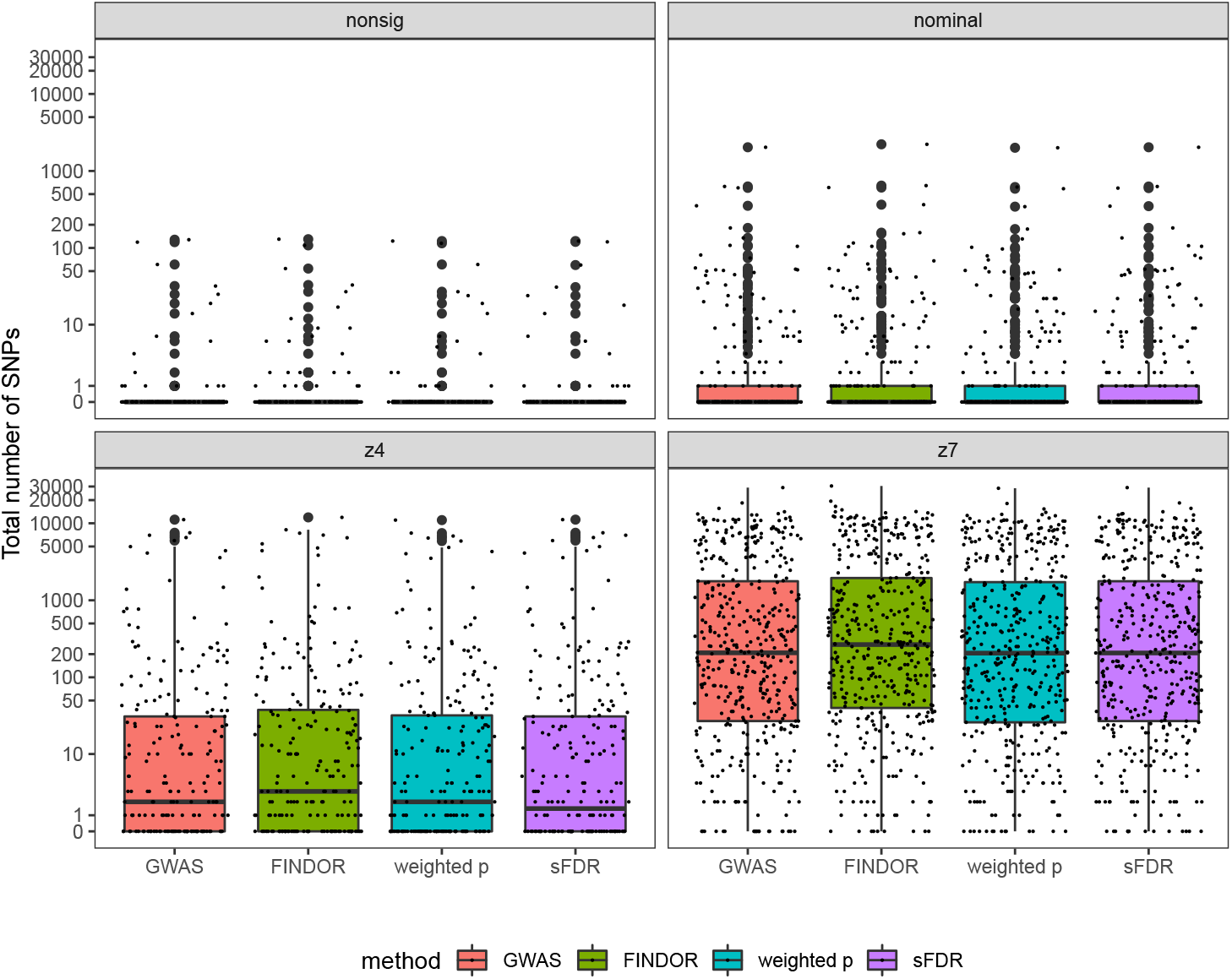
Results of counting significant SNPs instead of loci for the total numbers of genome-wide significant SNPs of the UK Biobank GWAS application study, before and after data-integration with functional annotations, stratified by the four phenotype categories. In each figure, the total number of significant loci identified based on the UK Biobank GWAS data alone serves as a baseline. The GWAS baseline box-plot is followed by the box-plots for the total numbers of significant loci after integrating the UK Biobank GWAS summary statistics with functional annotations using FINDOR (using 75 individual annotation scores), and the weighted p-value and stratified FDR control methods (each using the CADD meta-score), analyzing 7,895,174 variants for each of the 1,132 UK Biobank traits. The 1,132 traits were rated by Nealelab as having medium to high confidence for their heritability estimates, and they fall into four categories: nonsig (182 traits; heritability testing p-value *p* > 0.05), nominal (277 traits; *p* < 0.05), z4 (235 traits; *p* < 3.17 × 10^−5^), and z7 (438 traits; *p* < 1.28 × 10^−12^). Independent loci were defined using PLINK’s LDclumping algorithm with a 1 Mb window and an *r*^2^ threshold of 0.1.

**Figure S23:**
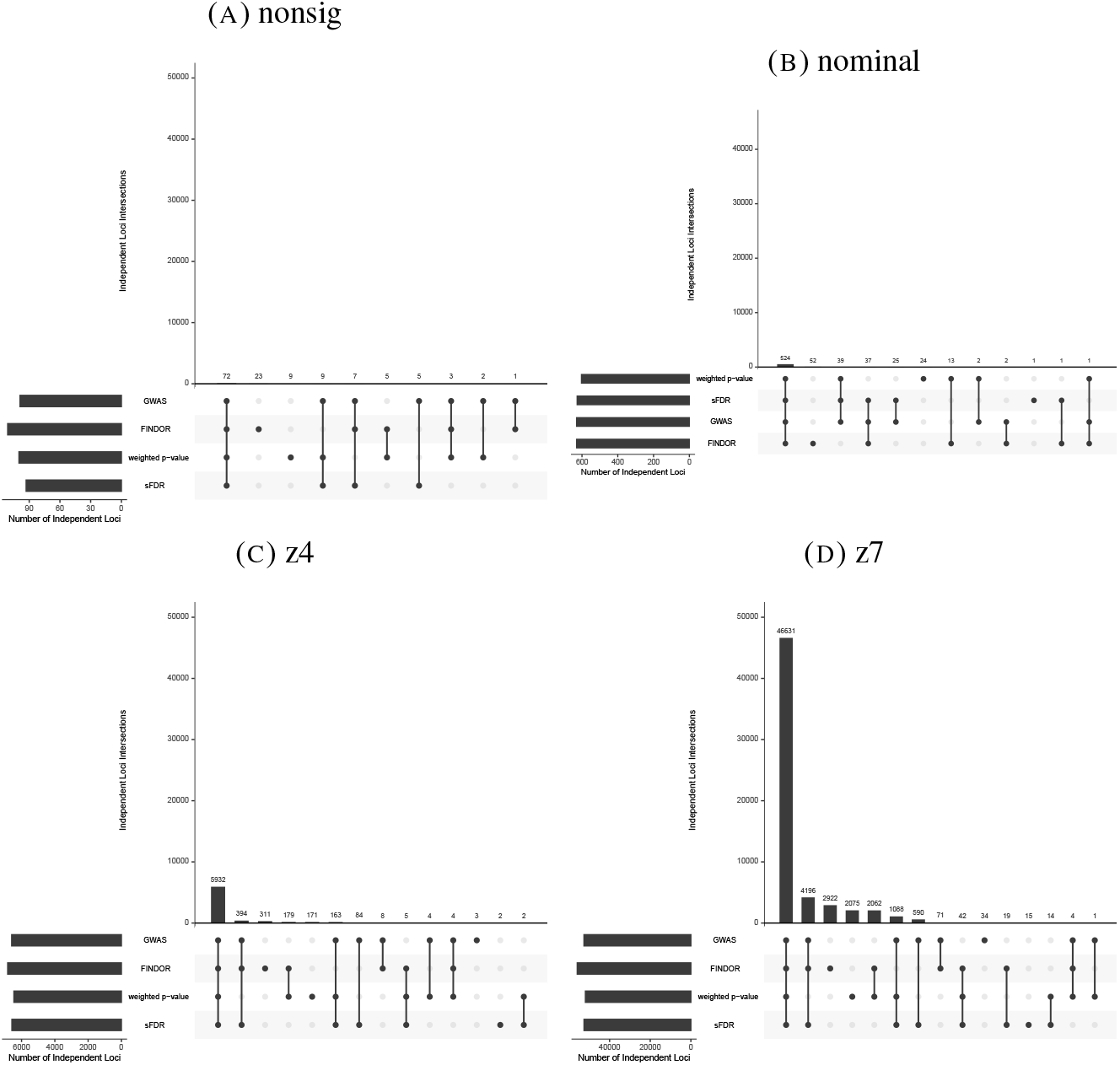
Intersection of significant, independent loci of the UK Biobank GWAS application study, before and after data-integration with functional annotations, stratified by the four phenotype categories. The three data-integration methods integrated the UK Biobank GWAS summary statistics with functional annotations using FINDOR (using 75 individual annotation scores), and the weighted p-value andstratified FDR control methods (each using the CADD meta-score), analyzing 7,895,174 variants for each of the 1,132 UK Biobank traits. The 1,132 traits were rated by Nealelabas having medium to high confidence for their heritability estimates, and they fall intofour categories: nonsig (182 traits; heritability testing *p* > 0.05), nominal (277 traits; *p* < 0.05), z4 (235 traits; *p* < 3.17 × 10^−5^), and z7 (438 traits; *p* < 1.28 × 10^−12^). Independent loci were defined using PLINK’s LDclumping algorithm with a 1 Mb window and an *r*^2^ threshold of 0.1.

**Figure S24:**
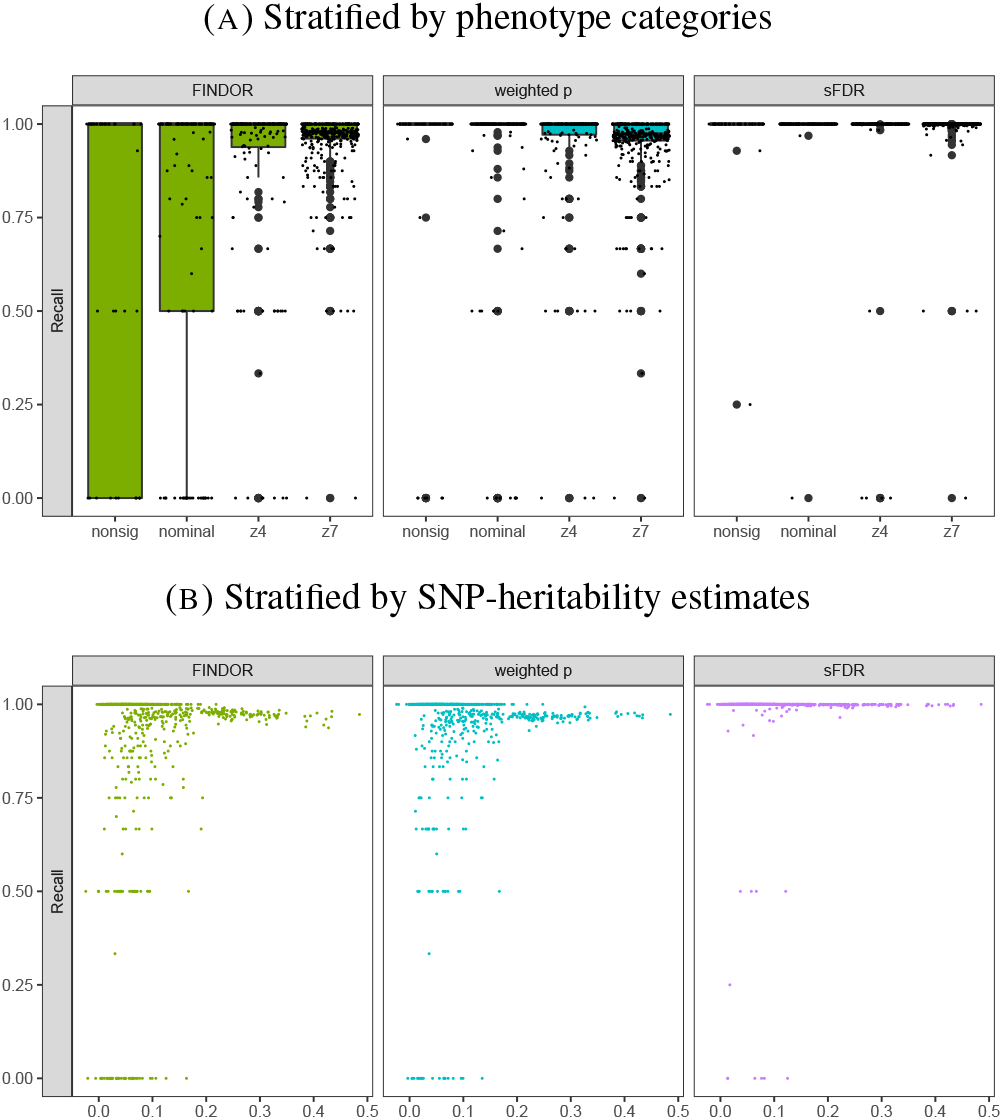
*Recall* results of the UK Biobank GWAS application study stratified by (A) the four phenotype categories (B) SNP-heritability estimates, for all 722 traits with *m*_1,*t*>0_. *Recall_t_* = *TP_t_*/*m*_1,*t*_, where *m*_1,*t*_ is the number of genome-wide significant independent loci prior to data-integration for trait *t*, and *TP_t_* is the number of true positives after data-integration. The three data-integration methods integrated the UK Biobank GWAS summary statistics with functional annotations using FINDOR (using 75 individual annotation scores), and the weighted p-value andstratified FDR control methods (each using the CADD meta-score), analyzing 7,895,174 variants for each of the 1,132 UK Biobank traits. The 1,132 traits were rated by Nealelabas having medium to high confidence for their heritability estimates, and they fall intofour categories: nonsig (182 traits; heritability testing *p* > 0.05), nominal (277 traits; *p* < 0.05), z4 (235 traits; *p* < 3.17 × 10^−5^), and z7 (438 traits; *p* < 1.28 × 10^−12^). Independent loci were defined using PLINK’s LDclumping algorithm with a 1 Mb window and an *r*^2^ threshold of 0.1.

**Figure S25:**
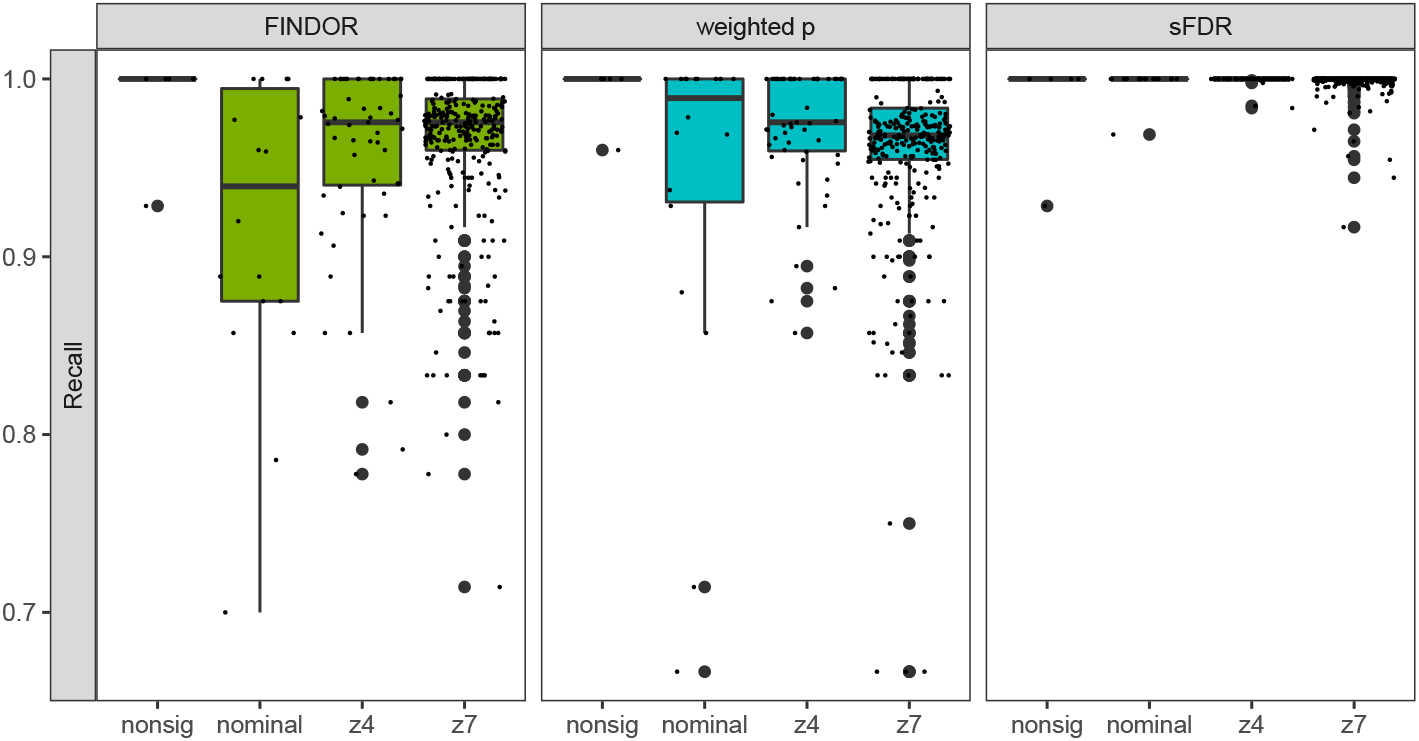
*Recall* results of the UK Biobank GWAS application study stratified by the four phenotype categories, for 402 traits with *m*_1,*t*>5_. *Recall_t_* = *TP_t_*/*m*_1,*t*_, where *m*_1,*t*_ is the number of genome-wide significant independent loci prior to data-integration for trait *t*, and *TP_t_* is the number of true positives after data-integration. The three data-integration methods integrated the UK Biobank GWAS summary statistics with functional annotations using FINDOR (using 75 individual annotation scores), and the weighted p-value andstratified FDR control methods (each using the CADD meta-score), analyzing 7,895,174 variants for each of the 1,132 UK Biobank traits. The 1,132 traits were rated by Nealelabas having medium to high confidence for their heritability estimates, and they fall intofour categories: nonsig (182 traits; heritability testing *p* > 0.05), nominal (277 traits; *p* < 0.05), z4 (235 traits; *p* < 3.17 × 10^−5^), and z7 (438 traits; *p* < 1.28 × 10^−12^). Independent loci were defined using PLINK’s LDclumping algorithm with a 1 Mb window and an *r*^2^ threshold of 0.1.

**Figure S26:**
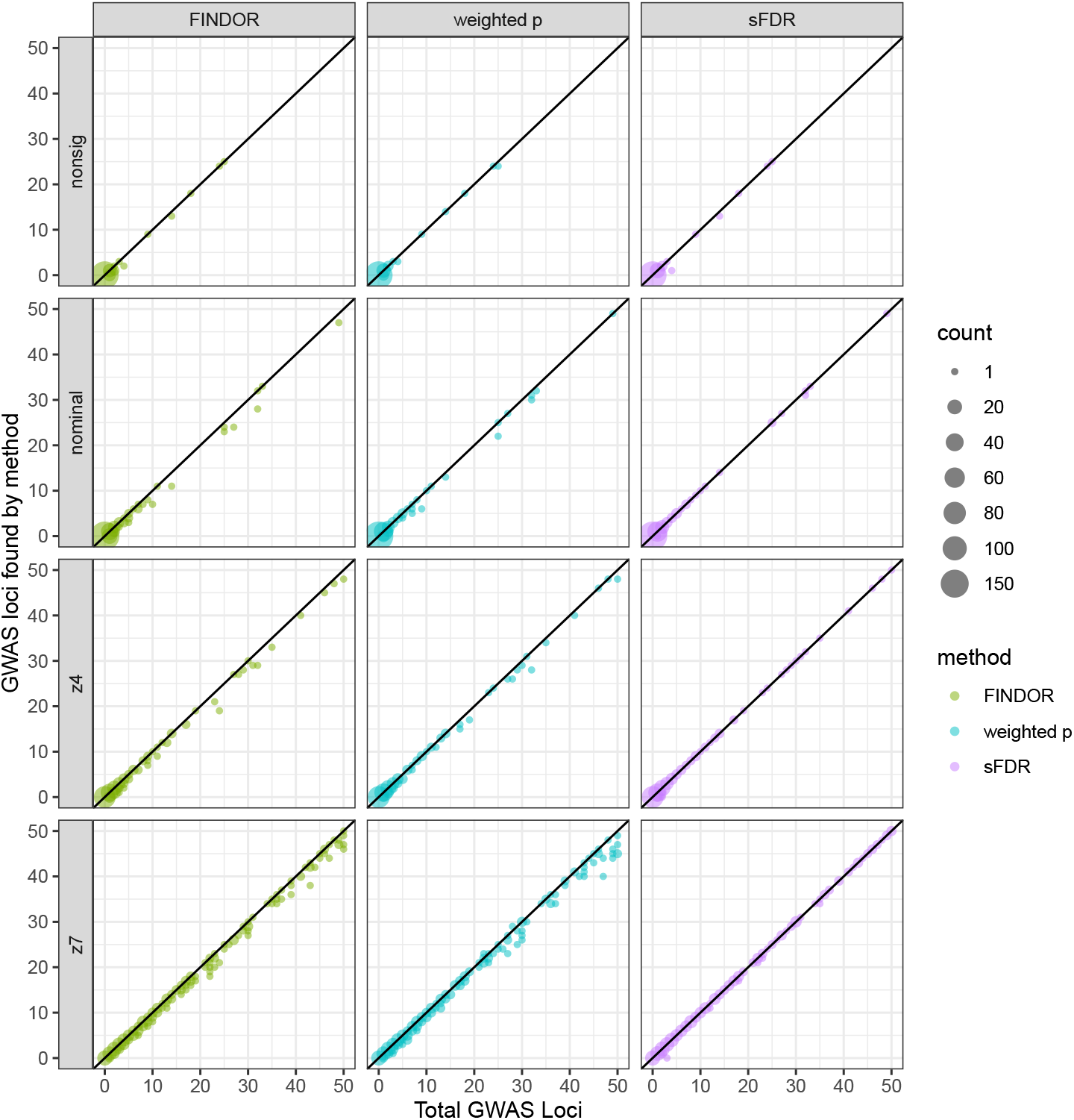
Contrast of the numbers of significant loci before and after data-integration for 942 traits with *m*_1,*t*_ ≤ 50. *m*_1,*t*_ is the number of genome-wide significant independent loci prior to data-integration for trait *t*, The three data-integration methods integrated the UK Biobank GWAS summary statistics with functional annotations using FINDOR (using 75 individual annotation scores), and the weighted p-value and stratified FDR control methods (each using the CADD meta-score), analyzing 7,895,174 variants for each of the 1,132 UK Biobank traits. The 1,132 traits were rated by Nealelabas having medium to high confidence for their heritability estimates, and they fall intofour categories: nonsig (182 traits; heritability testing *p* > 0.05), nominal (277 traits; *p* < 0.05), z4 (235 traits; *p* < 3.17 × 10^−5^), and z7 (438 traits; *p* < 1.28 × 10^−12^). Independent loci were defined using PLINK’s LDclumping algorithm with a 1 Mb window and an *r*^2^ threshold of 0.1.

**Figure S27:**
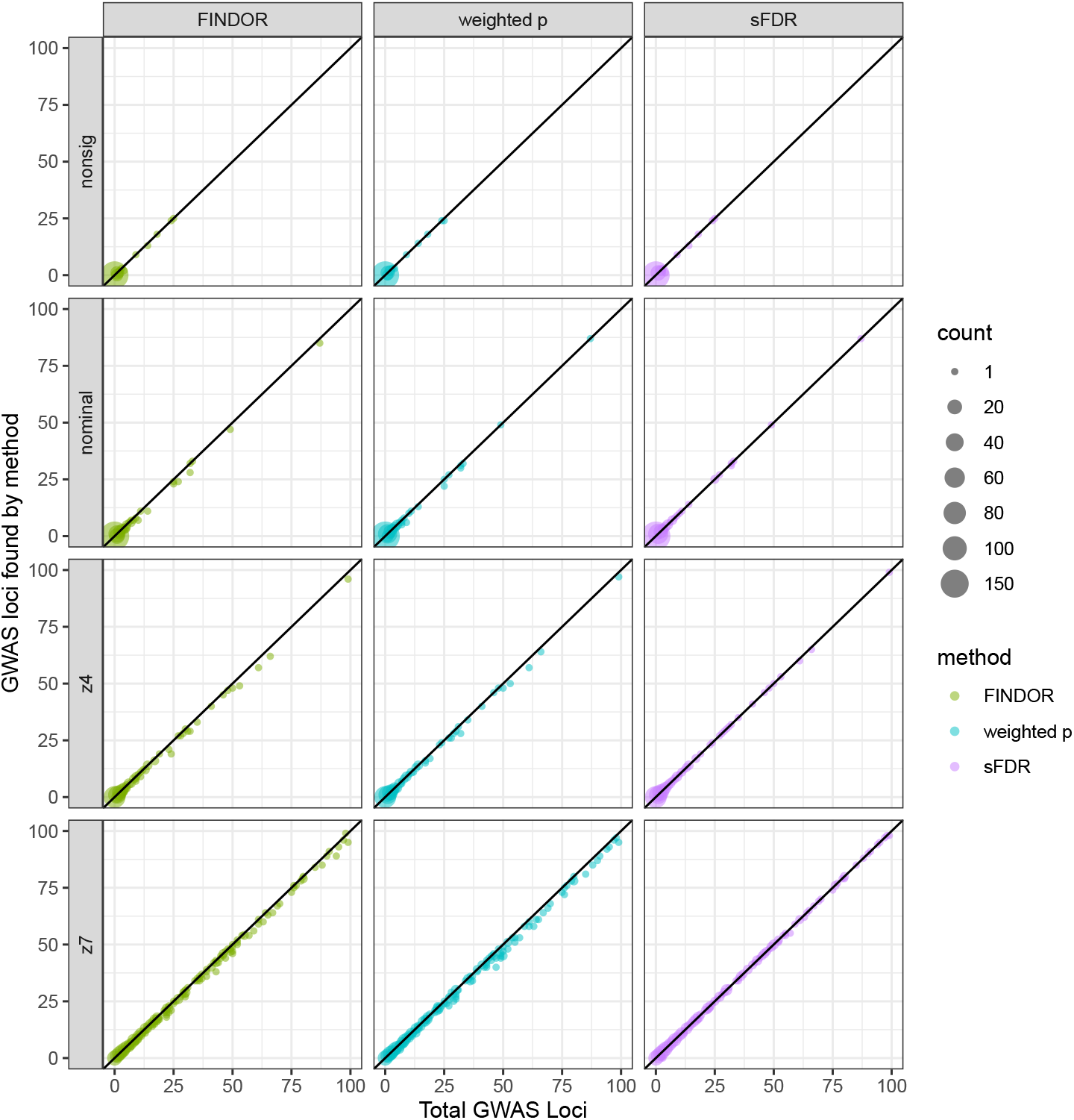
Contrast of the numbers of significant loci before and after data-integration for 980 traits with *m*_1,*t*_ ≤ 100. *m*_1,*t*_ is the number of genome-wide significant independent loci prior to data-integration for trait *t*, The three data-integration methods integrated the UK Biobank GWAS summary statistics with functional annotations using FINDOR (using 75 individual annotation scores), and the weighted p-value and stratified FDR control methods (each using the CADD meta-score), analyzing 7,895,174 variants for each of the 1,132 UK Biobank traits. The 1,132 traits were rated by Nealelabas having medium to high confidence for their heritability estimates, and they fall intofour categories: nonsig (182 traits; heritability testing *p* > 0.05), nominal (277 traits; *p* < 0.05), z4 (235 traits; *p* < 3.17 × 10^−5^), and z7 (438 traits; *p* < 1.28 × 10^−12^). Independent loci were defined using PLINK’s LDclumping algorithm with a 1 Mb window and an *r*^2^ threshold of 0.1.

**Figure S28:**
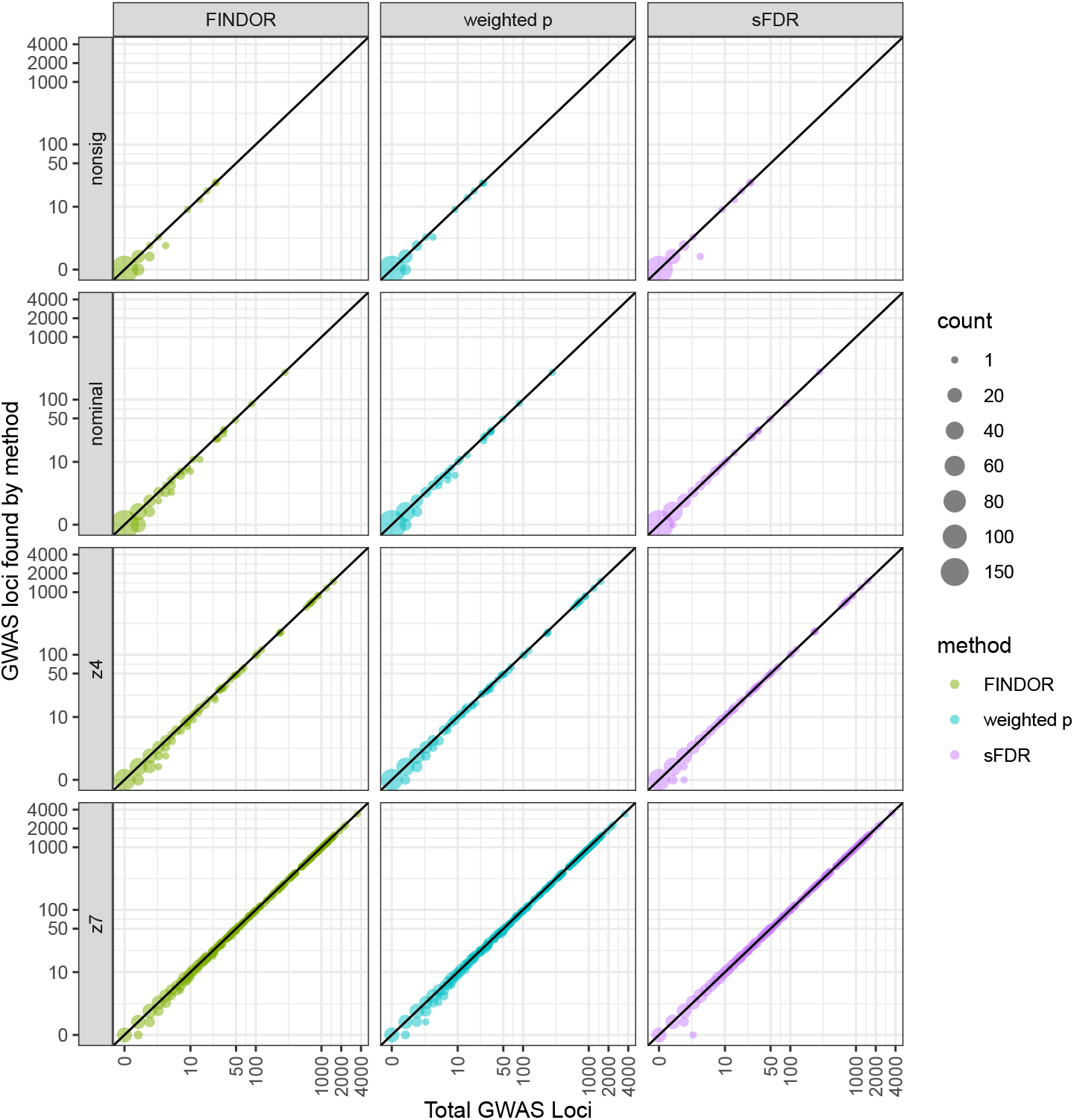
Contrast of the numbers of significant loci before and after data-integration for all 1,132 traits, i.e. *m*_1,*t*_ ≥ 0. *m*_1,*t*_ is the number of genome-wide significant independent loci prior to data-integration for trait *t*, The three data-integration methods integrated the UK Biobank GWAS summary statistics with functional annotations using FINDOR (using 75 individual annotation scores), and the weighted p-value and stratified FDR control methods (each using the CADD meta-score), analyzing 7,895,174 variants for each of the 1,132 UK Biobank traits. The 1,132 traits were rated by Nealelabas having medium to high confidence for their heritability estimates, and they fall intofour categories: nonsig (182 traits; heritability testing *p* > 0.05), nominal (277 traits; *p* < 0.05), z4 (235 traits; *p* < 3.17 × 10^−5^), and z7 (438 traits; *p* < 1.28 × 10^−12^). Independent loci were defined using PLINK’s LDclumping algorithm with a 1 Mb window and an *r*^2^ threshold of 0.1.

**Figure S29:**
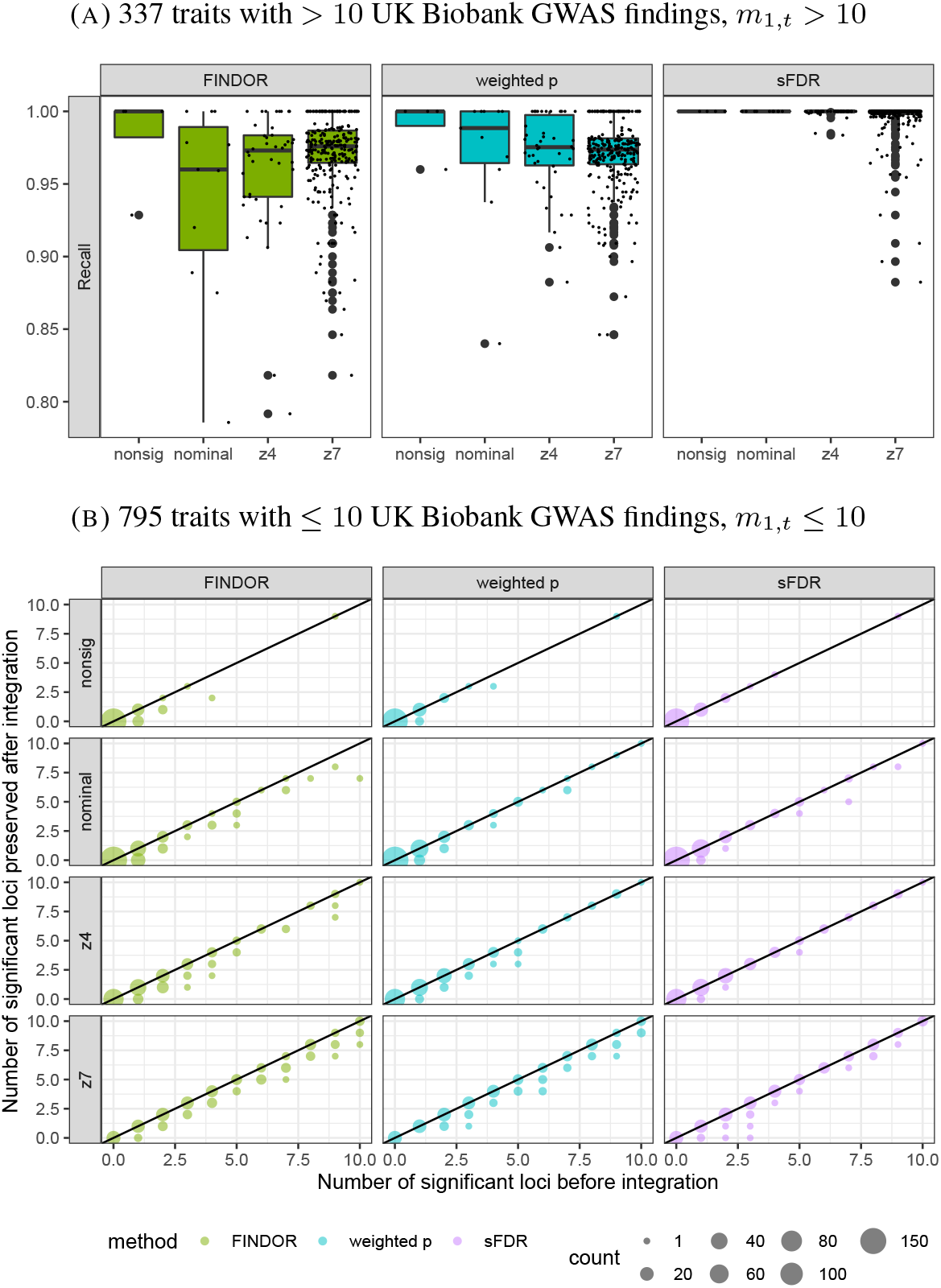
Eigen results of the UK Biobank GWAS application study, before and after data-integration with functional annotations, stratified by the four phenotype categories. (A) *Recall_t_* = *TP_t_*/*m*_1,*t*_, where *m*_1,*t*_ is the number of genome-wide significant independent loci prior to data-integration for trait *t*, and *TP_t_* is the number of true positives after data-integration. *Recall* estimation is not stable when *m*_1,*t*_ is small so for *m*_1,*t*_ ≤ 10, (B) contrasts the number of significant loci preserved after data-integration with *m*_1,*t*_. The three data-integration methods integrated the UK Biobank GWAS summary statistics with functional annotations using FINDOR (using 75 individual annotation scores), and the weighted p-value andstratified FDR control methods (each using the Eigen meta-score), analyzing 7,895,174 variants for each of the 1,132 UK Biobank traits. The 1,132 traits were rated by Nealelabas having medium to high confidence for their heritability estimates, and they fall intofour categories: nonsig (182 traits; heritability testing *p* > 0.05), nominal (277 traits; *p* < 0.05), z4 (235 traits; *p* < 3.17 × 10^−5^), and z7 (438 traits; *p* < 1.28 × 10^−12^). Independent loci were defined using PLINK’s LDclumping algorithm with a 1 Mb window and an *r*^2^ threshold of 0.1.

**Figure S30:**
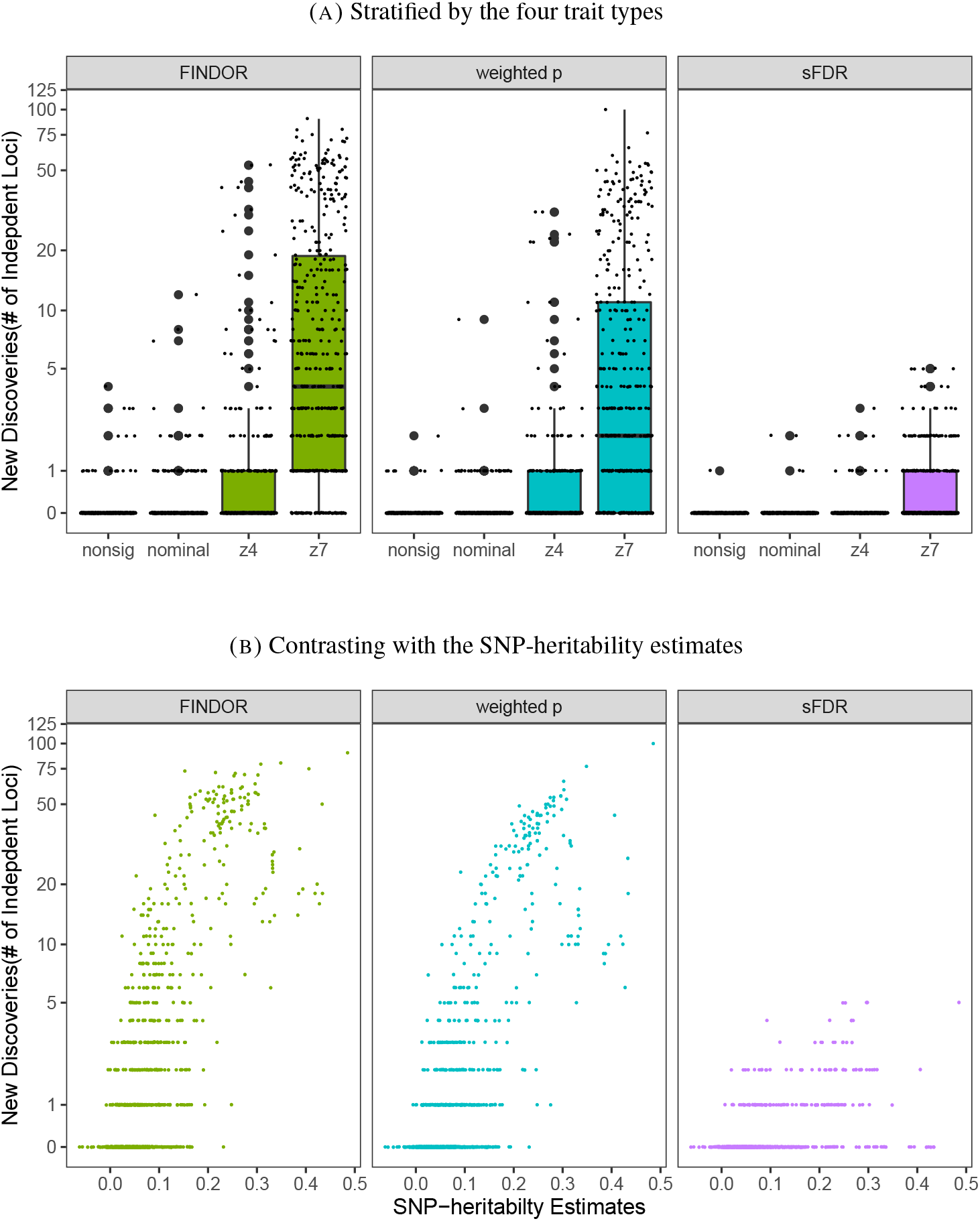
The Eigen results of the total *New Discoveries* of the application study (A) stratified by the four trait types or (B) Contrasting with the SNP-heritability estimates. The three data-integration methods integrated the UK Biobank GWAS summary statistics with functional annotations using FINDOR (using 75 individual annotation scores), and the weighted p-value andstratified FDR control methods (each using the Eigen meta-score), analyzing 7,895,174 variants for each of the 1,132 UK Biobank traits. The 1,132 traits were rated by Nealelabas having medium to high confidence for their heritability estimates, and they fall intofour categories: nonsig (182 traits; heritability testing *p* > 0.05), nominal (277 traits; *p* < 0.05), z4 (235 traits; *p* < 3.17 × 10^−5^), and z7 (438 traits; *p* < 1.28 × 10^−12^). Independent loci were defined using PLINK’s LDclumping algorithm with a 1 Mb window and an *r*^2^ threshold of 0.1.

**Figure S31:**
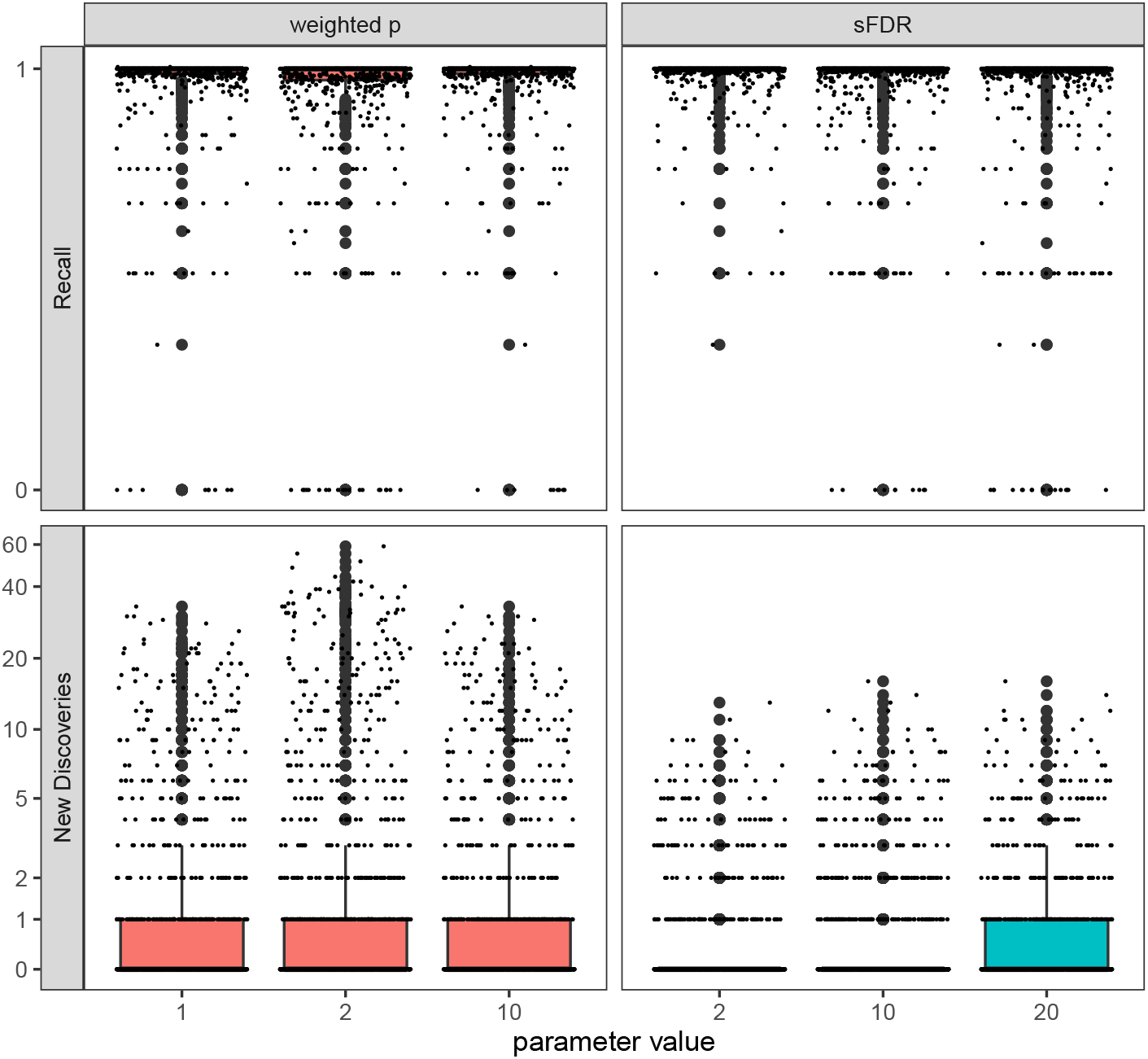
Performance of weighted p-value and sFDR methods using different weighting schemes and stratification for the UK Biobank GWAS application study. The two data-integration methods integrated the UKBiobank GWAS summary statistics with CADD meta-score, analyzing 7,895,174 variants for each of the 1,132 UK Biobank traits. Independent loci were defined using PLINK’s LDclumping algorithm with a 1 Mb window and an *r*^2^ threshold of 0.1. For the weighted p-value approach, the parameter values represent *β* = 1, 2 and 10 for the cumulative weighting scheme, 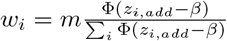, where *z_i,add_* is the CADD meta-score for variant *i*. For the sFDR approach, the parameter values 2, 10 and 20 represent the number of strata used, where the stratification is based on quantiles of *z_i,add_*, the CADD meta-score for variant *i*.

